# Oxygen-independent chemogenetic protein tags for live-cell fluorescence microscopy

**DOI:** 10.1101/2020.08.03.234815

**Authors:** Aditya Iyer, Maxim Baranov, Alexander J Foster, Shreyans Chordia, Gerard Roelfes, Geert van den Bogaart, Bert Poolman

## Abstract

Fluorescent proteins enable targeted visualization of biomolecules in living cells, but their maturation is oxygen-dependent and they are susceptible to aggregation and/or suffer from poor photophysical properties. Organic fluorophores are oxygen-independent with superior photophysical properties, but targeting biomolecules in vivo is challenging. Here, we introduce two oxygen-independent chemogenetic protein (OICP) tags that impart fluorogenicity and fluorescence lifetime enhancement to bound organic dyes. We present a photo- and physicochemical characterization of thirty fluorophores interacting with two OICPs and conclude that aromatic planar structures bind with high specificity to the hydrophobic pockets of the proteins. The binding specificity of the tags and the superior photophysical properties of organic fluorophores enable microscopy of living bacterial and eukaryotic cells. The exchange of photobleached dye for unbleached fluorophore enables prolonged live-cell imaging. Our protein tags provide a general tool for investigating (sub)cellular protein localization and dynamics, protein-protein interactions, and microscopy applications under strictly oxygen-free conditions.

## INTRODUCTION

Biochemistry is evolving from mostly in vitro studies of macromolecules to in vivo analyses of complex processes in living cells, wherein macromolecules and multiprotein complexes are mapped three-dimensionally with high spatial and temporal resolution and full functionality. To attain this, fluorescence live-cell imaging techniques have traditionally relied on tagging specific proteins in their native cellular environment with genetically encoded fluorescent proteins (FPs)(Rodriguez et al., 2016; Shaner et al., 2005). FPs are target-specific but often fall short in photophysical characteristics when benchmarked against organic fluorophores. Additionally, the anaerobic conditions of the gut and other environments pose a significant barrier for FP-based fluorescence imaging for investigating the physiology of endogenous microorganisms(Tropini et al., 2017). This is because FPs require molecular oxygen for chromophore maturation(Tsien, 1998) with the exception of flavin mononucleotide-dependent fluorescent protein((Drepper et al., 2007)) and the recently discovered bilirubin-dependent UnaG protein(Kumagai et al., 2013), and this requirement is a major obstacle hindering the studies of anaerobic microorganisms or cellular compartments low in oxygen.

Alternative to FPs, modified organic fluorophores, and/or bait proteins exist that capture a specific ligand coupled to organic fluorophores. Organic fluorophores typically have better photophysical characteristics compared to FPs such as greater photostability, longer fluorescent lifetimes, higher quantum yields and a wider spectral range, and importantly do not require oxygen for fluorescence(Balleza et al., 2018; Cranfill et al., 2016; Grimm et al., 2016; Jing and Cornish, 2011; Kocaoglu and Carlson, 2016; Shaner et al., 2005). Organic fluorophores are notorious for non-specific interactions with cellular components and high background signals. Not surprisingly, strategies to make smaller FPs with improved photophysical characteristics(Dippel et al., 2020; Drepper et al., 2007; Nagai et al., 2002), developing organic fluorophores with enhanced specificity and fluorogenicity (enhanced fluorescence upon binding target)(Kowada et al., 2015; Song et al., 2014), and peptide-tags like SNAP-tag® and HaloTag® have gained traction(Grimm et al., 2017; Jing and Cornish, 2011; Kocaoglu and Carlson, 2016; Wang et al., 2020). The application of these important tools can be limited by high nonspecific staining(Bosch et al., 2014) and poor cell permeability of the dyes(Keppler et al., 2004). Recently, SNAP-tag® and HaloTag® substrates have been covalently coupled to 6-TAMRA derivatives (MaP probes) that displayed enhanced cell permeability and fluorogenicity(Wang et al., 2020). However, as of now such strategies require specific chemistry for ligand binding; and employ ligands that irreversibly and/or covalently bind to the modified fluorophore. Furthermore, synthesis of such modified fluorophores is nontrivial and their availability is limited. Whereas the covalent linkage of SNAP-tag® and HaloTag® substrates is advantageous in some experimental designs, it may not be always desirable as a non-covalent reversible binding of fluorophores can be preferable for long-term imaging as fluorophores can be exchanged for non-photobleached ones.

To address the aforementioned issues and alleviate the need to synthesize or modify commercially available fluorophores, we present a straightforward labeling strategy that achieves fluorogenicity, lifetime enhancement, and target recognition of non-covalently bound dyes in a variety of organisms. We introduce a self-labeling protein tagging system that combines the best of genetic tags and organic fluorophores developed through a chemogenetic approach. Our approach exploits the unique biochemical properties of two small, bacterial transcriptional factors, namely: resistance antibiotic multiple regulator(Yamasaki et al., 2013, 2019) (RamR) and lactococcal multidrug resistance repressor(Madoori et al., 2009; Roelfes, 2019; Takeuchi et al., 2014) (LmrR) that differ in sequence, molecular weight and structure (**Supplementary Fig. 1**). Both RamR (from the Gram-negative bacterium *Salmonella typhimurium*(Yamasaki et al., 2019)) and LmrR (from the Gram-positive bacterium *Lactococcus lactis*(Agustiandari et al., 2008)) are homo-dimeric proteins that contain hydrophobic pocket(s) where planar organic compounds bind non-covalently with high specificity. Under native conditions, RamR and LmrR act by repressing the synthesis of multidrug efflux pumps, and this effect is removed upon binding to organic compounds such as antibiotics. The hydrophobic pocket(s) are attractive scaffolds since they bind a variety of aromatic molecules.

**Figure 1.**
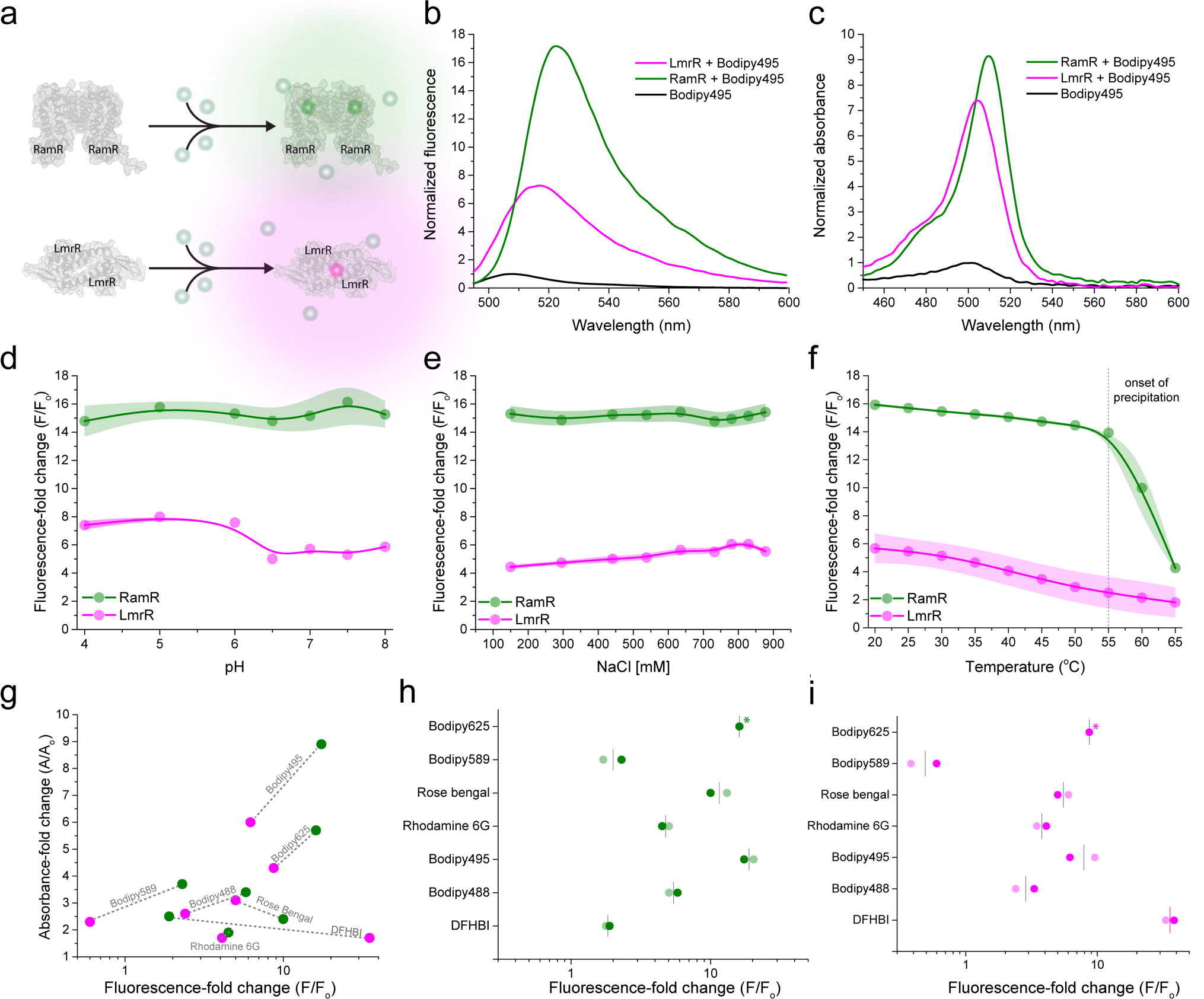
In vitro characterization of OICP tags. (a) Working principle of OICP tags: RamR (green) and LmrR (magenta). (b) Fluorescence emission spectra depicting fold-change in fluorescence emission intensity of Bodipy495 after adding of OICP tags. (c) Absorption spectra depicting fold-change in the absorption of Bodipy495 after the addition of OICP tags. (d-f) Fold-change in Bodipy495 fluorescence emission intensity across pH (d), NaCl concentration (e) and temperature (f). The dotted line in panel f indicates the onset of partial precipitation of RamR. The solid lines in panels d-f are spline fits and the color-shaded regions represent s.d. over three independent measurements. g) Correlation plots of fold-change in absorbance and fluorescence for 7 fluorogenic dyes. Grey dotted lines connect the values for the respective dye. (h-i) Effect of oxygen on the fluorogenicity of dyes in the presence of OICP tags. The fold change in fluorescence in the absence of oxygen (faded points) is comparable with that in the presence of oxygen (solid points). Fold-change in fluorescence of Bodipy625 in the absence of oxygen (indicated with asterisks) could not be measured due to the lack of the appropriate excitation source.

## RESULTS

We envisioned that seclusion of planar organic fluorophores into the hydrophobic binding pockets within LmrR(Mejías et al., 2020) and RamR (hereafter named oxygen-independent chemogenetic protein (OICP) tags) could improve their photophysical properties, which depends on the distinct chemical environment of the protein binding sites. Furthermore, we used engineered variants of LmrR to minimize unspecific interactions of the tags with cell components (see Methods section). Indeed, *in vitro* characterization of the spectroscopic properties of 30 organic fluorophores, several have applications in super-resolution microscopy and single-particle tracking, demonstrated that BODIPY, SNAP- and Halo-tag conjugated dyes and rhodamine-based fluorophores show a significant and correlated increase in fluorescence (up to 35-fold) with absorbance increase (up to 10-fold) in the presence of OICP tags, while that of others was either quenched or not affected (**Figure 1a-c, Table 1** and **Supplementary Figs. 2-18**). We verified the promiscuity of the hydrophobic dye-binding pockets using organic dyes that attain fluorogenicity in hydrophobic environments (**Table 1: non-specific intercalators**).

**Table 1.**
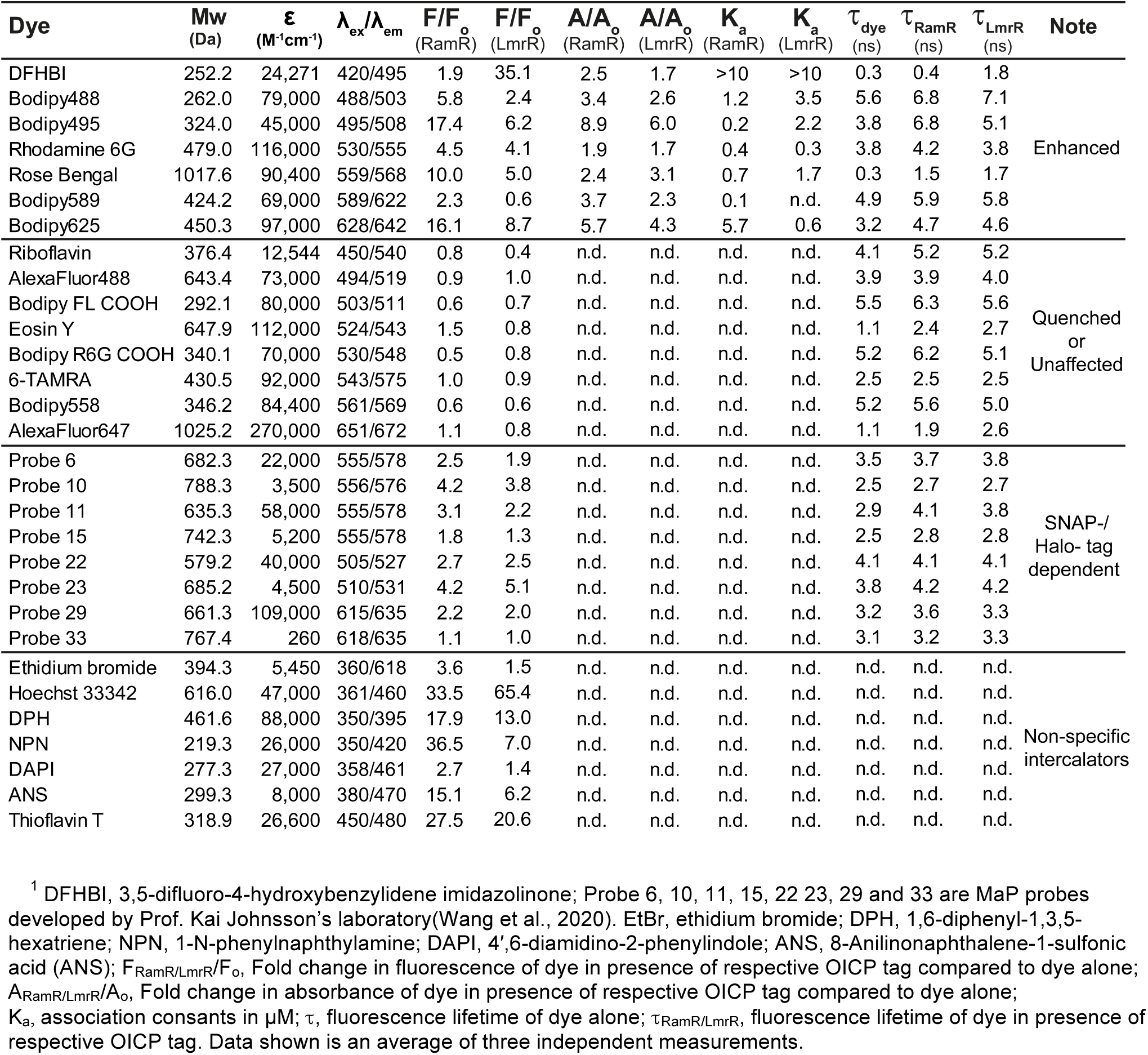
Spectral properties of tested fluorophores in the presence of OICP tags^1^.

**Figure 2.**
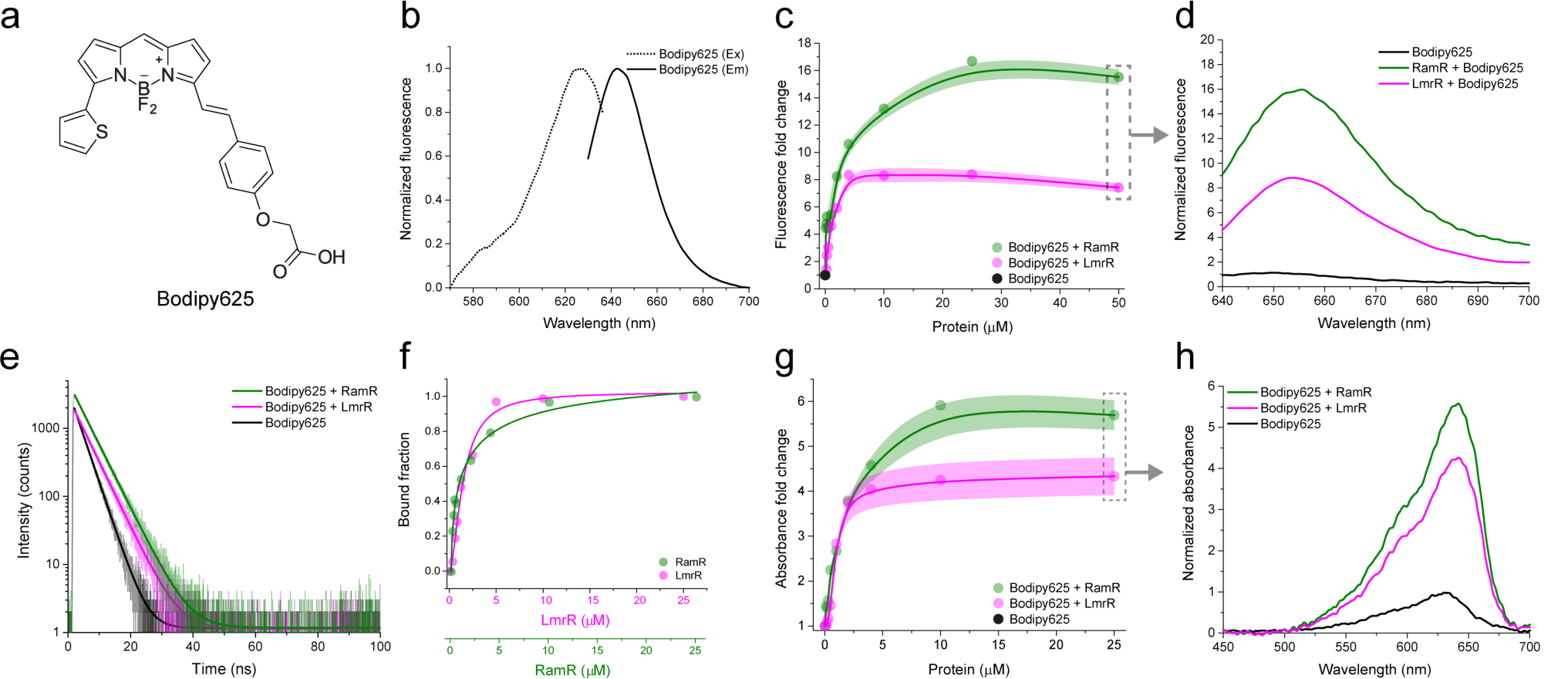
Representative characterization of OICP tags (RamR: green and LmrR: magenta) with Bodipy625. (a) Structure of Bodipy625. (b) Excitation (dotted line) and emission (solid line) spectra of Bodipy625. (c) Fluorescence fold change of Bodipy625 on titration with OICP tags. (d) Fluorescence emission spectra of Bodipy625 with OICP tags at a protein:dye molar ratio of 50:1. (e) Absorbance fold-change of Bodipy625 on titration with OICP tags. Solid lines represent spline fits and shaded regions represent s.d. over three independent measurements in panel (c) and (e). (f) Absorption spectra of Bodipy625 with OICP tags at a protein:dye molar ratio of 25:1. (g) Fluorescence lifetime spectra of Bodipy625 with OICP tags at a protein:dye molar ratio of 25:1 fit with a mono-exponential decay function (solid lines). (h) The bound fraction of Bodipy625 with OICP tags ascertained from a Hill fit. All experiments were performed at 30 °C in 20 mM K-MOPS, 150 mM NaCl buffered at pH 7.0.

We first characterized the physicochemical robustness of the purified OICP tags in detail *in vitro*. Using Bodipy495, the dye with the highest fluorogenicity amongst the seven fluorogenic dyes, we show that both OICP tags are insensitive to pH in the range from 4 to 8, salt concentrations up to 880 mM, temperatures up to 55 °C (**Figure 1d-f**) and crowding agent Ficoll70 (**Supplementary Fig. 19**), allowing applications in diverse environments. The fluorogenic behavior of the seven dyes is accompanied by corresponding increases in absorbance (**Figure 1g, Table 1, and Supplementary Fig 26)**. We also show that the dyes remain fluorescent in the absence and presence of oxygen (**Figure 1h-i**).

Next, we evaluated in-depth the spectral properties of 30 commonly available organic dyes with our OICP tags. A representative characterization of OICP tags with Bodipy625 is shown in **Figure 2**; the same characterization of 29 other dyes are shown in **Supplementary Figs. 2-18**. With most dyes (except DFHBI), the fluorescence enhancement, the enhanced absorbance, and lifetime increase are higher with RamR than LmrR. The pertinent data are summarized in Table 1.

Overall, our *in vitro* characterization of 30 organic dyes demonstrates the applicability of our tagging system in living cells. Our approach is easily extended to study the localization and dynamics of proteins in sub-compartments within living cells. To illustrate this, we tagged and labeled test proteins in the cytoplasm, inner membrane (penicillin-binding protein 5; PBP5), and periplasm (osmotically inducible protein Y; OsmY) of *E. coli* (**Figure 3a**). We chose Bodipy495 and Bodipy625 for these measurements since they show the highest fluorogenic behavior (**Table 1**), negligible background staining, and high cell permeabilities compared to other enhanced dyes (**Supplementary Fig. 20-23**). For cytoplasmic staining, we observe some non-specific binding of Bodipy495, likely to the inner membrane (**Figure 3a: Cytoplasm**), which is absent in controls (**Supplementary Fig. 20**). We confirmed specific targeting and labeling of the test proteins by performing fluorescence recovery after photobleaching measurements to compare the diffusion coefficients of test protein-OICP tag-dye conjugates with the same test proteins conjugated to a fluorescent protein, SuperFolder mTurquiose2ox(SfTq2) (**Figure 3b**). We find that the diffusion coefficients of the test proteins with dye conjugated-OICP and SfTq2 conjugated-OICP tags are comparable, with no indications for higher oligomer formation or protein aggregation.

**Figure 3.**
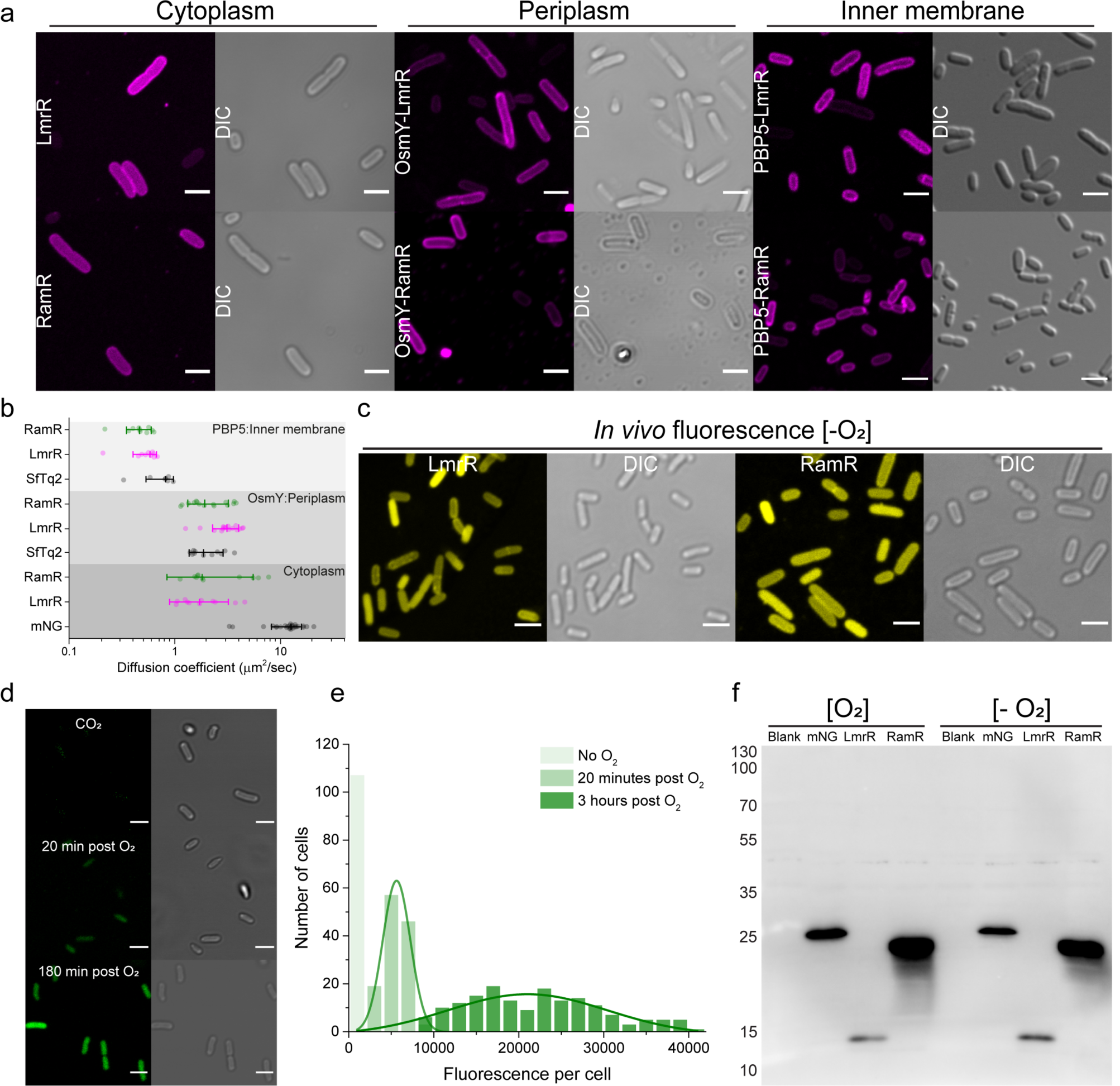
Compartmental labeling and oxygen-free imaging of OICP tags in *E. coli*. (a) Confocal and accompanying differential interference contrast (DIC) micrographs of live *E. coli* cells expressing OICP-tagged cytoplasmic (LmrR or RamR only), periplasmic (OsmY) and inner membrane protein (PBP5) labeled with Bodipy495. (b) Diffusion coefficients of the aforementioned cytoplasmic, periplasmic, and inner membrane proteins compared with independent control proteins: fluorescent SuperFolder mTurquiose2ox(SfTq2) and mNeongreen (mNG) from FRAP measurements. (c-e) Confocal and accompanying differential interference contrast (DIC) micrographs of live *E. coli* cells expressing cytoplasmic OICP tags labeled with Bodipy625 under strictly anaerobic conditions (c), expressing cytoplasmic mNG protein (d); the integrated fluorescence histograms of the panel (d) are shown (e). (f) Western blots of OICP tags (15 and 23 kDa for LmrR and RamR, respectively) and mNG protein (26.6 kDa) expressed in the *E. coli* cytoplasm under aerobic and anaerobic conditions. Scale bars are 3 µm.

The oxygen-independent fluorescence enhancement in *E. coli* cultures expressing OICP-tagged proteins under strictly anaerobic growth conditions (**Figure 3c**) and with purified OICP tags (**Figure 1h-i**) remained comparable to that under aerobic conditions. The cell-to-cell fluorescence intensity variation of **Figure 3a** and **3c** is most likely due to differences in protein expression as we observe it with dye conjugated-OICP and SfTq2 conjugated-OICP tags. We benchmarked our OICP tags against the brightest available fluorescent protein: mNeongreen (mNG)(Shaner et al., 2013). Under our experimental conditions, mNG fluorescence was completely absent under anaerobic conditions, and the fluorescence developed upon exposure of the cells to oxygen (**Figure 3d-e**). Both mNG and OICP tags expressed well under aerobic or anaerobic conditions (**Figure 3f**). The expression of OICP tags and their subsequent labeling with organic dyes do not affect the cell morphology of exponentially growing *E. coli, L. lactis*, and *S. cerevisiae* cells (**Supplementary Fig. 25**).

We illustrate the usefulness of our labeling technology by targeting proteins with Bodipy625 in the cytoplasm of live Gram-negative (*E. coli*) and Gram-positive bacteria (*L. lactis*), and the lower eukaryote *Saccharomyces cerevisiae* (**Figure 4**, and mitochondria of human embryonic kidney (HEK) cells (**Supplementary Figs. 24**). We chose Bodipy625 since it showed no detectable interactions with membranes and maximum permeability when compared to the other six fluorogenic dyes (**Supplementary Fig. 22-24**) in *E. coli, L. lactis*, and *S. cerevisiae* cells.

**Figure 4.**
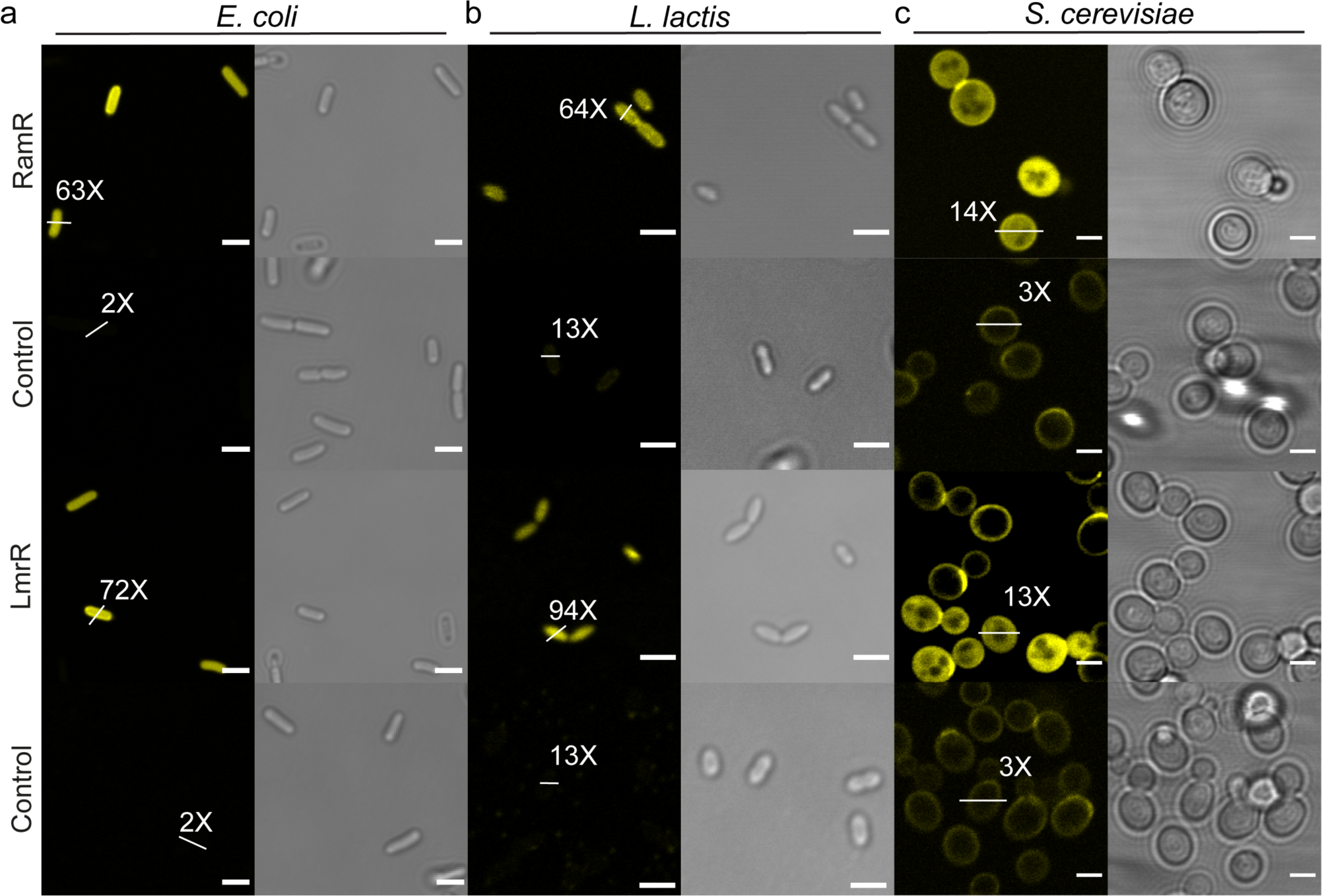
Live-cell imaging of OICP tags in prokaryotic and eukaryotic cells. (a) Fluorescence confocal and accompanying differential interference contrast (DIC) micrographs of *E. coli, L. lactis* and *S. cerevisiae* cytoplasm with Bodipy625 (yellow) from at least two independent biological replicates. White bars across arbitrarily picked cells depict signal to background fluorescence ratios. Scale bars: 3 µm.

To estimate the fraction of labeled *E. coli, L. lactis*, and *S. cerevisiae* cells, we quantified the fluorescence of live cells for the seven most useful dyes, using fluorescence flow-cytometry (**Supplementary Figs. 22-24**). The fractions of labeled cells were highest with Bodipy625 for *E. coli* (77% for LmrR and 82% for RamR) and *L. lactis* (95% for LmrR and 91% for RamR). For *S. cerevisiae* and HEK cells, only Bodipy625 entered the cytoplasm and we observe a good fraction of labeled *S. cerevisiae* cells (32% for LmrR and 53% for RamR). We note that the approach worked less well with *S. cerevisiae* than with bacteria as only Bodipy625 permeates the cell envelope of yeast and binds OICP tags. Moreover, in contrast to that in *E. coli* or *L. lactis*, significant background fluorescence was observed with Bodipy625 in HEK cells. The higher background fluorescence of some dyes is most likely due to non-specific binding to intracellular membranes and/or proteins. Additionally, we observe that Bodipy625 tends to accumulate specifically in mitochondria even in non-transfected cells(**Supplementary Figs. 24b-c**).

The OICP tags natively form high-affinity dimers, but we did not observe any fluorescent punctate spots in live cells, and thus we have no indication that our tags cause protein aggregation (**Supplementary Figs. 20-21**). Dynamic non-specific interactions with other proteins and RNA/DNA cannot be ruled out completely, but so far we have no indications that they pose a problem to the cell.

## DISCUSSION

The promiscuity in binding planar organic dyes in OICP tags stems from their biological function as transcriptional factors, proteins that bind drug-like molecules signaling the transcription of drug-exporting proteins. We have analyzed 30 planar organic dyes and observed an array of spectral effects after the binding of the dyes to the OICP tags. For some of the dyes, a distinct redshift and increase in fluorescence emission intensity (classified as *enhanced* in Table 1) is observed in the presence of both OICP tags, which correlates with an accompanying increase in dye absorbance and fluorescence lifetime. These fluorophores bind with moderate to high affinity to the OICP tags, accompanied by a significant increase in fluorescence lifetimes, indicating their future applicability in fluorescence lifetime imaging microscopy studies. In general, dyes that show enhanced fluorescence, spectral enhancements are generally more pronounced with RamR than LmrR except for DFHBI.

It is tempting to speculate that the aforementioned differences arise from the presence of a single dye-binding pocket in LmrR (formed at the dimer interface) and two apparent dye-binding pockets in RamR (**Fig. 1a**). However, the fold-changes in fluorescence scale non-linearly when comparing RamR and LmrR, which may be due to an excited-state cross-talk between the two bound dye molecules in RamR. In dyes wherein the emission fluorescence is either quenched or remain unaffected, the correlation between absorption, fluorescence, and fluorescent lifetimes is not apparent. A peculiar case is Eosin Y, wherein fluorescence lifetimes are equally enhanced in RamR and LmrR, but the fluorescence emission intensity is quenched for LmrR and enhanced for RamR. Interactions of OICP tags with AlexaFluor 488 were not detectable, suggesting little or no specificity for these molecules. The structural promiscuity of the binding sites in both proteins allows capturing of organic molecules not limited to this study, but also opens up possibilities of testing other fluorophores with desired photochemical properties in our “plug-and-play” approach. For instance, most of the MaP dyes(Wang et al., 2020) exhibit fluorogenicity with our OICP tags (2-5 fold) and enhanced fluorescence lifetimes (20%), circumventing the need for specific chemistry between the tag and the ligand (**Table 1** and **Supplementary Fig. 18**).

Although the promiscuity of the binding sites of the OICP tags enables the use of a wide range of dyes, it also precludes us from delineating precisely the underlying processes and/or interactions that contribute to the fluorogenicity and lifetime enhancements. The selective interaction of OICP tags with the organic dyes arises likely from the geometry and hydrophobicity of the binding pockets that are more accessible and/or hydrophobic than other cellular structures. We observe fluorescence enhancements of intercalators with OICP tags that are consistent with those typically observed in non-polar environments. The stability of the interaction between OICP tags and organic dyes up to 800 mM NaCl further suggests that OICP tags interact with the dyes predominantly through hydrophobic contacts. Overall, the two OICP proteins impose a somewhat different (hydrophobic) environment to the dyes, which could be exploited further by protein engineering. It should be possible to design monomeric variants of RamR by mutating residues at the dimer interface and decrease the size of the non-essential regions unimportant for fluorogenicity.

In addition to hydrophobic interactions, π−π stacking involving aromatic residues (Phe-155 in RamR and Trp96 in LmrR) is critical for dye binding(Roelfes, 2019; Yamasaki et al., 2019, 2013). Indeed, the crystal structure of RamR with Rhodamine 6G confirms the non-polar nature of the binding pocket (**Supplementary Fig. 26, panel a-b**). The presence of tryptophan, phenylalanine, and other hydrophobic amino acids in the binding pockets of OICP tags results in multiple hydrophobic contacts and π-π stacking interactions with the electron-donating and/or electron-withdrawing groups of the organic dye. These interactions likely perturb the existing electron densities and concomitantly the excitation and emission transition dipoles. The resulting properties are additionally influenced by several factors, including changes in radiative and non-radiative decay rates (affecting quantum yields), viscosity, dye conformational changes in the binding pocket not limited to tautomeric changes, reduced rotational freedom due to steric hindrance or forced planarization of out-of-plane twisted moieties influencing the HOMO-LUMO gap in the organic dye. The overall net outcome of these interactions results in a specific spectral signature for each dye-OICP pair. Importantly, the OICP tags are stable across a wide range of physicochemical conditions, including pH, temperature, ionic strength and oxygen, and environments mimicking in vivo crowding (excluded volume effects); the latter has been mimicked by using Ficoll70 as a synthetic macromolecular crowding agent. While the affinity of binding of dyes to OICP tags is not affected by Ficoll70, the fluorogenicity is decreased (**Supplementary Fig. 19**). By fine-tuning the chemical environment of the OICP tags, specific photophysical properties of the interacting dyes can be exploited for *in vivo* and *in vitro* applications.

In conclusion, we report the development and extensive characterization of two chemogenetic protein tags that enable the use of organic fluorophores for live-cell imaging and dynamic studies in bacterial, and lower eukaryotic cells and facilitate *in vitro* applications under a wide range of conditions. The current inapplicability of our OICP tags to mammalian cells calls for explorative studies with dyes not yet tested with our system and shown to have low background signals or modification of existing dyes to reduce non-specific background interactions inside the cells. Our method allows the use of cheap and widely available organic fluorophores spanning the ultraviolet-visible-infrared spectrum, fast and non-covalent labeling, and direct application in fluorescence lifetime imaging studies. The stability of our OICP tags across a wide physiological range provides an imaging tool for single-molecule studies into anaerobic gut microbes and other cells and organelles in environments low in oxygen, and visualization and physiological studies of bacteria (and their sub-compartments) living in extreme environments. We envision OICP tags to be applicable in the burgeoning field of cellular cartography, in particular for (a) real-time fluorescence monitoring for high-throughput screening of protein production and dynamics in anaerobic gut microbes(Geva-Zatorsky et al., 2015) and extremophiles(Maslov et al., 2018); (b) real-time monitoring of protein stability and turnover(Ignatova and Gierasch, 2004); (c) acquisition of long single-molecule trajectories to characterize the diffusion of proteins in the membranes; and (d) super-resolution imaging(Wang et al., 2019) and FLIM studies of cells in a wide range of environments(Adhikari et al., 2019).

## METHODS

### Construction of OICP tag plasmids

#### Escherichia coli

The pET-17b plasmids encoding *lmrR* and *ramR* under an IPTG inducible T7 promoter provided by Prof. Gerard Roelfes (University of Groningen) were used for protein expression and purification experiments. The DNA-binding capability of the LmrR was removed by substituting two lysine residues (K55 and K59) in the DNA-binding region of the protein by the negatively charged aspartic acid and neutral glutamine, respectively(Bos et al., 2013). The resulting LmrR (K55D/K59Q) allowed easier purification of the protein and has reduced interactions with DNA. For facilitating studies in *E. coli* BW25113, the corresponding plasmids for *lmrR* and *ramR* expression in the cytoplasm were created under the control of an arabinose-inducible promoter on a pBAD24 vector. Fusion constructs of *lmrR* and *ramR* with the periplasmic protein OsmY at the N-terminus of the resulting fusion were created on a pBAD24 vector harboring an arabinose-inducible promoter. Fusion constructs of *lmrR* and *ramR* with the inner membrane protein PBP5 (penicillin-binding protein 5) were created on the pNM077 vector harboring an IPTG-inducible *trc* promoter provided by Prof. Tanneke Blaauwen (University of Amsterdam). Amplification of *lmrR* and *ramR* genes and corresponding plasmid backbones for USER based cloning(Bitinaite et al., 2007) was performed using forward and reverse primers that added a uracil residue instead of a thymine residue at flanking regions (see Appendix 2 for primer sequences). The USER® reaction for ligating fragments was performed as per manufacturer’s instructions(Bitinaite et al., 2007), followed by the heat-shock transformation of chemically competent *E. coli* MC1061. Positive colonies were selected on LB-ampicillin (100 µg/ml) plates and isolated plasmids were confirmed by DNA sequencing (Eurofins Genomics Germany GmbH). Thereafter the sequence-verified plasmids were transformed into *E. coli* BW25113 and stored as glycerol stocks (30% glycerol) until further use. The DNA sequences for constructs are provided in Appendix 3.

#### Lactococcus lactis

All experiments were performed on the *L. lactis* strain NZ9000 *ΔlmrR*(Agustiandari et al., 2008). NZ9000 is a *L. lactis* MG1363 strain derivative containing the *pepN::nisR/K*(De Ruyter et al., 1996) substitution, which in the presence of the inducer, nisin A, switches on the expression of genes from the *nisA* promoter. By using the pNZ8048 vector housing the *nisA* promoter, pNZ8048-*lmrR* was constructed(Agustiandari et al., 2008). The strain NZ9000 *ΔlmrR*(Agustiandari et al., 2008) lacks a chromosomal copy of the *lmrR* gene that served as a control for experiments with OICP tags. Both NZ9000 *ΔlmrR*(Agustiandari et al., 2008) strain and the plasmid pNZ8048-*lmrR* were provided by Prof. Arnold Driessen (University of Groningen). For the construction of nisin A-inducible plasmids for *ramR*, we cloned the native *ramR* gene into a pNZC3GH vector harboring a nisinA-inducible *nisA* promoter. However, we failed to observe any protein expression with this construct, which possibly arose because of codon-mismatch from the GC-rich native *ramR* sequence (Appendix 3). Next, we codon-optimized the *ramR* gene for *L. lactis* using the graphical codon usage analyzer tool(Fuhrmann et al., 2004) and synthesized the *ramR* gene fragment(Geneart®, Regensburg, Germany). The amplification of the *ramR* gene and corresponding plasmid pNZC3GH backbone for USER based cloning(Bitinaite et al., 2007) was performed using forward and reverse primers that added a uracil residue instead of a thymine residue at flanking regions. The resulting plasmid pNZC3GH*-ramR* was confirmed by DNA sequencing (Eurofins Genomics Germany GmbH) and transformed into the *L. lactis* strain NZ9000 *ΔlmrR*(Agustiandari et al., 2008) by electroporation. The native and codon-optimized DNA sequences for *ramR* are provided in Appendix 3.

#### Saccharomyces cerevisiae

We chose the cytoplasmic protein adenylosuccinate synthase (Ade12) for studies with our OICP tags because of uniform cytoplasmic fluorescence(Munder et al., 2016). For plasmid cloning, *E. coli* MC1061 was used for cloning and plasmid storage. The *ade12* gene was amplified using PCR from *S. cerevisiae* BY4742 chromosomal DNA using forward and reverse primers that added a uracil residue instead of a thymine residue at flanking regions. For plasmid backbone amplification, the multicopy plasmid pRSII426 (housing the selectable *ura3* gene and allowing expression of a target protein from a constitutive ADH1 promoter) was used with forward and reverse primers that added a uracil residue instead of a thymine residue at flanking regions. The USER® reaction was performed as per manufacturer’s instructions(Bitinaite et al., 2007) followed by the heat-shock transformation of chemically competent *E. coli* MC1061. Positive colonies were selected on a LB chloramphenicol (32 µg/ml) plates and isolated plasmids were confirmed by DNA sequencing (Eurofins Genomics Germany GmbH).

The correct plasmids were then transformed to *S. cerevisiae* strain BY4709 strains lacking the *ura3* gene enabling uracil based selection. Transformation of plasmids into *S. cerevisiae* was performed as described elsewhere(Drew et al., 2008) with some minor modifications. In short, single colonies were inoculated into 5 ml yeast extract peptone dextrose (YPD) media and incubated at 30 °C, 200 rpm overnight. The following day cells were diluted to OD_600_ ∼0.1 in 50 ml of media and grown at 30 °C, 200 rpm until a target OD_600_ ∼0.4-0.6 was reached. Once the target OD_600_ was reached cells were pelleted, the supernatant was removed and pellets resuspended in sterile H_2_O to wash. Cells were kept on ice throughout the transformation procedure. The wash step was repeated and then cells were resuspended in 1 ml 0.1 M lithium acetate. Cells were pelleted and resuspended in the required volume of 0.1 M Lithium acetate before 50 µl was added to (50 %w/v) PEG4000 and samples were vortexed until homogenous. 25 µl of (2 mg/ml) single-stranded salmon sperm was added to the cell suspension and vortexed again. Finally, 50 µl of corresponding plasmid DNA (250 ng - 1 µg) was added to the cell suspension before vortexing and incubation at 30 °C with shaking for 30 minutes. Heat-shock was carried out for 25 minutes at 42 °C with shaking before cells were pelleted, resuspended in 200 µl of sterile H_2_O, and plated onto uracil lacking agar plates. After plates were incubated at 30 °C for 48 - 72 hours single colonies were selected for re-streaking on selective agar plates to attain a monoclonal population. Single colonies from the monoclonal population were then selected for confirmation of positive clones by plasmid isolation and subsequent sequencing of the coding region (Eurofins Genomics Germany GmbH).

#### HEK-293T cells

pmTurquoise2-Mito was a gift from Dorus Gadella(Goedhart et al., 2012) (Addgene plasmid # 36208). Cox8A-RmrR-FLAG is a fusion construct comprising of 29 amino acids of COX8A (a mitochondrial targeting signal), RamR codon-optimized for mammalian expression and a FLAG-tag. Expression was in pcDNA3.1 from the constitutive CMV (human cytomegalovirus) promotor. The complete sequence of the RamR construct used is given in Appendix 3.

### Protein expression and purification

Chemically competent BL21(DE3) *E. coli* cells were transformed with a pET17b expression vectors carrying cyto-LmrR and cyto-RamR constructs under the control of a T7 RNA polymerase promoter (p(*T7*)). Single colonies were picked and inoculated into a starter culture of 10 ml of fresh lysogeny broth (LB) medium (10 g/l tryptone, 5 g/l yeast extract, and 10 g/l NaCl) containing 100 µg/mL of ampicillin and grown at 37 °C with 180 rpm shaking overnight. The following day, the saturated LB culture was diluted 100-fold in a 500 ml fresh LB medium in a 2 l Erlenmeyer flask containing 100 µg/ml of ampicillin and allowed to grow at 37 °C with 180 rpm shaking until the culture reached an OD_600_ of 0.84-0.90. At this stage, isopropyl β-D-1-thiogalactopyranoside (IPTG) at a final concentration of 1 mM was added to induce the expression of the target protein and expressions were carried out at 30 °C with 180 rpm shaking overnight. The next day, cells were harvested by centrifugation (6000 rpm, JA10, 20 min, 4 °C, Beckman). The pellet was re-suspended in 15-20 ml of 50 mM NaH_2_PO_4_, pH 8.0, 150 mM NaCl with half a tablet of mini complete EDTA-free protease inhibitor cocktail (Roche) and 1mM of the serine protease inhibitor Phenylmethanesulfonyl fluoride (PMSF). Next, DnaseI (final concentration, 0.1 mg/mL) and MgCl_2_ (final concentration, 10mM) were added. Sonication was carried out (tip diameter 6mm, 75% (200W) for 8 min (10 s on, 15 s off). Additional sheer forcing with a syringe and a long needle was applied for at least 2 times. Henceforth, all steps were carried out at 4°C.

The cell lysates obtained after sonication were centrifuged (16000 rpm, JA-17, 45 min, 4 °C, Beckman) to remove unlysed cells and other high molecular weight debris. The cell-free extract was then filtered and equilibrated with 5 ml of pre-equilibrated Strep-Tactin® column material for 1 h (mixed at 200 rpm on a rotary shaker). The column was washed with 3 x 2 CV (column volume) of re-suspension buffer (same as buffer used before) and eluted multiple times with 0.5 CV (6-7 times) of elution buffer (resuspension buffer containing 5 mM desthiobiotin). Fractions were analyzed on a 12% polyacrylamide SDS-Tris Tricine gel followed by Coomassie Blue staining. Fractions containing protein (excluding the first elution) were concentrated in centrifugal filters and re-buffered to 20 mM K-MOPS, pH 7.0, 150 mM NaCl using dialysis (reduces DNA contamination in elution fractions). The concentration of purified LmrR and RamR was determined by using the calculated extinction coefficient obtained from Protparam on the Expasy server (ε_280_ for LmrR monomer = 25,440 M^-1^ cm^-1^ and that of RamR monomer = 29,450 M^-1^ cm^-1^). Expression yields typically were 30-40 mg/l for LmrR and 40-50 mg/l for RamR. Aliquots of 500 µl were flash-freezed using liquid nitrogen until use. Thawed protein samples for analysis were not frozen a second time and used within 24 hours.

For purification of HisTag containing proteins, a Ni^2+^-Sepharose resin was used. The resin was pre-equilibrated in 50 mM KPi, 150 mM NaCl, pH 7.0, with 10 mM imidazole. The cell-free extract was added to the Ni2+-Sepharose resin (0.5 ml bed volume per 10 mg total protein) and nutated for 3 h after which the resin was washed with 20 column volumes of 50 mM KPi, 150 mM NaCl, pH 7.0, with 50 mM imidazole. Proteins were then eluted with 50 mM KPi, 150 mM NaCl, pH 7.0, with 500 mM imidazole in 500 µl aliquots. The most concentrated fractions were run on a Superdex15 Increase 10/300 GL size-exclusion column (GE Healthcare) in 20 mM K-MOPS, pH 7.0 with 150 mM NaCl. Protein containing fractions were pooled and concentrated to 5 mg/ml in a Vivaspin 500 (3 kDa) centrifugal concentrator (Sartorius AG), after which they were aliquoted, flash-frozen in 500 µl aliquots and stored at -80 °C.

### Absorption and fluorescence spectroscopy

Organic dyes were prepared as stock solutions (2.5–5 mM) in DMSO and diluted for spectroscopy to a final concentration of 1 µM in 20 mM K-MOPS, 150 mM NaCl, pH 7.0 such that the DMSO concentration did not exceed 0.5% (v/v). Purified RamR and LmrR were then added and incubated for less than a 1 minute before the measurement of absorption and fluorescence spectra. All measurements (3 independent replicates) were taken at 30 °C. Samples were incubated for 1-2 minutes after mixing gently with a pipette and thereafter measured in black polystyrene, µClear® bottom, 96-well plates (Greiner Bio-One, cat. 655096) and for samples having excitation wavelength < 400 nm on 96-well plates with a UV-compatible optical bottom (Greiner Bio-One, cat. 655801) on a TECAN Spark® 10M microplate reader. Reported values are averages of 3 independent experiments. The obtained results are concisely summarized in **Table 1** in the main text and an extended version of the table with details of organic dyes and error bars for each measurement are summarized in **Supplementary Table 1**. Experiments to determine bound fraction was performed for each dye by keeping the dye concentration constant to avoid any inner filter effects due to increasing dye concentrations. By keeping the dye concentration constant at 1 µM, protein concentrations were increased until the fluorescence emission intensity saturated resulting in an apparent binding curve. The association constant K_a_ was then estimated by fitting the data points to a Hill model. The same samples were also used for absorption spectroscopy to validate if the spectral changes observed in fluorescence emission emanated from changes in absorption. Fluorescence emission and absorption spectra of the organic dyes in presence of OICP tags were normalized against that of the organic dyes alone giving a fold-change (from peak values) in absorption and fluorescence emission(panel c and e in **Supplementary Figs. 2-18**). At saturation values of OICP tags with the organic dyes, a normalized fluorescence emission spectra and absorption spectra are depicted in panel d and f in **Supplementary Figs. 2-18**. The excitation and emission bandwidths were set to 5 nm for all measurements.

For pH, NaCl and temperature scans, purified RamR and LmrR were used at a final concentration of 4 µM and 50 µM, respectively, in 20 mM MOPS, 150 mM NaCl, pH 7.0 with Bodipy495. The buffer pH was adjusted using KOH (K-MOPS). Buffer exchange for pH scan measurements was performed using Thermo Scientific™ Zeba™ Spin desalting columns and the pH was verified subsequently using a pH meter. For pH values in the range 4-7, a citric acid-Na_2_HPO_4_ buffer was used supplemented with 150 mM NaCl. For pH values between pH values 6.5 and 8, 50 mM sodium phosphate buffers supplemented with 150 mM NaCl were used at 30 °C keeping the dye and OICP tag concentration constant. To probe the stability of our OICP tags with increasing ionic strengths, we modulated the concentration of NaCl in 20 mM K-MOPS, 150 mM NaCl, pH 7.0 buffer keeping the dye, and OICP tag concentration constant. The color-shaded regions represent the standard deviation (s.d.) over three independent measurements.

Temperature scan measurements were performed in Teflon sealed quartz cuvettes in an FP-8300 spectrofluorimeter (Jasco, Inc.) equipped with a temperature modulation system (Julabo GmbH). The temperature rise gradient was 1 °C/min and an additional minute was allowed to equilibrate samples at a given temperature. Samples were excited at 480 nm and the emission spectra were acquired from 495 to 600 nm with 1 nm data interval and 5°C temperature intervals from 20 °C to 60 °C. Excitation and emission bandwidths were kept constant at 5 nm. Reported values are averages of independent experiments. For temperature scan, beyond 55 °C, visible white precipitates could be observed for OICP tags indicated with a dotted line in **Fig. 1f**.

For measurements under strictly anaerobic conditions, 20 mM K-MOPS, 150 mM NaCl buffered at pH 7.0 was prepared and equilibrated in the CO_2_ hood at least for a week before measurements.

### Fluorescence lifetime

The fluorescence lifetime measurements were acquired at a 10 MHz repetition rate for 30 seconds on a MicroTime 200 confocal microscope (PicoQuant, Berlin, Germany) on glass-bottom dishes (Willco Wells®, cat. HBST-3522). Purified RamR and LmrR were used at a final concentration of 4 µM and 50 µM respectively in 20 mM K-MOPS, 150 mM NaCl, pH 7.0 with 1 µM of organic dye. The exponential-tail fitting of the lifetime decay was done with the SymphoTime software. Reported values are averages of independent experiments and the accompanying error bars represent s.d. at ambient temperature (∼ 25 °C). The laser excitation modules employed in the MicroTime 200 confocal microscope were 440 nm, 485 nm, 532 nm, 595 nm, and 640 nm.

### Fluorescence confocal microscopy and phase contrast microscopy

#### Preparation of glass slides

To ensure the immobility of *E. coli* cells we used (3-aminopropyl)triethoxysilane (APTES)-treated glass cover slides. The glass slides were first cleaned by sonicating them for 1 hr in 5 M KOH, followed by rinsing at least 10 times with MilliQ and blowing off the remaining MilliQ with pressurized nitrogen. Next, the glass slides were immersed in acetone containing 2 % v/v APTES for 30 minutes at room temperature. Thereafter we removed the acetone and APTES and rinsed the slides 10 times with MilliQ. Again, the remaining MilliQ was blown off with pressurized nitrogen. The glass slides were used within 2 days of preparation. The cells were concentrated to an OD_600_ ∼ 1 and after adding 20 µl cells on the APTES slide, a clean object slide was put on the top and the whole assembly was put up inverted in the microscope stage.

For *L. lactis* cells, the glass slides were first cleaned by sonicating them for 1 hr in 5 M KOH, followed by rinsing at least 10 times with MilliQ and blowing off the remaining MilliQ with pressurized nitrogen. The glass slides were used within 2 days of preparation. The cells were concentrated to an OD_600_ ∼ 1 and after adding 20 µl cells on the cleaned slide, a clean object slide was put on the top and the whole assembly was put up inverted in the microscope stage.

For *S. cerevisiae* cells, a similar glass slide cleaning protocol as aforementioned for *L. lactis* cells was followed and the cleaned slides were used in a stick-Slide 8 well chamber (Ibidi GmbH, cat. 80828). The cells were concentrated to an OD_600_ ∼ 0.5 and 100 µl cells were added in each chamber.

#### Preparation of E. coli for fluorescence confocal microscopy and phase contrast microscopy

For each experiment, a glycerol stock of *E. coli* BW25113 with desired LmrR/RamR variant on a pBAD plasmid was stabbed with a sterile pipette tip and deposited in 3 ml lysogeny broth (LB Lennox: 10 g/l tryptone, 5 g/l yeast extract, 5 g/l NaCl) containing 0.2 % v/v glycerol and 100 µg/mL ampicillin. The LB medium was then incubated at 37 °C with 200 rpm shaking. The next day, the saturated LB culture was diluted 100-fold in a 3 ml fresh LB medium containing 0.2 % v/v glycerol and 100 µg/ml ampicillin. The LB medium was then incubated at 37 °C with 200 rpm shaking until the culture reached an OD_600_ of 0.7-0.8. At this stage, protein expression was induced using 0.1% w/v arabinose, and the cultures were then incubated at 30 °C with 200 rpm shaking overnight. The following day, the saturated LB culture was diluted 100-fold in a 3 ml fresh LB medium containing 0.2 % v/v glycerol, 0.1% arabinose, 100 µg/ml ampicillin and allowed to grow until the culture reached an OD_600_ of 0.4-0.6. These cells were now directly used for labeling and consecutive imaging. For dye labeling, 0.5 ml of OD_600_ of 0.6 cultures were centrifuged at 11000 g for 1 minute and a final concentration of 15 µM dye was added. The pellet was gently resuspended and the cell suspension was kept at 30 °C for 30 minutes. Thereafter, the suspension was centrifuged at 11000 g for 1 minute and the cell pellet was washed 3 times with 1 ml of LB medium. The washing step was repeated 3 times to ensure the removal of free dye. The resulting suspension was now used for confocal fluorescence microscopy and phase-contrast microscopy.

#### Preparation of L. lactis for fluorescence confocal microscopy and phase contrast microscopy

For experiments with LmrR, a glycerol stock of *Lactococcus lactis* NZ9000 *ΔlmrR* cells with desired LmrR/RamR variant on a nisin-inducible plasmid was stabbed with a sterile pipette tip to obtain a small number of cells. *Lactococcus lactis* NZ9000 was grown in M17 medium (Difco, Franklin Lakes, NJ, USA) supplemented with 1% (w/v) glucose (GM17) and 5 µg/ml chloramphenicol at 30°C without shaking. We incubated the cultures at 30 °C, without shaking since *L. lactis* is facultatively anaerobic. The next day, the saturated culture was diluted 100X in a fresh 3 ml GM17 medium containing 5 µg/ml chloramphenicol and incubated at 30 °C without shaking until OD_600_ reached 0.3. At this stage, we added 2 µl of nisinA solution (filtered supernatant from a *L. lactis* NZ9700 culture), and the culture was allowed to grow overnight at 30 °C without shaking. On the morning of the next day, about 100 µl of culture was added to 4 ml of fresh GM17 medium containing 5 µg/ml chloramphenicol and 2 µl nisinA solution, to yield an OD_600_ ∼ 0.1. The cultures were then incubated at 30 °C until OD_600_ reached 0.4. These cells were now directly used for labeling and consecutive imaging. For dye labeling, 0.5 ml of OD_600_ of 0.6 cultures were centrifuged at 11000 g for 1 minute and a final concentration of 15 µM dye was added. The pellet was gently resuspended and the cell suspension was kept at 30 °C for 30 minutes. Thereafter, the suspension was centrifuged at 11000 g for 1 minute and the cell pellet was washed 3 times with 1 ml of LB medium. The washing step was repeated 3 times to ensure the removal of free dye. The resulting suspension was now used for confocal fluorescence microscopy and phase-contrast microscopy.

#### Preparation of S. cerevisiae for fluorescence confocal microscopy, phase contrast microscopy, and flow cytometry experiments

*S. cerevisiae* was grown in a minimal synthetic defined (SD) media including a yeast nitrogen base lacking riboflavin and folic acid, ammonium sulfate, and 2% glucose as a carbon source. Riboflavin was not added to prevent interactions with OICP tags. For each experiment, single colonies from uracil lacking synthetic defined (SD *URA-*) plates were inoculated into 5 ml SD URA- media and incubated at 30 °C, 200 rpm overnight. The following day cells were diluted to OD_600_ ∼0.02 in 5 ml of media and grown at 30 °C, 200 rpm, and maintained in exponential phase for three consecutive days. On the day of the experiment, once the OD_600_ reached 0.5, cells were pelleted at 8000g for 1 minute in 1.5 ml sterile Eppendorf tubes, the supernatant was removed and pellets resuspended in sterile SD *URA-*media to a final OD_600_ of 1. These cells were now directly used for labeling and consecutive imaging. For dye labeling, 0.5 ml of OD_600_ of 0.6 cultures were centrifuged at 11000 g for 1 minute and a final concentration of 15 µM dye was added. The pellet was gently resuspended and the cell suspension was kept at 30 °C for 30 minutes. Thereafter, the suspension was centrifuged at 11000 g for 1 minute and the cell pellet was washed 3 times with 1 ml of LB medium. The washing step was repeated 3 times to ensure the removal of free dye. The resulting suspension was now used for confocal fluorescence microscopy and phase-contrast microscopy.

#### Preparation of mammalian cells (HEK293T) for transfection and fluorescence confocal microscopy

100,000 HEK293T cells were cultured in a 35 mm imaging dish with a glass bottom and an imprinted 50 µm cell location grid (Ibidi; Cat.No:81148) in DMEM medium supplemented with 1% (v/v) Na-pyruvate, 1% (v/v) antibiotics, 1% (v/v) glutamine and 10% (v/v) fetal calf serum (FCS). JetPEI (PolyPlus; Cat.No: 101-10N) was used to co-transfect cells with pmTurquoise2-Mito and Cox8-RamR-FLAG constructs. 16 hr post-transfection, the cells were washed with phenol-red free, serum-free, antibiotic-free RPMI imaging medium (ThermoFisher; Cat.No: 11835030) containing 1% glutamine. Bodipy625 was diluted in the imaging medium to a final concentration of 450 nM. After 15 min of incubation at 37°C, the free dye was washed away with the imaging medium. Cells were imaged live at 37°C by confocal imaging and their positions on the grid were marked. Subsequently, the cells then were washed with PBS, fixed with 4% paraformaldehyde (PAF) for 15 min, and permeabilized with 0.1% Triton-X100 in PBS for 5 min. Cells were immunolabeled with mouse IgG1 Anti-Flag antibody (Sigma; Cat. No: F1804) over-night at dilution 1:200 in PBS. Next, cells were washed with PBS and labeled with secondary donkey-anti-mouse antibody conjugated to Alexa Fluor 568 (ThermoFisher; Cat.No: A10037) at dilution 1:400 in PBS for 30 min. Finally, cells located at the stored positions on the grid were imaged by confocal microscopy.

#### Imaging

Confocal laser scanning microscopy (LSM 710, Carl Zeiss AG Jena, Germany) equipped with a C-Apochromat 40x/1.2 NA objective was used for *in vivo* fluorescence imaging of live *E. coli, L. lactis, S. cerevisiae* cells. 405 nm, 488 nm 543 nm and 632 nm lasers were employed for fluorescence excitation. For all measurements, data were acquired within 20 minutes and thereafter a fresh slide was used. The stage temperature was maintained at 30°C. We recorded 16-bit images at randomized positions on the glass slide with 512 x 512 pixels (34.19 µm x 34.19 µm) and analyzed at least 100 cells for each dye with a corresponding OICP tag. All images were collected under identical conditions of power and gain for a given dye. For anaerobic fluorescence imaging, experiments were performed in a sterile glove box maintained constantly under a 5% CO_2_ environment. Control experiments to verify oxygen unavailability were performed by assessing mNG fluorescence in *E. coli* BW25113 housing a pBAD-*mNG* plasmid in a TECAN multi-well plate reader inside the glove box. To avoid any recovery of the fluorescent protein mNeonGreen (mNG) fluorescence during cell harvesting, all steps until slide preparation were done in the sterile glove box. The same samples when exposed to air gained fluorescence with saturated over time. For anaerobic imaging, cytoplasmic LmrR and RamR proteins were expressed from a pBAD24 plasmid under conditions identical to those performed in presence of oxygen.

Phase-contrast images were acquired using an Axio Observer Z1 microscope (Carl Zeiss, Jena, Germany) equipped with a C-Apochromat 100x/1.49 NA objective for imaging of live *E. coli, L. lactis, S. cerevisiae* cells. For all measurements, data were acquired within 20 minutes and thereafter a fresh slide was used. The stage temperature was maintained at 30°C. We recorded 16-bit images at randomized positions on the glass slide with 1024 x 1024 pixels (66.05 µm x 66.05 µm) and analyzed at least 100 cells for each dye with a corresponding OICP tag. Cell aspect ratios (length/width) were obtained using the MicrobeJ plugin in Fiji. Fiji(Schindelin et al., 2012) was used for all image analysis of confocal and phase-contrast microscopy images.

For imaging HEK cells, a confocal laser scanning microscope (LSM800, Carl Zeiss AG Jena, Germany) equipped with a 63X oil immersion objective was used. A 640 nm laser was employed for Bodipy625 excitation. The stage temperature was maintained at 30°C. All images were collected under identical conditions of power and gain for a given dye.

### Fluorescence Recovery After Photobleaching (FRAP)

We performed fluorescence recovery after photo-bleaching (FRAP; see Figure 1a,b) on an LSM710 Zeiss confocal laser scanning microscope (Zeiss, Oberkochen, Germany) as reported previously by our group(Mika et al., 2014; Schavemaker et al., 2017), based on a previously described method(Elowitz et al., 1999). We programmed the microscope to take three images (pre-bleach), then photobleached the cell at one of the poles, and finally record the recovery of the fluorescence over time. We ensured that we picked cells lying flat on one position without exhibiting any rotational or translational motion during the time of measurement, not undergoing cell division, and having no neighbors that would obscure the analysis.

### Flow cytometry

Live cells were prepared and labeled identically as for confocal fluorescence microscopy. For *E. coli, L. lactis* and *S. cerevisiae* cells, firstly using the FSC/SSC gating, cell debris was removed from the main cell population. A positivity threshold gate for each sample was defined based on unlabeled (0%) and labeled control cells expressing no protein (< 3%). An identical positivity threshold gate was applied to all samples for a given organic dye. Samples were measured on an LSR-II flow cytometer (BD Bioscience) with 10000 events for each sample and analyzed with Kaluza Analysis 2.1 software (Beckman Coulter, CA, USA). The data was acquired (18 bits digitalization in 5 decades) using DIVA 8.0 software and saved as FCS 3.0 or 3.1 files. The following laser and corresponding filter sets (dye: laser; peak/bandwidth) were employed: a) DHFBI: 405; 525/15,b) Bodipy488 and Bodipy495: 488; 530/30, c) Rhodamine 6G and Rose Bengal: 561; 585/15, d) Bodipy589: 561; 615/20 and e) Bodipy625: 635; 660/30). At least two independent biological replicate measurements were performed for each sample. For DFHBI, the excitation laser and the fluorescence filter sets are not ideal, which may result in an underestimation of the fraction of labeled cells. The gating strategy is given in **Appendix 4**. For HEK293T cells, a starting cell population per sample was collected with the stopping rule of 30,000 events per preliminary gate drawn in FSC/SSC in a CytoFlex LX (Beckman Coulter) flow cytometer. Next, the cells were analyzed placing gates on PE (phycoerythrin) channel indicating FLAG expression labeled with Alexa Fluor 568. APC (allophycocyanin) filter was used to detect Bodipy-625 fluorescence. Negative/high background populations were defined by unstained cells visible in the PE channel and un-transfected but Bodipy-625 pulsed cells in APC channel. All experimental data were analyzed in Kaluza Analysis 2.1 software.

### Size-Exclusion Chromatography with Multiangle Laser Light Scattering (SEC-MALLS) detection

The column was equilibrated with 20 mM K-MOPS, 150 mM NaCl (pH 7.0). The Superdex 200 column used for the SEC-MALLS analysis was equilibrated with 20 mM K-MOPS, 150 mM NaCl buffered at pH 7.0, filtered through 0.1 µm pore size VVLP filters (Millipore)) and subsequently the buffer was recirculated through the system for 16 h at 0.5 ml/min. This allowed the buffer to pass several times through the degasser and the pre-injection filter, thereby thoroughly removing air and particles and allowed a judgement of the stability of detector baselines. 400 µl of protein solution (0.4 mg/ml) was injected and the data from the three detectors were imported by the ASTRA software package version 5.3.2.10 (Wyatt Technologies). For instrument calibration and analysis, aldolase protein was used as an internal standard.

### Gel electrophoresis and Western blot

*E. coli* cells expressing cytoplasmic LmrR, RamR and mNG tagged with a C-terminal 6-HisTag were initially centrifuged at 11000g to remove spent media and the cell pellet was resuspended in 50 mM KPi, 100 mM NaCl (PBS) buffered at pH 7.0 to a final OD_600_ of 6. Samples were thereafter loaded with 5X SDS loading buffer, heated at 90°C for 5 minutes, and were separated using a 15% SDS-polyacrylamide gel. Proteins were blotted onto polyvinylidene difluoride (PVDF) membranes for 35 minutes at a constant current of 0.08 A. Blots were blocked for 2 hours in freshly prepared 0.2%(w/v) I-block® in 0.2% (w/v)Tween20 containing 50 mM KPi, 100 mM NaCl (PBST) buffered at pH 7.0. HRP-conjugated anti-HisTag antibodies were incubated for 1 hour at 1:6000 dilution in PBST buffer and washed 3 times for 5 min in 0.2%(w/v) I-block® in PBST buffer, followed by washing 3 times for 5 minutes in PBS buffer. For the chemiluminescent readout, the Super Signal West Pico (Thermo Scientific) substrate was used according to the manufacturer’s instructions. Chemiluminescent detection was performed on a LAS-4000 mini (Fujifilm, Düsseldorf, Germany).

## Supporting information

Supplemental Methods and Data for Chemogenetics Paper Iyer et al

## ACKNOWLEDGMENTS

We thank our colleagues in the Membrane Enzymology group at the University of Groningen for their valuable suggestions; Dr. Jonas Cremer (University of Groningen/Stanford University) who helped with the cultivation and analysis of cells under strictly anaerobic conditions; Prof. Tanneke Blaauwen (University of Amsterdam) who kindly provided the fluorescent SuperFolder mTurquoise (sfTq2) and PBP5-sfTq2 constructs; Geert Mesander of the University Medical Center Groningen for assistance in fluorescence-activated cell sorting measurements; Dr. Viktor Krasnikov for assistance in fluorescence lifetime measurements; Prof. Kai Johnsson for providing the MaP dyes; Dr. Arnold Driessen (University of Groningen) for kindly providing the pNSC8048 plasmid (carrying the lmrR gene) and *L. lactis* lmrR knockout strain, and Michele Partipilo for providing the pNZC3GH plasmid. This research was funded by an ERC Advanced grant (ABCVolume; #670578).

## AUTHOR INFORMATION

## AUTHOR CONTRIBUTIONS

A.I., S.C., G.R., GvdB, and B.P. conceived the project and designed the experiments. A.I. and S.C. designed and performed cloning experiments in *E. coli, L. lactis* and *S. cerevisiae* and purified OICP tags. A.I. performed most of the bacterial cloning, microscopy experiments, and analyzed the data. A.I. and A.J.F. designed and performed cloning and microscopy experiments in *S. cerevisiae*. M.B. designed and performed experiments in mammalian cells. A.I. and B.P. wrote the paper.

## ADDITIONAL FILES

The complete *in vitro* and *in vivo* characterization of 30 organic dyes with OICP tags, representative images of labeled *E. coli, L. lactis, S. cerevisiae*, and HEK cells with 7 fluorogenic dyes, the effect of Ficoll70 on fluorogenicity, the gating strategy for flow cytometry and accompanying data with the fraction of labeled cells, the effect of dye labeling on cell morphology, the strains, and plasmids used in the study, is available in supporting information.

## COMPETING INTERESTS

The authors declare no competing financial interests.

## ABBREVIATIONS

BODIPY: 4,4-difluoro-4-bora-3a,4a-diaza-s-indacene;
s.d.: standard deviation;
HOMO: highest occupied molecular orbital;
LUMO: lowest unoccupied molecular orbital

## Notes

### Competing Interest Statement

The authors have declared no competing interest.

