## Supplemental Methods and Data for Chemogenetics Paper Iyer et al for "Oxygen-independent chemogenetic protein tags for live-cell fluorescence microscopy"

**Supplementary Figures 1-26**

**Appendix 1: Strains used in the study**

**Appendix 2: Plasmids used in the study**

**Appendix 3: DNA sequences (5' to 3') of constructs used in the study**

**Appendix 4: Gating strategy used for flow cytometry studies**

**Supporting references**

a

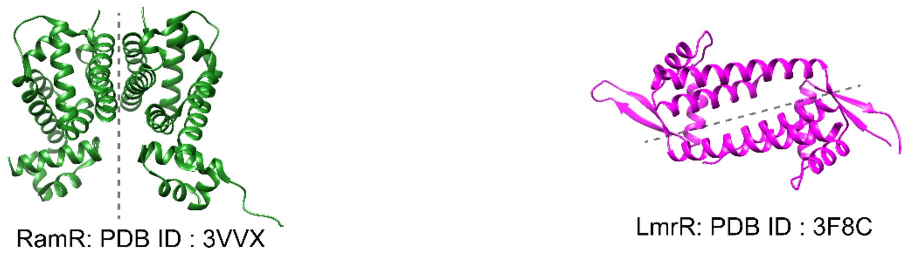


b

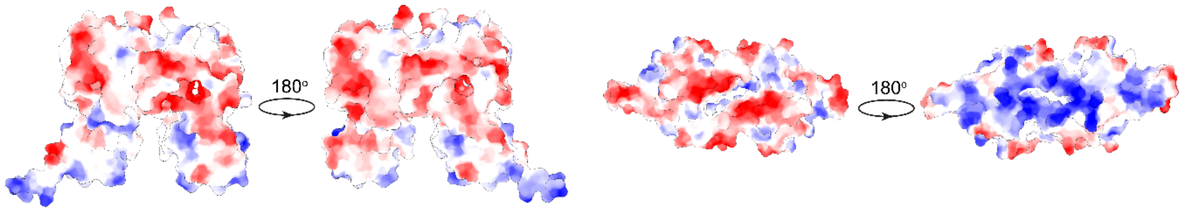


c

Primary sequence: RamR
MVARPKSEDKKQALLEAATQAIAQSGIAASTAVIAR
NAGVAEGTLFRYFATKDELINTLYLHLKQDLCQSMI
MELDRSITDAKMMTRFIWNSYISWGLNHPARHRAI
RQLAVSEKLTKETEQRADDMFPELRDLCHRSVLM
VFMSDEYRAFGDGLFLALAETTMDFAARDPARAG
EYIALGFEAMWRALTREEQ

Primary sequence: LmrR
MGAEIPKEMLRAQTNVILLNVLKQGDNYVYGIIKQV
KEASNGEMELNEATLYTIFDRLEQDGIISSYWGDES
QGGRRKYYRLTEIGHENMRLAFESWSRVDKIIENLE
ANKKSEAIKAA

d

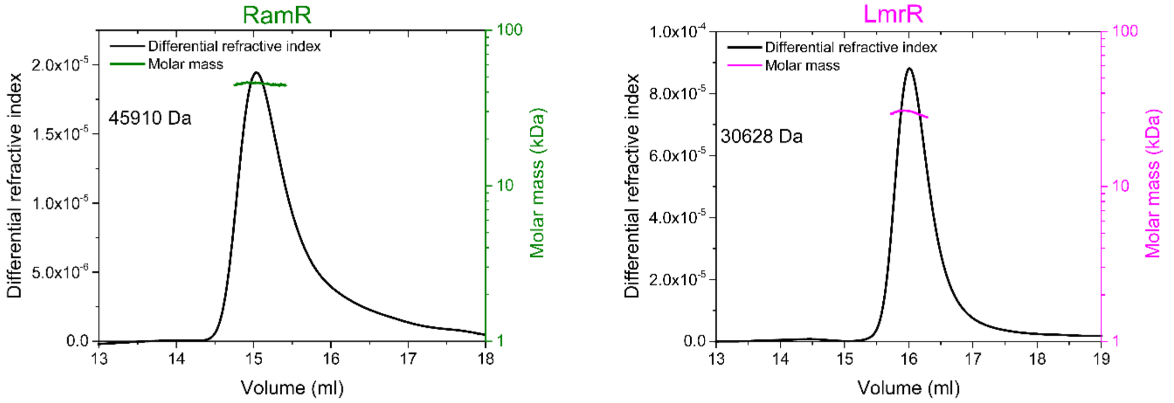


**Supplementary Figure 1:** (a) Protein structures by x-ray crystallography of oxygen-independent chemogenetic protein (OICP) tags: RamR and LmrR. (b) Coulombic surface mapping of the electrostatic potential of OICP tags depicted red for negative potential, white near neutral, and blue for positive potential. (c) The primary sequences of the OICP tags used in the study. (d) Size-exclusion chromatography coupled the multi-angle light scattering (SEC-MALS) analysis of OICP tags to determine the native molecular weight of the proteins. Black lines, signals from the refractive index detector (left-hand Y-axis); solid colored lines, calculated protein molecular weights (right–hand Y-axis). RamR (green) and LmrR (magenta) were observed to have a molecular weight of ~45.9 kDa and ~30.6 kDa respectively, which correspond to dimeric structures. All experiments were performed at 30 °C in 20 mM K-MOPS, 150 mM NaCl buffered at pH 7.0.

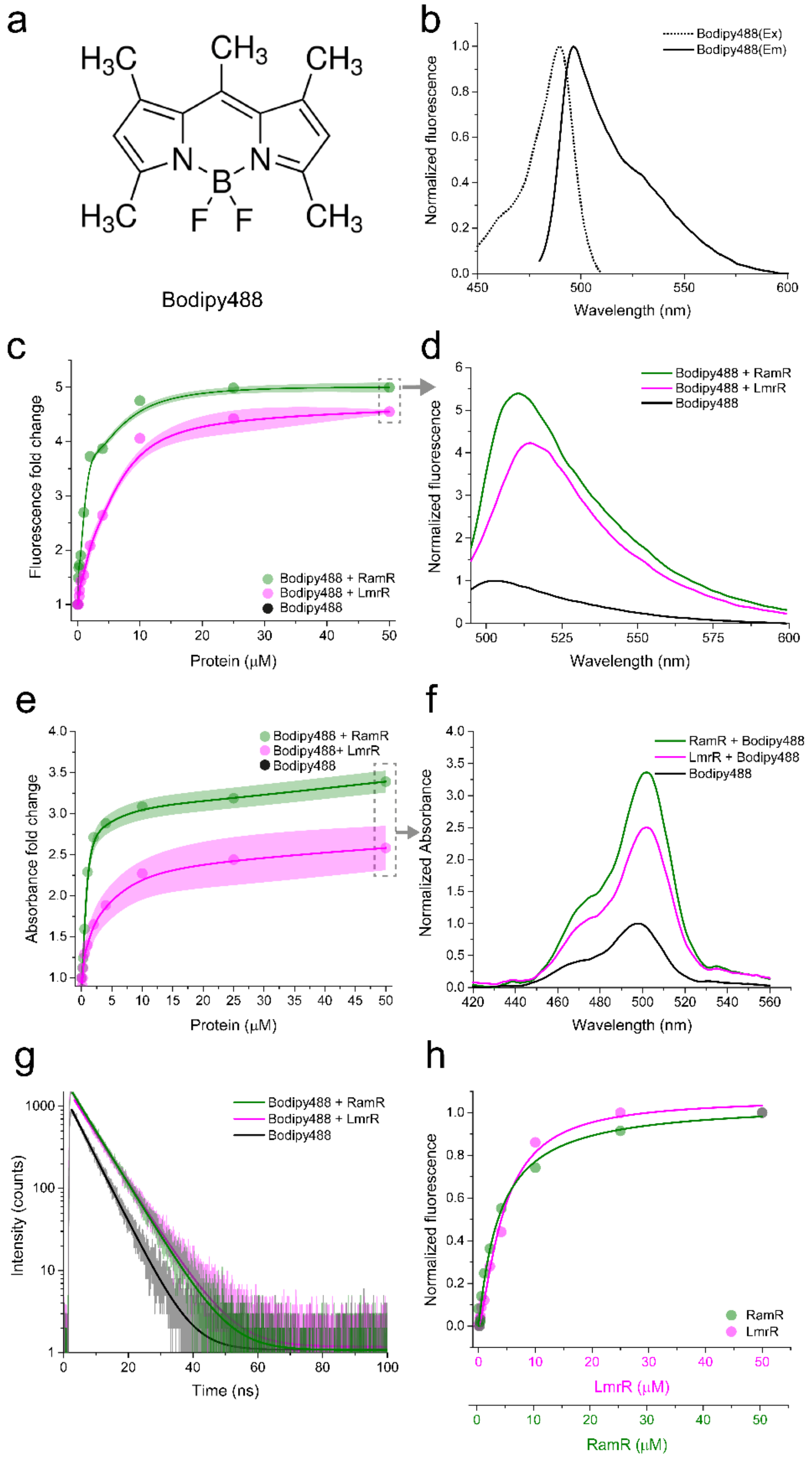


**Supplementary Figure 2**: Characterization of OICP tags (RamR: green and LmrR: magenta) with Bodipy488. (a) Structure of Bodipy488. (b) Excitation (dotted line) and emission (solid line) spectra of Bodipy488. (c) Fluorescence fold change of Bodipy488 on titration with OICP tags. (d) Fluorescence emission spectra of Bodipy488 with OICP tags at a protein:dye molar ratio of 50:1. (e) Absorbance fold-change of Bodipy488 on titration with OICP tags. Solid lines represent spline fits and shaded regions represent s.d. over three independent measurements in panel (c) and (e). (f) Absorption spectra of Bodipy488 with OICP tags at a protein:dye molar ratio of 50:1. (g) Fluorescence lifetime spectra of Bodipy488 with OICP tags at a protein:dye molar ratio of 50:1 fit with a mono-exponential decay function (solid lines). (h) Bound fraction of Bodipy488 with OICP tags ascertained from a Hill fit. All experiments were performed at 30 °C in 20 mM K-MOPS, 150 mM NaCl buffered at pH 7.0.

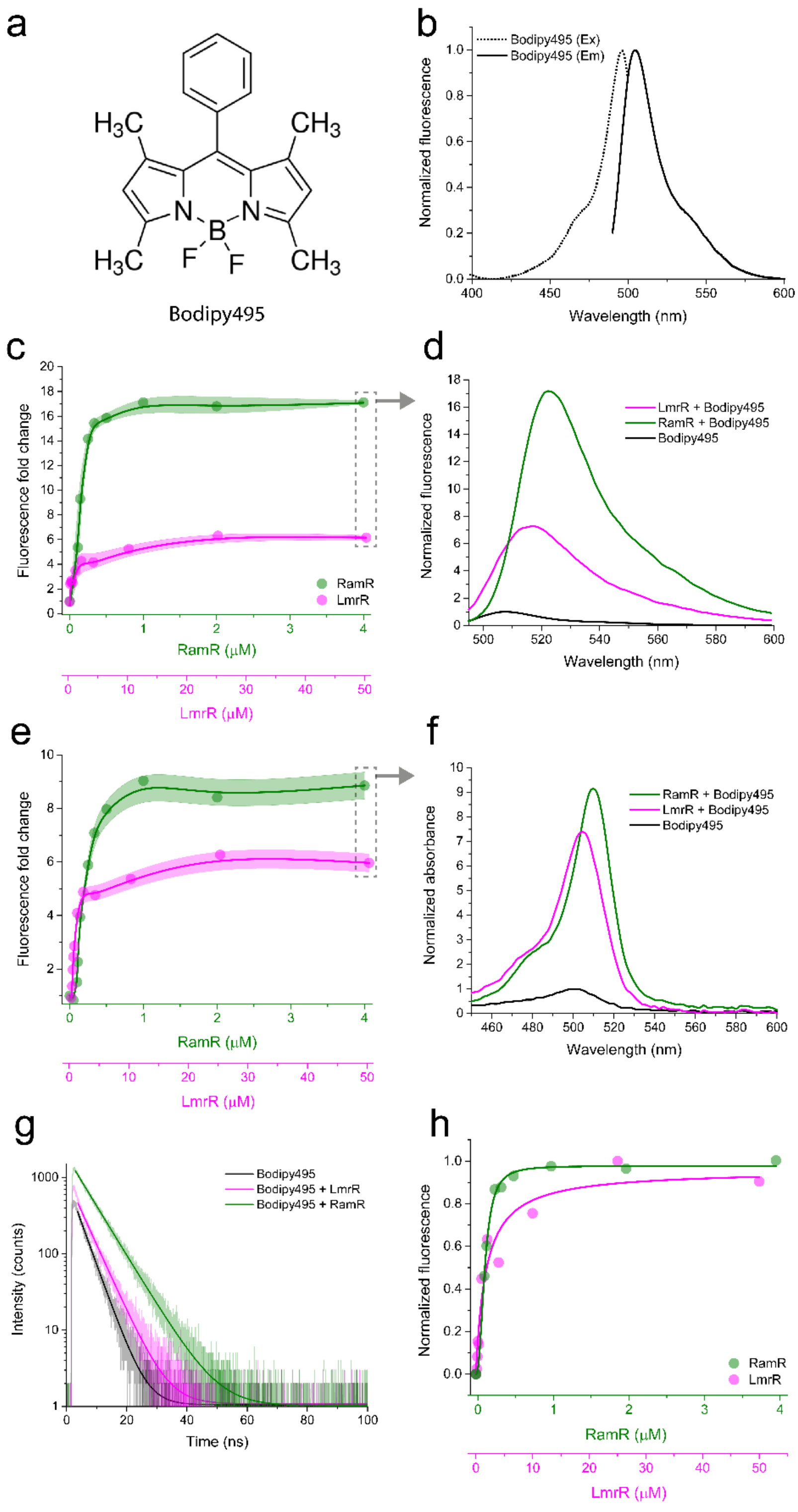


**Supplementary Figure 3**: Characterization of OICP tags (RamR: green and LmrR: magenta) with Bodipy495. (a) Structure of Bodipy495. (b) Excitation (dotted line) and emission (solid line) spectra of Bodipy495. (c) Fluorescence fold change of Bodipy495 on titration with OICP tags. (d) Fluorescence emission spectra of Bodipy495 with RamR and LmrR at a protein:dye molar ratio of 4:1 and 50:1 respectively. (e) Absorbance fold-change of Bodipy495 on titration with OICP tags. Solid lines represent spline fits and shaded regions represent s.d. over three independent measurements in panel (c) and (e). (f) Absorption spectra of Bodipy495 with RamR and LmrR at a protein:dye molar ratio of 4:1 and 50:1 respectively. (g) Fluorescence lifetime spectra of Bodipy495 with RamR and LmrR at a protein:dye molar ratio of 4:1 and 50:1 respectively fit with a mono-exponential decay function (solid lines). (h) Bound fraction of Bodipy495 with OICP tags ascertained from a Hill fit. All experiments were performed at 30 $^{o}$C in 20 mM K-MOPS, 150 mM NaCl buffered at pH 7.0.

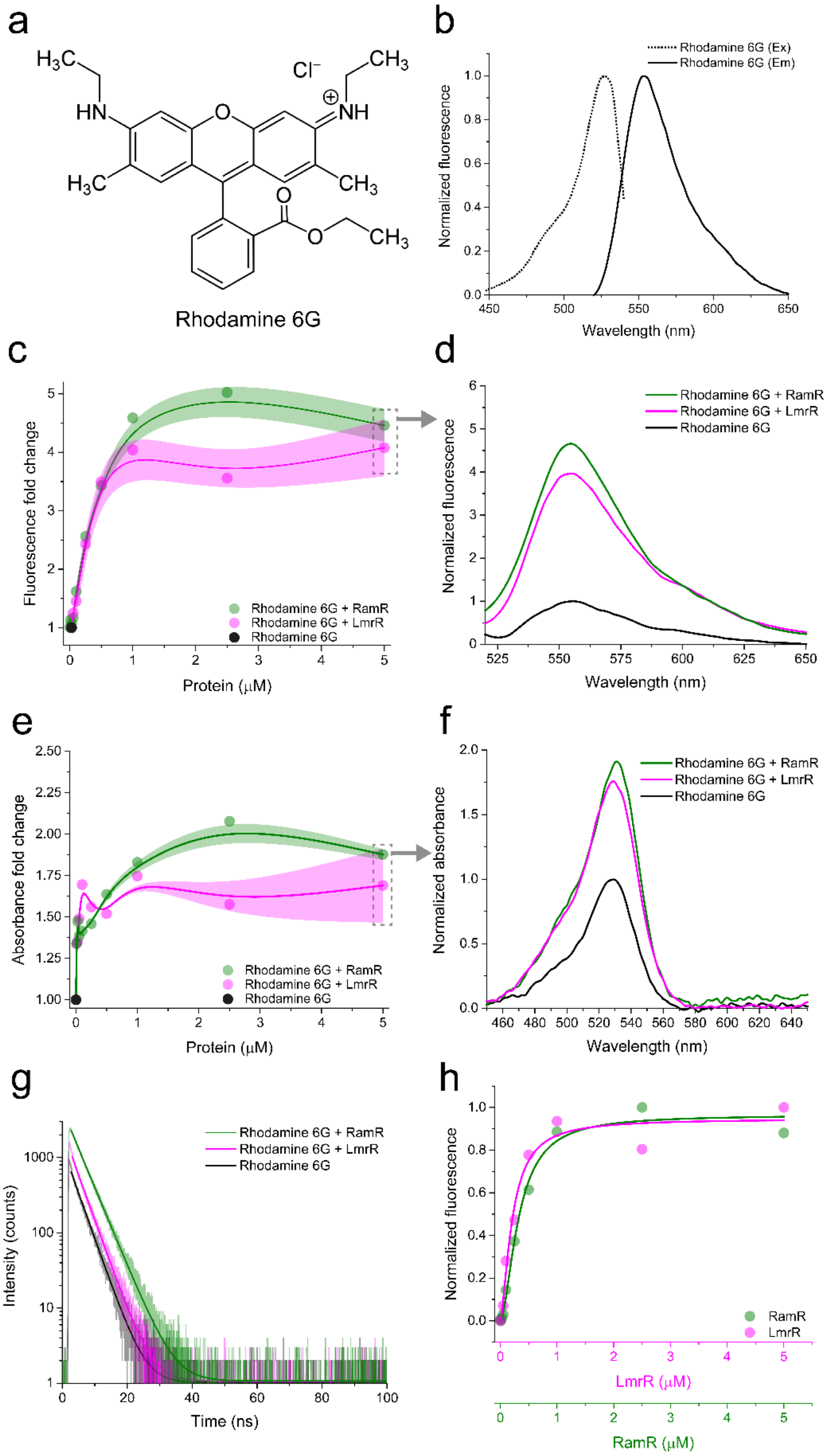


**Supplementary Figure 4**: Characterization of OICP tags (RamR: green and LmrR: magenta) with Rhodamine 6G. (a) Structure of Rhodamine 6G. (b) Excitation (dotted line) and emission (solid line) spectra of Rhodamine 6G. (c) Fluorescence fold change of Rhodamine 6G on titration with OICP tags. (d) Fluorescence emission spectra of Rhodamine 6G with OICP tags at a protein:dye molar ratio of 5:1. (e) Absorbance fold-change of Rhodamine 6G on titration with OICP tags. Solid lines represent spline fits and shaded regions represent s.d. over three independent measurements in panel (c) and (e). (f) Absorption spectra of Rhodamine 6G with OICP tags at a protein:dye molar ratio of 5:1. (g) Fluorescence lifetime spectra of Rhodamine 6G with OICP tags at a protein:dye molar ratio of 5:1 fit with a mono-exponential decay function (solid lines). (h) Bound fraction of Rhodamine 6G with OICP tags ascertained from a Hill fit. All experiments were performed at 30 °C in 20 mM K-MOPS, 150 mM NaCl buffered at pH 7.0.

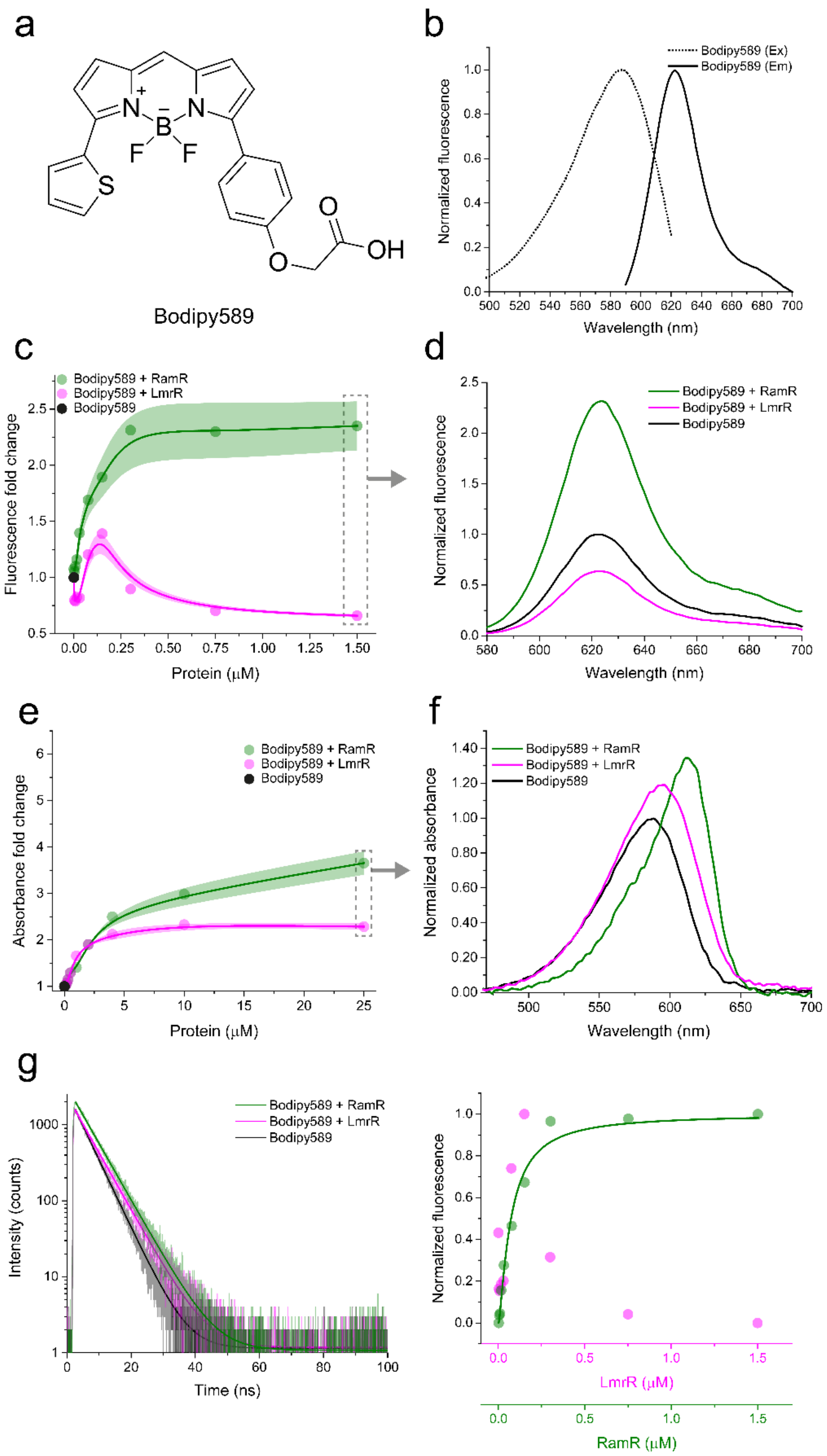


**Supplementary Figure 5**: Characterization of OICP tags (RamR: green and LmrR: magenta) with Bodipy589. (a) Structure of Bodipy589. (b) Excitation (dotted line) and emission (solid line) spectra of Bodipy589. (c) Fluorescence fold change of Bodipy589 on titration with OICP tags. (d) Fluorescence emission spectra of Bodipy589 with OICP tags at a protein:dye molar ratio of 1.5:1. (e) Absorbance fold-change of Bodipy589 on titration with OICP tags. Solid lines represent spline fits and shaded regions represent s.d. over three independent measurements in panel (c) and (e). (f) Absorption spectra of Bodipy589 with OICP tags at a protein:dye molar ratio of 1.5:1. (g) Fluorescence lifetime spectra of Bodipy589 with OICP tags at a protein:dye molar ratio of 1.5:1 fit with a mono-exponential decay function (solid lines). (h) Bound fraction of Bodipy589 with OICP tags ascertained from a Hill fit. All experiments were performed at 30 °C in 20 mM K-MOPS, 150 mM NaCl buffered at pH 7.0.

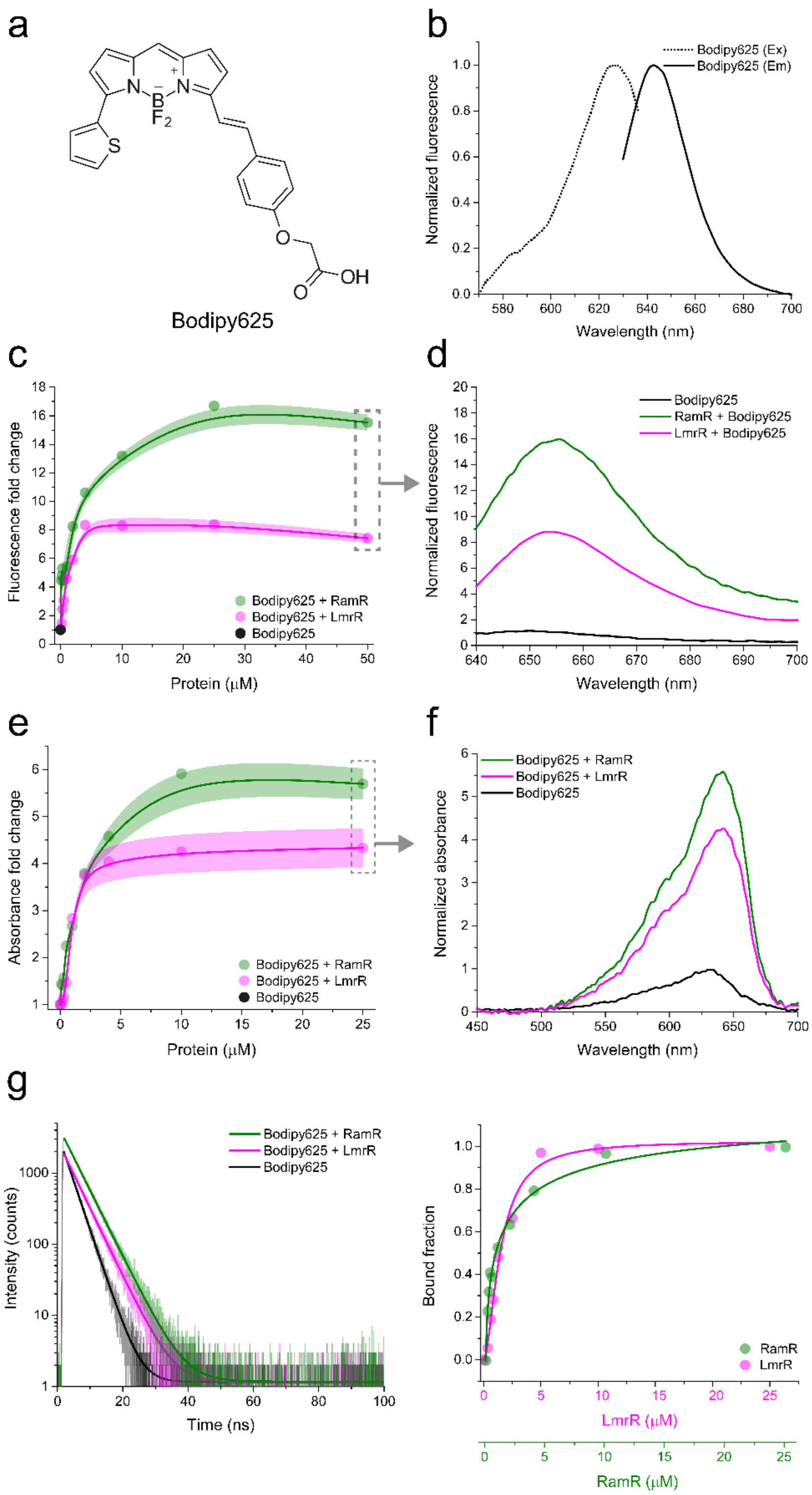


**Supplementary Figure 6**: Characterization of OICP tags (RamR: green and LmrR: magenta) with Bodipy625. (a) Structure of Bodipy625. (b) Excitation (dotted line) and emission (solid line) spectra of Bodipy625. (c) Fluorescence fold change of Bodipy625 on titration with OICP tags. (d) Fluorescence emission spectra of Bodipy625 with OICP tags at a protein:dye molar ratio of 50:1. (e) Absorbance fold-change of Bodipy625 on titration with OICP tags. Solid lines represent spline fits and shaded regions represent s.d. over three independent measurements in panel (c) and (e). (f) Absorption spectra of Bodipy625 with OICP tags at a protein:dye molar ratio of 25:1. (g) Fluorescence lifetime spectra of Bodipy625 with OICP tags at a protein:dye molar ratio of 25:1 fit with a mono-exponential decay function (solid lines). (h) Bound fraction of Bodipy625 with OICP tags ascertained from a Hill fit. All experiments were performed at 30 $^{o}$C in 20 mM K-MOPS, 150 mM NaCl buffered at pH 7.0.

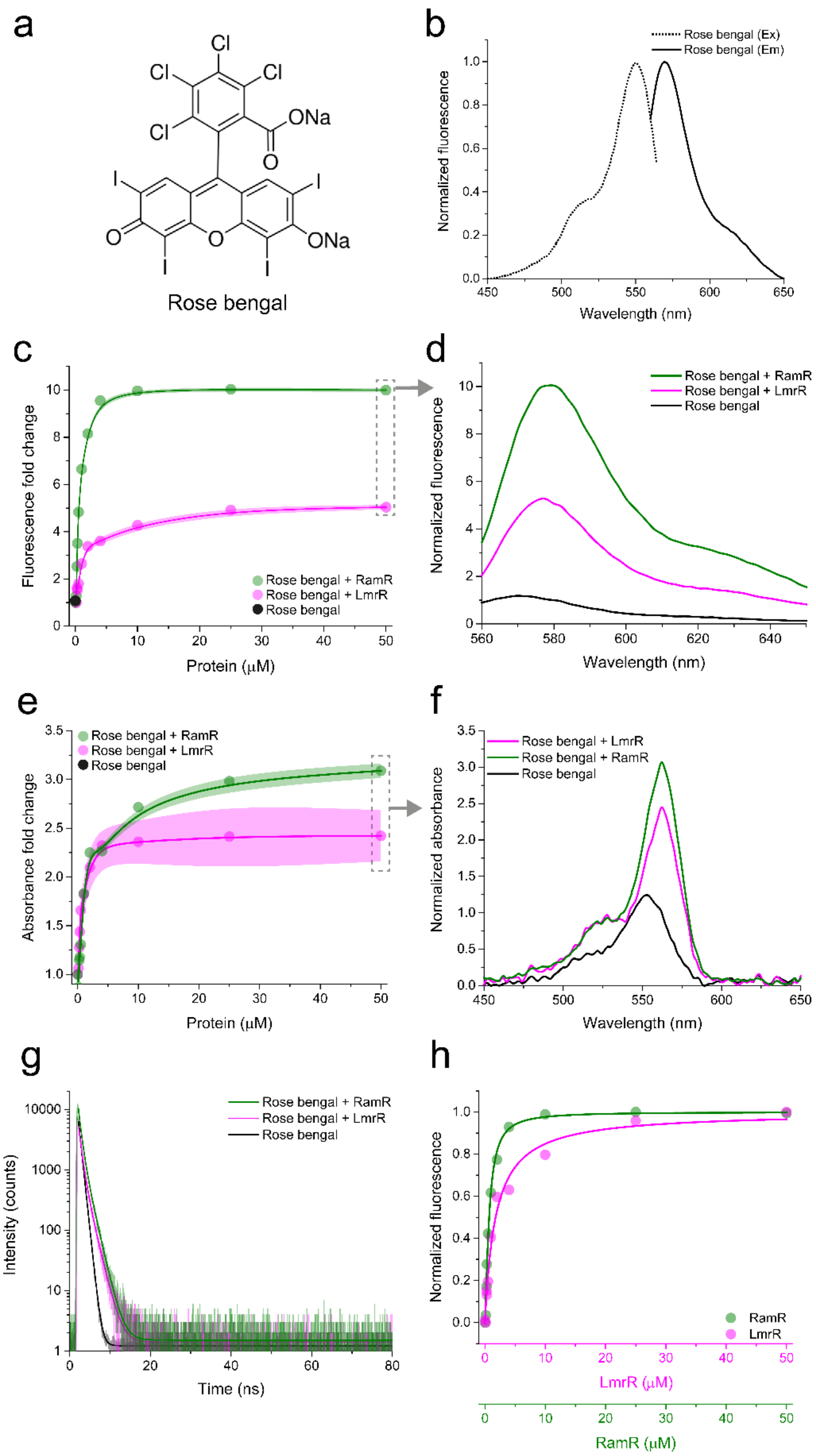


**Supplementary Figure 7**: Characterization of OICP tags (RamR: green and LmrR: magenta) with Rose bengal. (a) Structure of Rose bengal. (b) Excitation (dotted line) and emission (solid line) spectra of Rose bengal. (c) Fluorescence fold change of Rose bengal on titration with OICP tags. (d) Fluorescence emission spectra of Rose bengal with OICP tags at a protein:dye molar ratio of 50:1. (e) Absorbance fold-change of Rose bengal on titration with OICP tags. Solid lines represent spline fits and shaded regions represent s.d. over three independent measurements in panel (c) and (e). (f) Absorption spectra of Rose bengal with OICP tags at a protein:dye molar ratio of 50:1. (g) Fluorescence lifetime spectra of Rose bengal with OICP tags at a protein:dye molar ratio of 50:1 fit with a mono-exponential decay function (solid lines). (h) Bound fraction of Rose bengal with OICP tags ascertained from a Hill fit. All experiments were performed at 30 $^{o}$C in 20 mM K-MOPS, 150 mM NaCl buffered at pH 7.0.

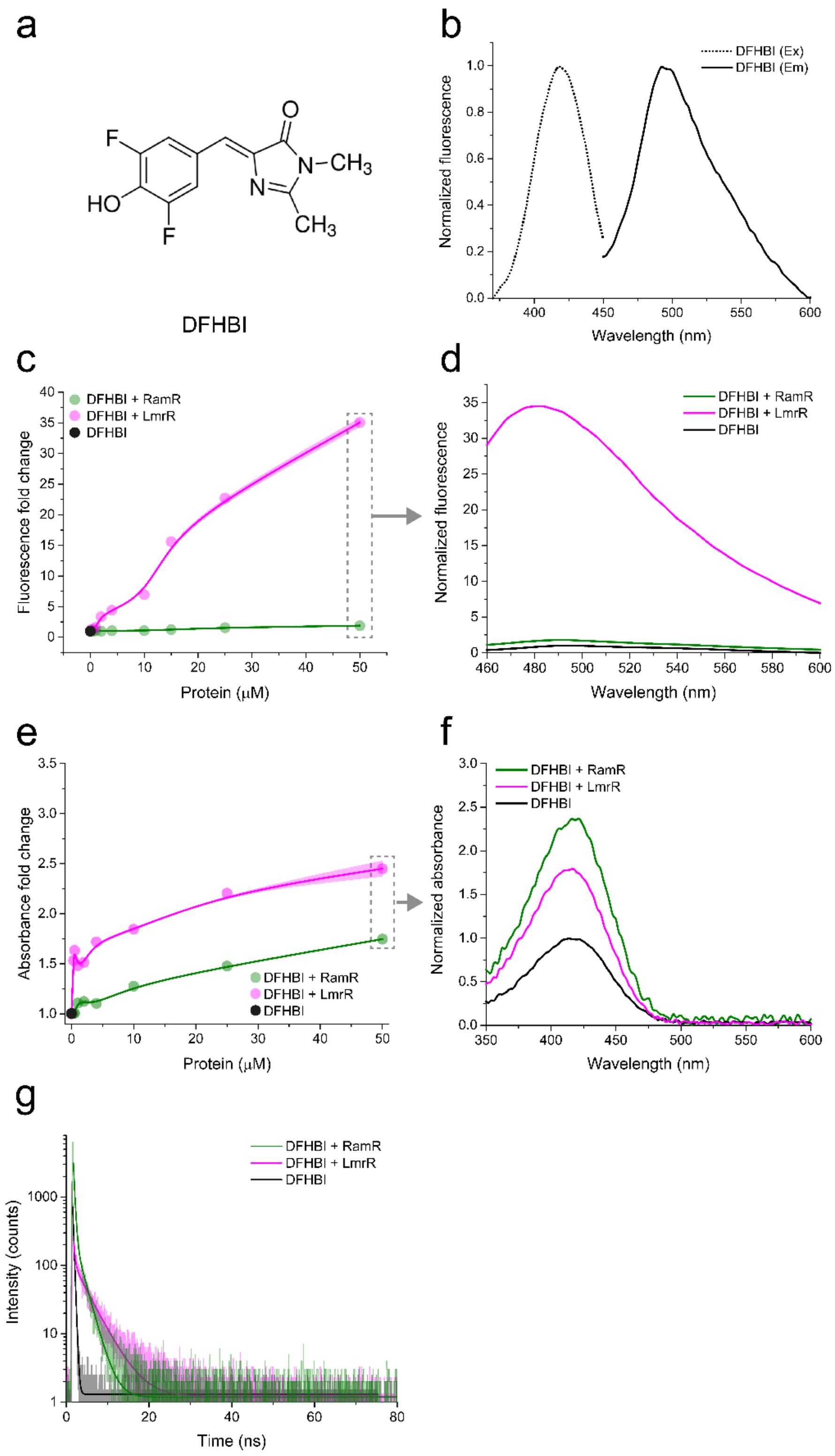


**Supplementary Figure 8**: Characterization of OICP tags (RamR: green and LmrR: magenta) with DFHBI. (a) Structure of DFHBI. (b) Excitation (dotted line) and emission (solid line) spectra of DFHBI. (c) Fluorescence fold change of DFHBI on titration with OICP tags. (d) Fluorescence emission spectra of DFHBI with OICP tags at a protein:dye molar ratio of 50:1. (e) Absorbance fold-change of DFHBI on titration with OICP tags. Solid lines represent spline fits and shaded regions represent s.d. over three independent measurements in panel (c) and (e). (f) Absorption spectra of DFHBI with OICP tags at a protein:dye molar ratio of 50:1. (g) Fluorescence lifetime spectra of DFHBI with OICP tags at a protein:dye molar ratio of 50:1 fit with a bi-exponential decay function (solid lines). All experiments were performed at 30 $^{o}$C in 20 mM K-MOPS, 150 mM NaCl buffered at pH 7.0.

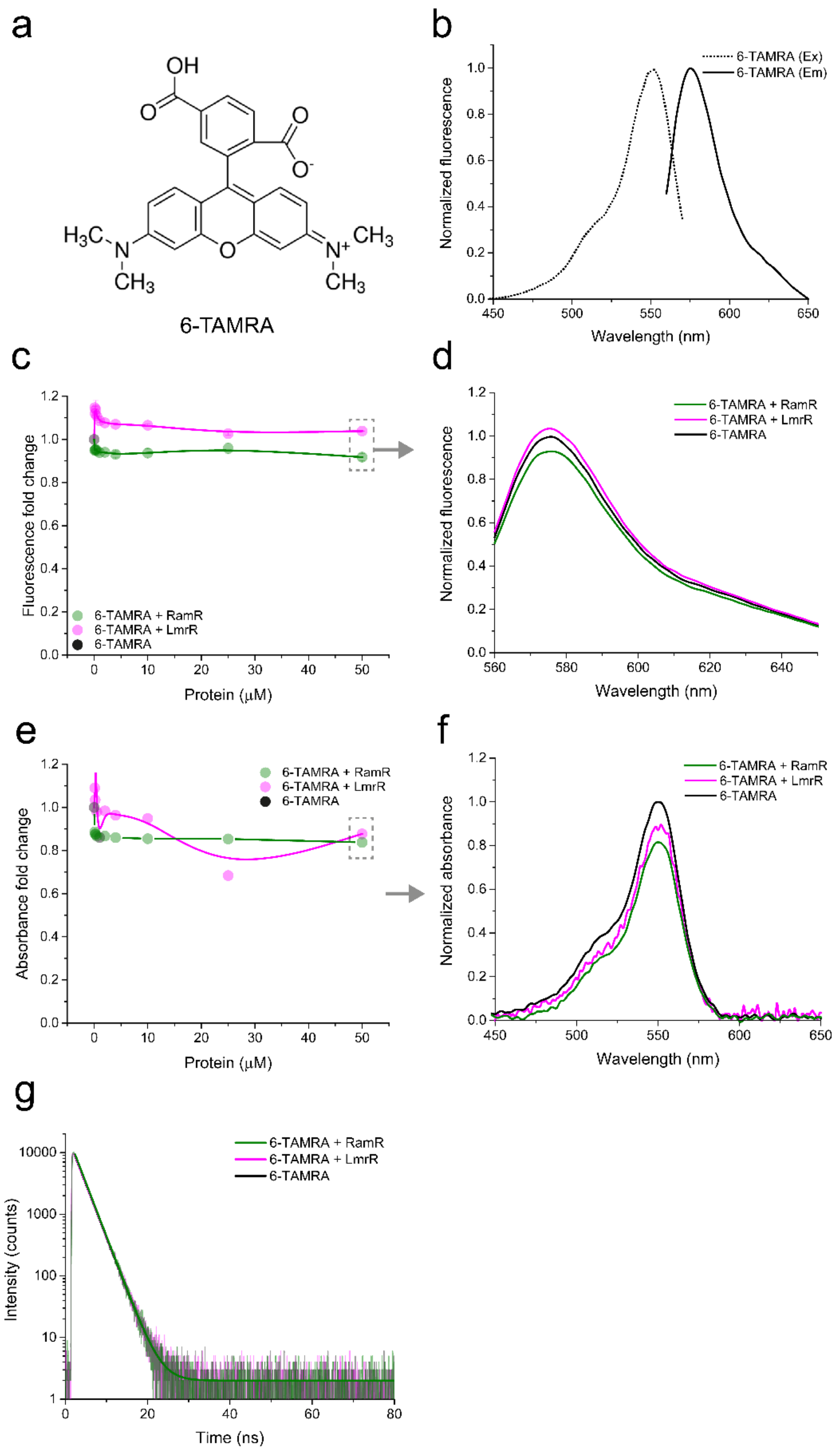


**Supplementary Figure 9**: Characterization of OICP tags (RamR: green and LmrR: magenta) with 6-TAMRA. (a) Structure of 6-TAMRA. (b) Excitation (dotted line) and emission (solid line) spectra of 6-TAMRA. (c) Fluorescence fold change of 6-TAMRA on titration with OICP tags. (d) Fluorescence emission spectra of 6-TAMRA with OICP tags at a protein:dye molar ratio of 50:1. (e) Absorbance fold-change of 6-TAMRA on titration with OICP tags. Solid lines represent spline fits and shaded regions represent s.d. over three independent measurements in panel (c) and (e). (f) Absorption spectra of 6-TAMRA with OICP tags at a protein:dye molar ratio of 50:1. (g) Fluorescence lifetime spectra of 6-TAMRA with OICP tags at a protein:dye molar ratio of 50:1 fit with a mono-exponential decay function (solid lines). All experiments were performed at 30 $^{o}$C in 20 mM K-MOPS, 150 mM NaCl buffered at pH 7.0.

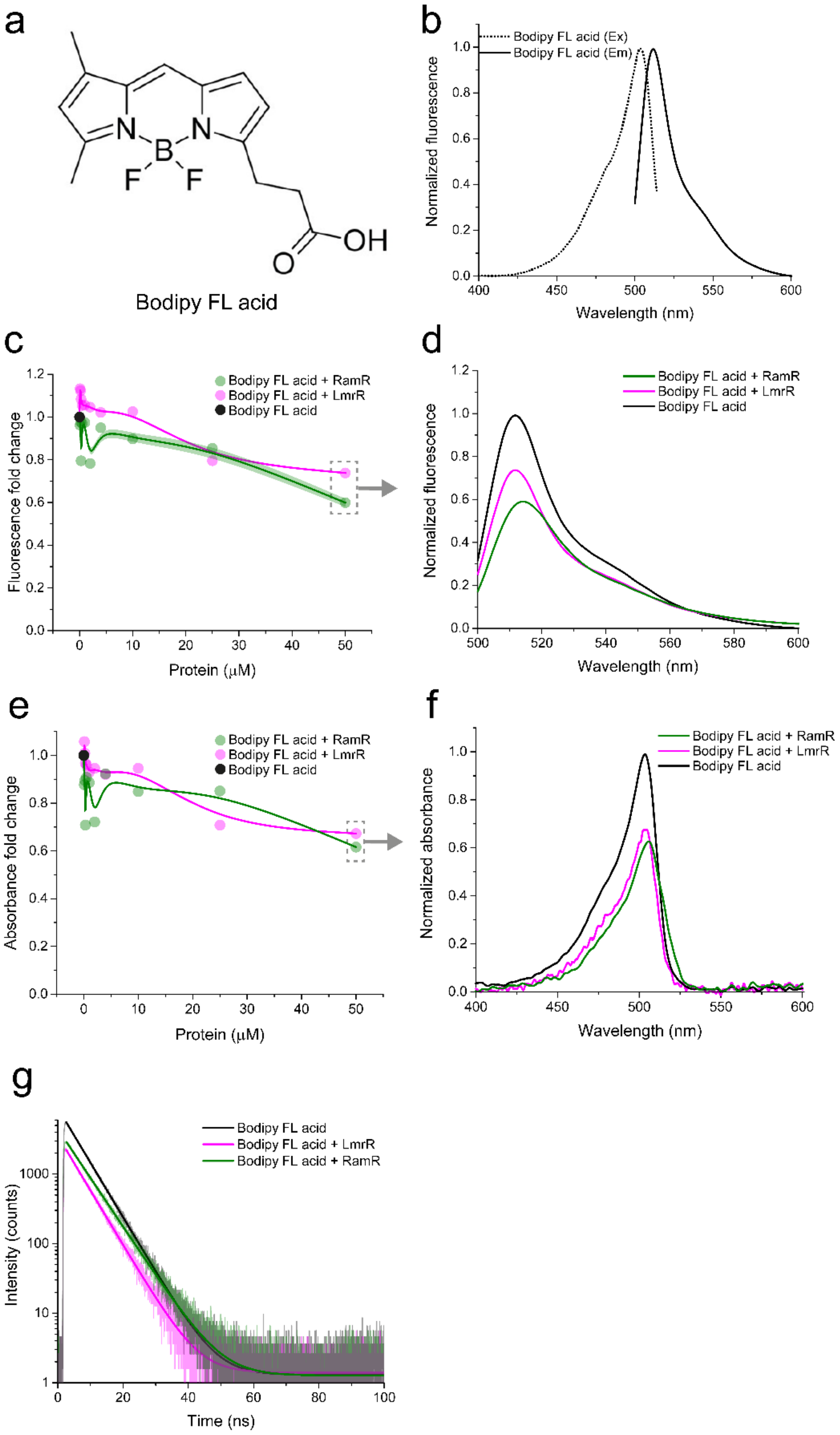


**Supplementary Figure 10**: Characterization of OICP tags (RamR: green and LmrR: magenta) with Bodipy FL acid. (a) Structure of Bodipy FL acid. (b) Excitation (dotted line) and emission (solid line) spectra of Bodipy FL acid. (c) Fluorescence fold change of Bodipy FL acid on titration with OICP tags. (d) Fluorescence emission spectra of Bodipy FL acid with OICP tags at a protein:dye molar ratio of 50:1. (e) Absorbance fold-change of Bodipy FL acid on titration with OICP tags. Solid lines represent spline fits and shaded regions represent s.d. over three independent measurements in panel (c) and (e). (f) Absorption spectra of Bodipy FL acid with OICP tags at a protein:dye molar ratio of 50:1. (g) Fluorescence lifetime spectra of Bodipy FL acid with OICP tags at a protein:dye molar ratio of 50:1 fit with a mono-exponential decay function (solid lines). All experiments were performed at 30 $^{o}$C in 20 mM K-MOPS, 150 mM NaCl buffered at pH 7.0.

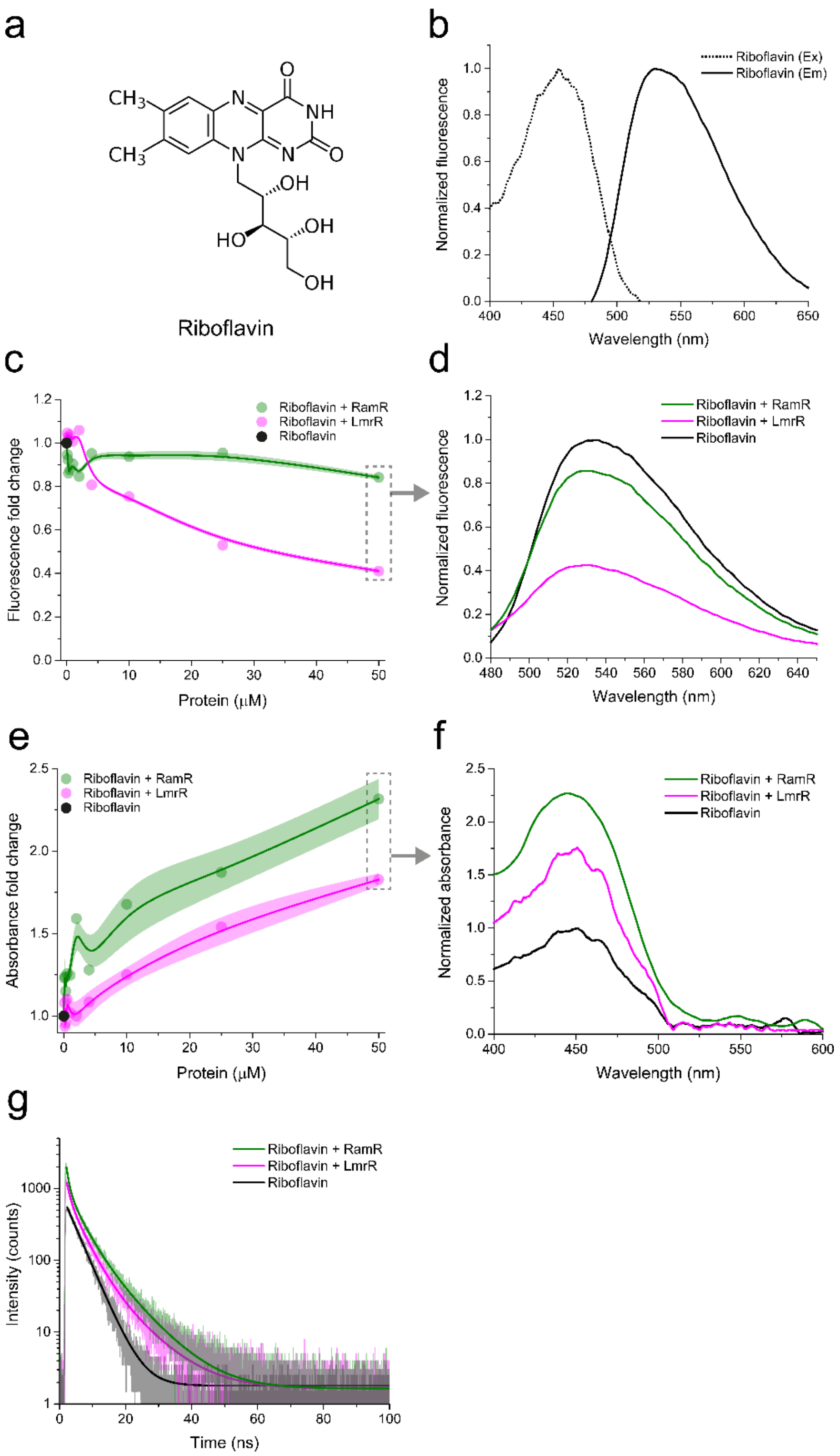


**Supplementary Figure 11**: Characterization of OICP tags (RamR: green and LmrR: magenta) with Riboflavin. (a) Structure of Riboflavin. (b) Excitation (dotted line) and emission (solid line) spectra of Riboflavin. (c) Fluorescence fold change of Riboflavin on titration with OICP tags. (d) Fluorescence emission spectra of Riboflavin with OICP tags at a protein:dye molar ratio of 50:1. (e) Absorbance fold-change of Riboflavin on titration with OICP tags. Solid lines represent spline fits and shaded regions represent s.d. over three independent measurements in panel c) and e). (f) Absorption spectra of Riboflavin with OICP tags at a protein:dye molar ratio of 50:1. (g) Fluorescence lifetime spectra of Riboflavin with OICP tags at a protein:dye molar ratio of 50:1 fit with a bi-exponential decay function (solid lines). All experiments were performed at 30 °C in 20 mM K-MOPS, 150 mM NaCl buffered at pH 7.0.

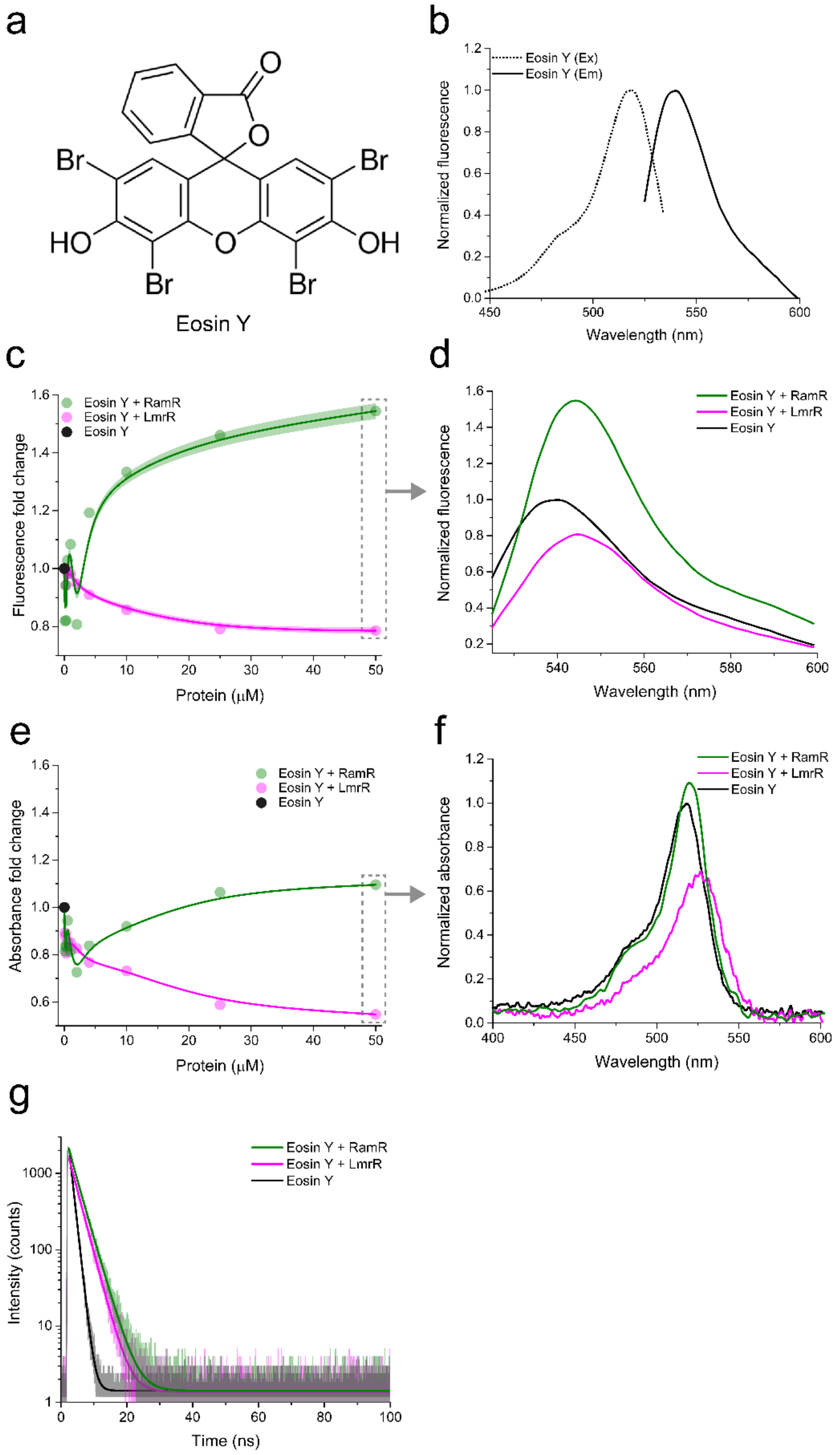


**Supplementary Figure 12**: Characterization of OICP tags (RamR: green and LmrR: magenta) with Eosin Y. (a) Structure of Eosin Y. (b) Excitation (dotted line) and emission (solid line) spectra of Eosin Y. (c) Fluorescence fold change of Eosin Y on titration with OICP tags. (d) Fluorescence emission spectra of Eosin Y with OICP tags at a protein:dye molar ratio of 50:1. (e) Absorbance fold-change of Eosin Y on titration with OICP tags. Solid lines represent spline fits and shaded regions represent s.d. over three independent measurements in panel c) and e). (f) Absorption spectra of Eosin Y with OICP tags at a protein:dye molar ratio of 50:1. (g) Fluorescence lifetime spectra of Eosin Y with OICP tags at a protein:dye molar ratio of 50:1 fit with a mono-exponential decay function (solid lines). All experiments were performed at 30 °C in 20 mM K-MOPS, 150 mM NaCl buffered at pH 7.0.

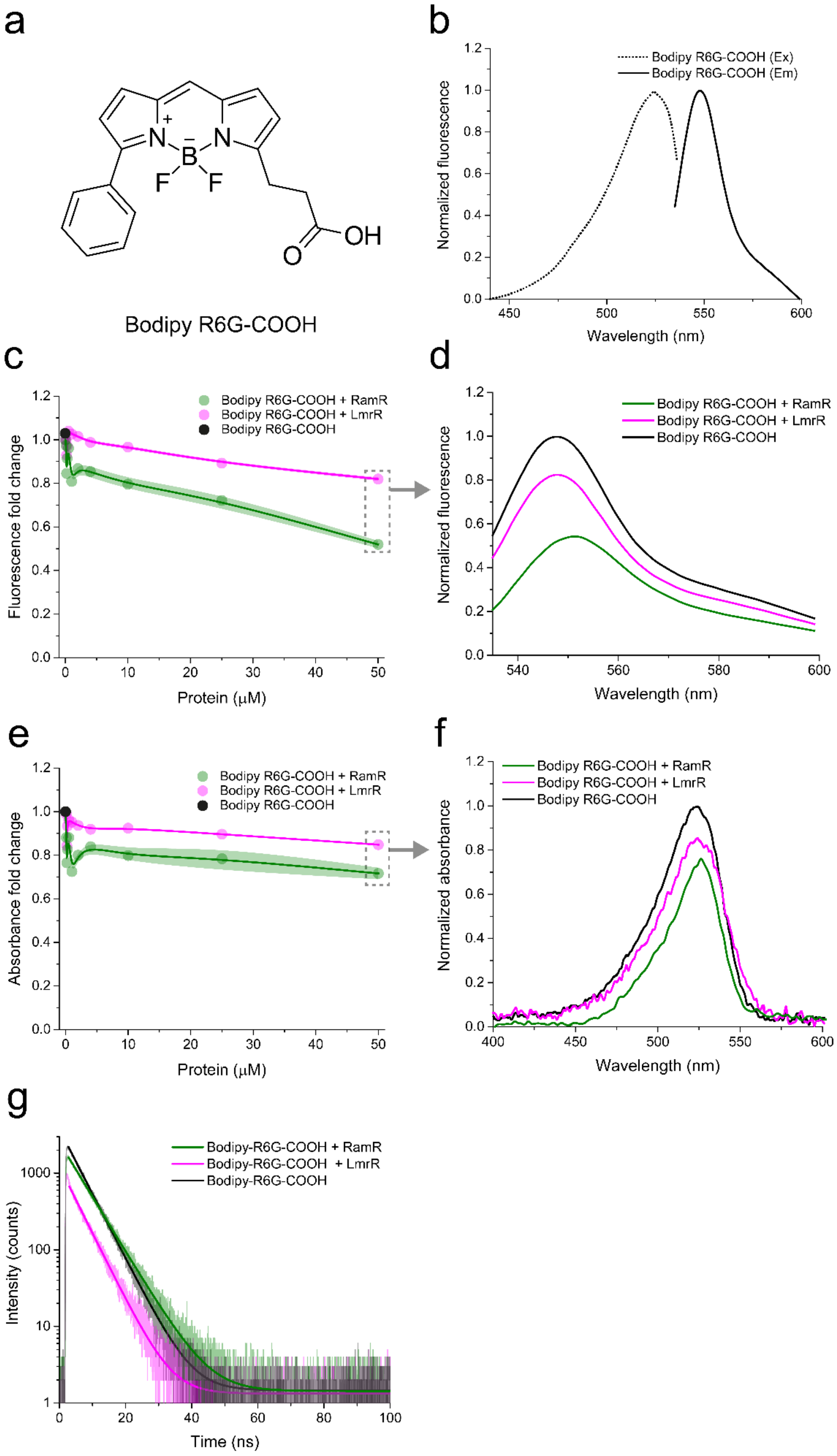


**Supplementary Figure 13**: Characterization of OICP tags (RamR: green and LmrR: magenta) with Bodipy R6G. (a) Structure of Bodipy R6G. (b) Excitation (dotted line) and emission (solid line) spectra of Bodipy R6G. (c) Fluorescence fold change of Bodipy R6G on titration with OICP tags. (d) Fluorescence emission spectra of Bodipy R6G with OICP tags at a protein:dye molar ratio of 50:1. (e) Absorbance fold-change of Bodipy R6G on titration with OICP tags. Shaded regions in panel c) and e) represent s.d. over three independent measurements. (f) Absorption spectra of Bodipy R6G with OICP tags at a protein:dye molar ratio of 50:1. (g) Fluorescence lifetime spectra of Bodipy R6G with OICP tags at a protein:dye molar ratio of 50:1 fit with a mono-exponential decay function (solid lines). All experiments were performed at 30 $^{o}$C in 20 mM K-MOPS, 150 mM NaCl buffered at pH 7.0.

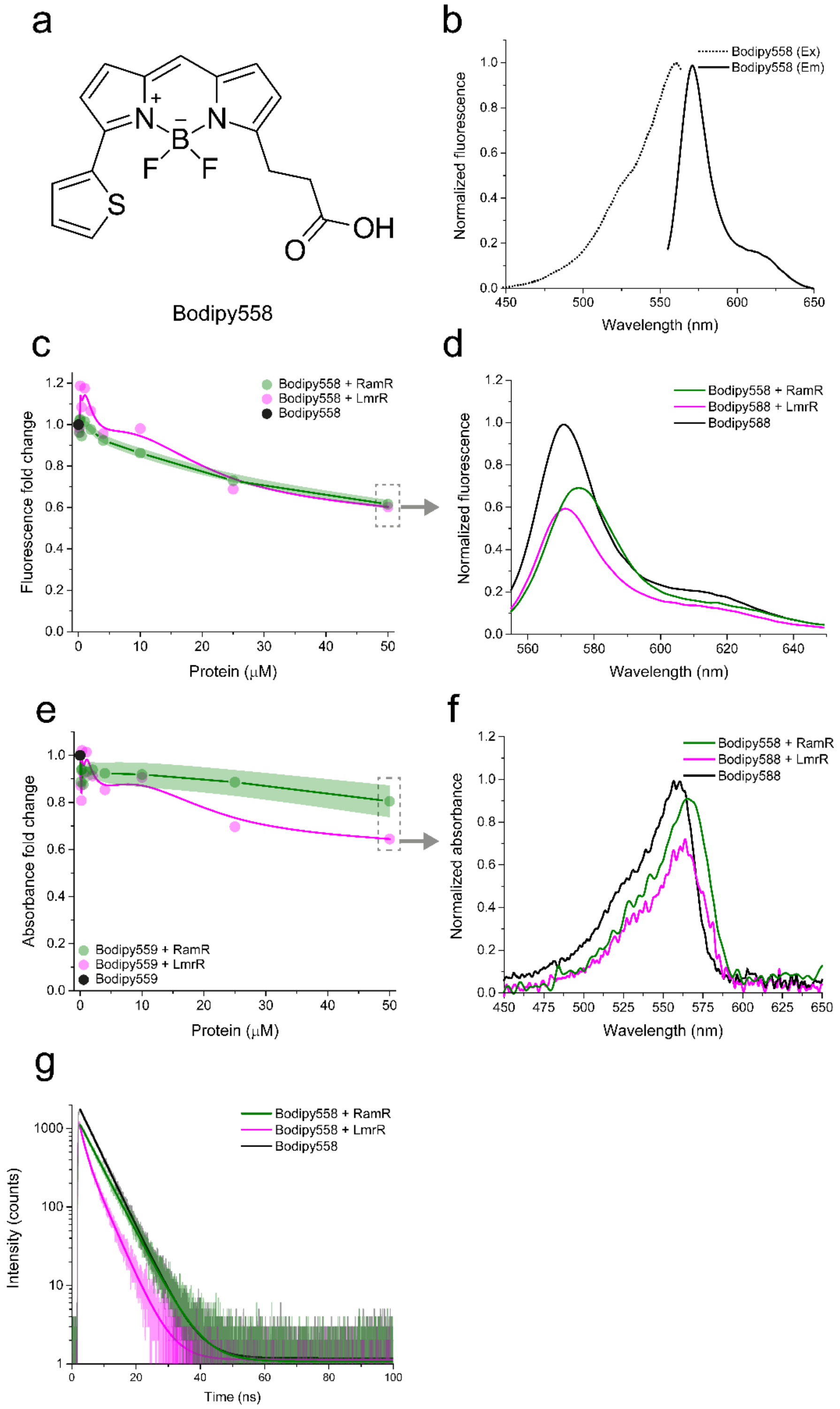


**Supplementary Figure 14**: Characterization of OICP tags (RamR: green and LmrR: magenta) with Bodipy558. (a) Structure of Bodipy558. (b) Excitation (dotted line) and emission (solid line) spectra of Bodipy558. (c) Fluorescence fold change of Bodipy558 on titration with OICP tags. (d) Fluorescence emission spectra of Bodipy558 with OICP tags at a protein:dye molar ratio of 50:1. (e) Absorbance fold-change of Bodipy558 on titration with OICP tags. Shaded regions in panel c) and e) represent s.d. over three independent measurements. (f) Absorption spectra of Bodipy558 with OICP tags at a protein:dye molar ratio of 50:1. (g) Fluorescence lifetime spectra of Bodipy558 with OICP tags at a protein:dye molar ratio of 50:1 fit with a mono-exponential decay function (solid lines). All experiments were performed at 30 °C in 20 mM K-MOPS, 150 mM NaCl buffered at pH 7.0.

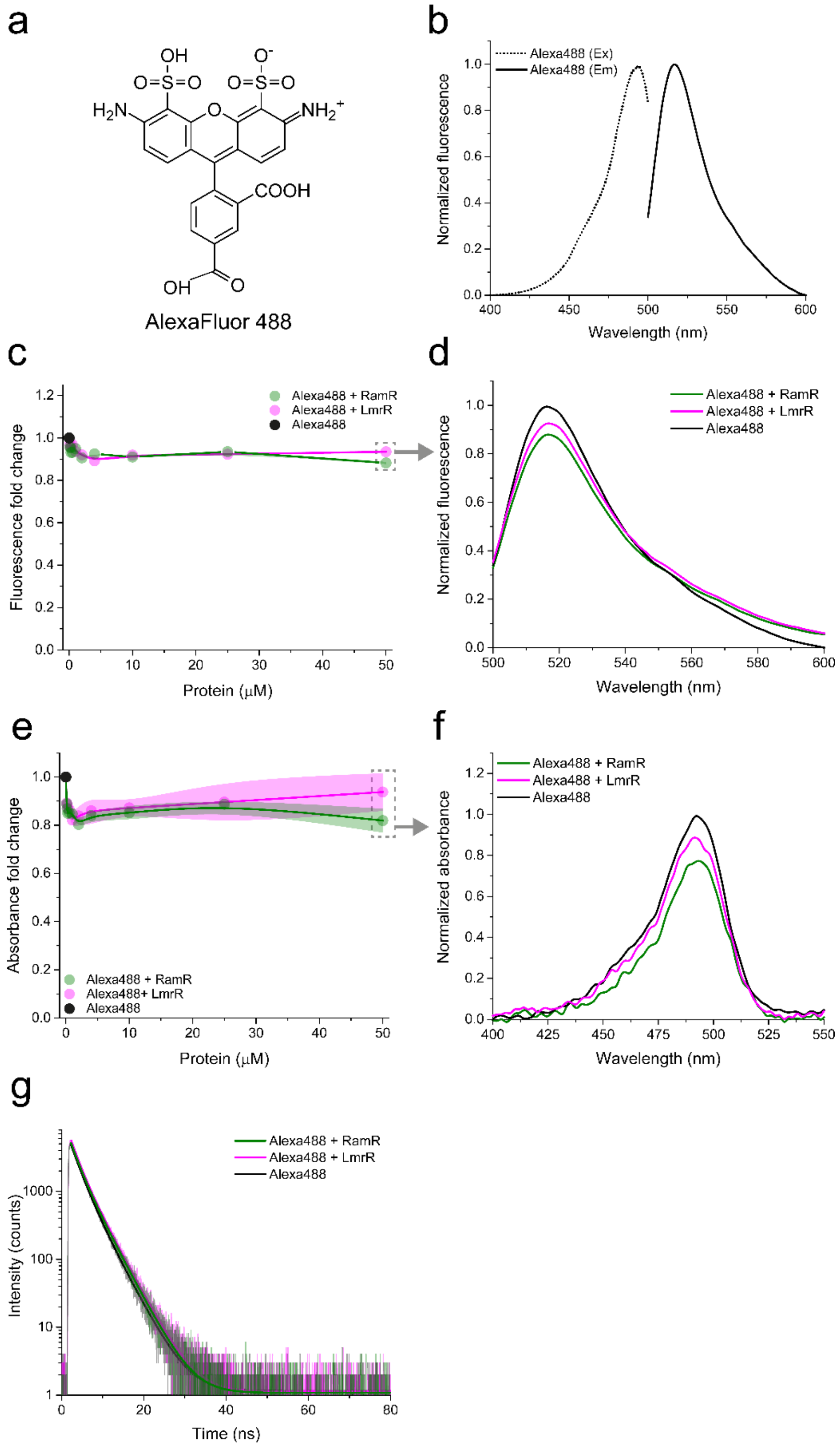


**Supplementary Figure 15**: Characterization of OICP tags (RamR: green and LmrR: magenta) with Alexa488. (a) Structure of Alexa488. (b) Excitation (dotted line) and emission (solid line) spectra of Alexa488. (c) Fluorescence fold change of Alexa488 on titration with OICP tags. (d) Fluorescence emission spectra of Alexa488 with OICP tags at a protein:dye molar ratio of 50:1. (e) Absorbance fold-change of Alexa488 on titration with OICP tags. Shaded regions in panel c) and e) represent s.d. over three independent measurements. (f) Absorption spectra of Alexa488 with OICP tags at a protein:dye molar ratio of 50:1. (g) Fluorescence lifetime spectra of Alexa488 with OICP tags at a protein:dye molar ratio of 50:1 fit with a mono-exponential decay function (solid lines). All experiments were performed at 30 °C in 20 mM K-MOPS, 150 mM NaCl buffered at pH 7.0.

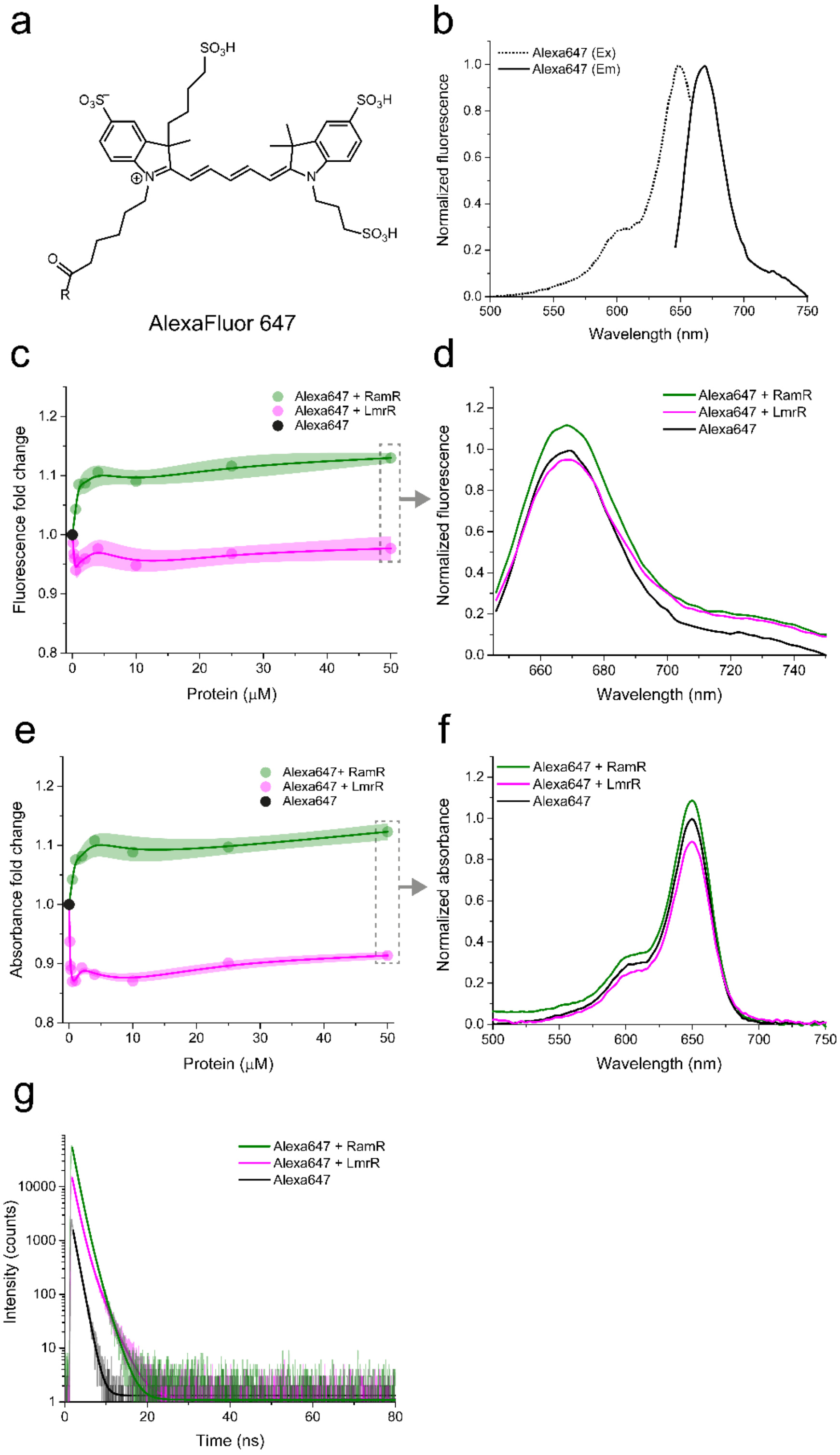


**Supplementary Figure 16**: Characterization of OICP tags (RamR: green and LmrR: magenta) with Alexa647. (a) Structure of Alexa647. (b) Excitation (dotted line) and emission (solid line) spectra of Alexa647. (c) Fluorescence fold change of Alexa647 on titration with OICP tags. (d) Fluorescence emission spectra of Alexa647 with OICP tags at a protein:dye molar ratio of 50:1. (e) Absorbance fold-change of Alexa647 on titration with OICP tags. Shaded regions in panel c) and e) represent s.d. over three independent measurements. (f) Absorption spectra of Alexa647 with OICP tags at a protein:dye molar ratio of 50:1. (g) Fluorescence lifetime spectra of Alexa647 with OICP tags at a protein:dye molar ratio of 50:1 fit with a mono-exponential decay function (solid lines). All experiments were performed at 30 °C in 20 mM K-MOPS, 150 mM NaCl buffered at pH 7.0.

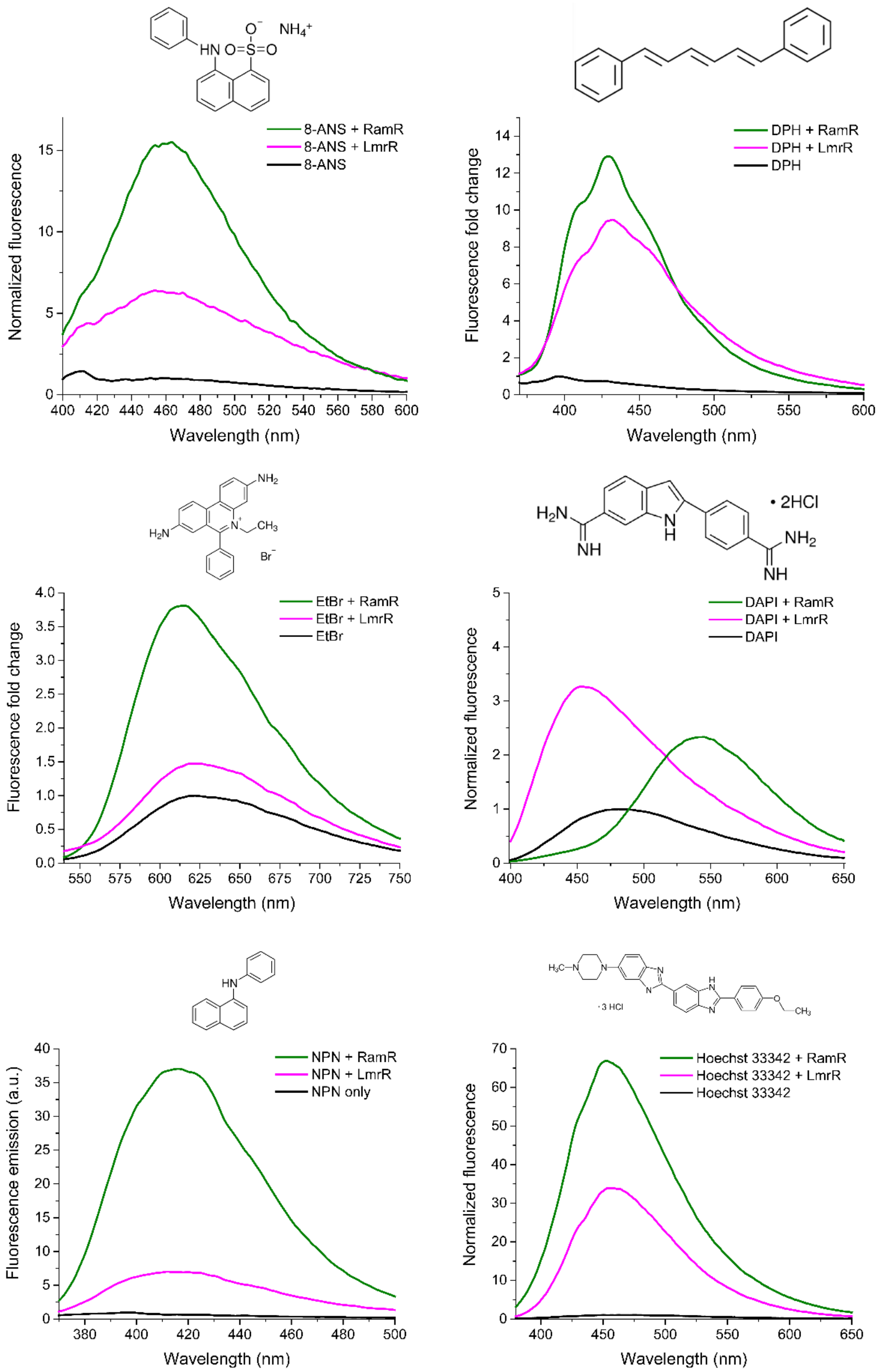


**Supplementary Figure 17**: Fluorescence emission spectra of nonspecific intercalating dyes with OICP tags (RamR: green and LmrR: magenta) with (a) *8*-Anilinonaphthalene-1-sulfonic acid (ANS) (b) 1,6-diphenyl-1,3,5-hexatriene (DPH) (c) ethidium bromide (d) 4′,6-diamidino-2-phenylindole (DAPI) (e) 1-N-phenylnaphthylamine (NPN) and (f) Hoechst 33342 at protein:dye molar ratio of 50:1. The corresponding dye structures are indicated above each panel. All experiments were performed at 30 °C in 20 mM K-MOPS, 150 mM NaCl buffered at pH 7.0.

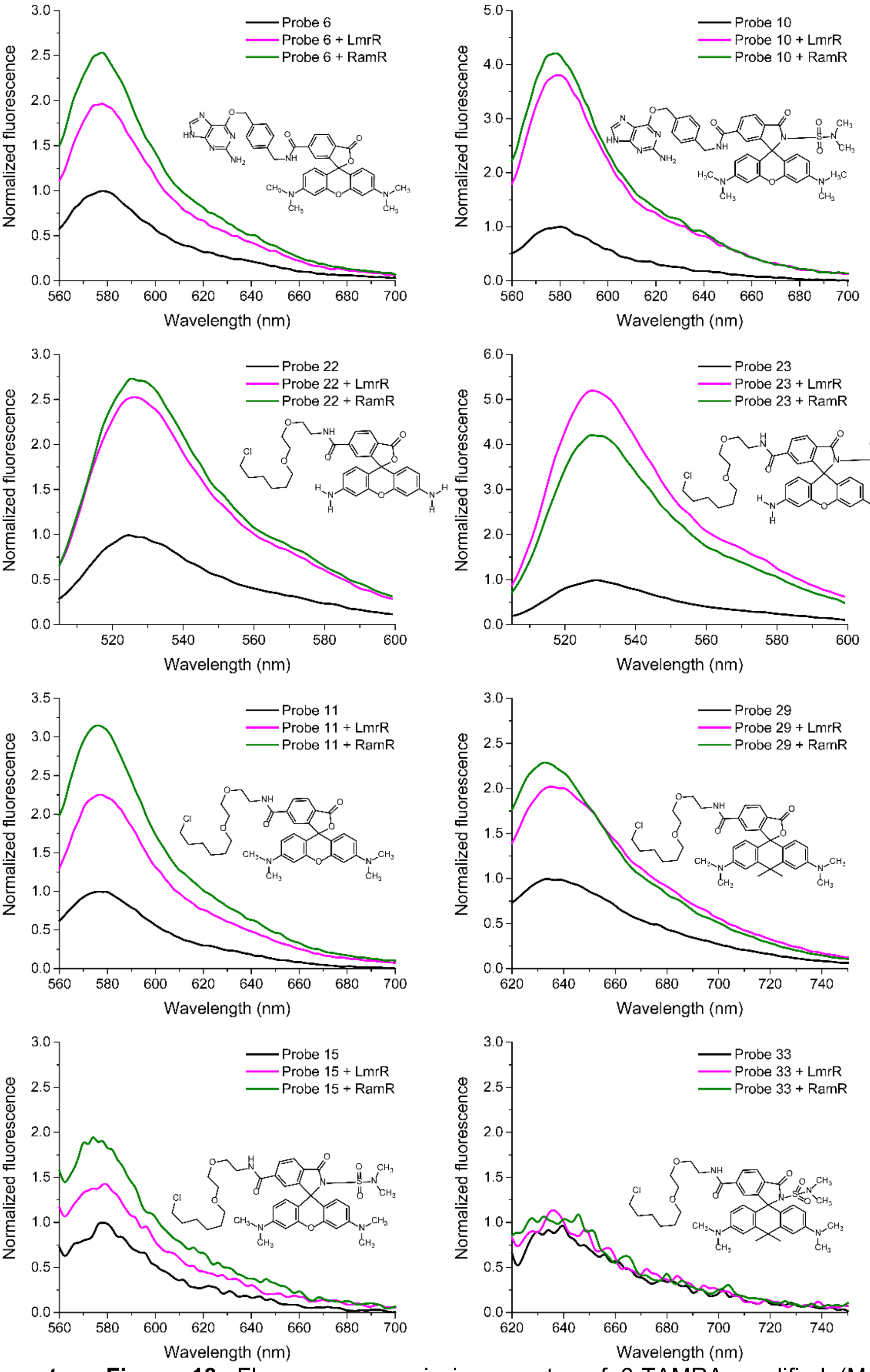


**Supplementary Figure 18**: Fluorescence emission spectra of 6-TAMRA modified (MaP) probes (Wang et al. 2020) with OICP tags at a protein:dye molar ratio of 50:1. MaP probes comprise of 6-TAMRA covalently linked to the SNAP-Tag ligand: $O^6$-benzyl guanidine (Probe 6 and Probe 10) or to the Halo-Tag ligand (Probe 11, 15, 22, 23, 29 and 33). All experiments were performed at 30 $^oC$ in 20 mM K-MOPS, 150 mM NaCl buffered at pH 7.0.

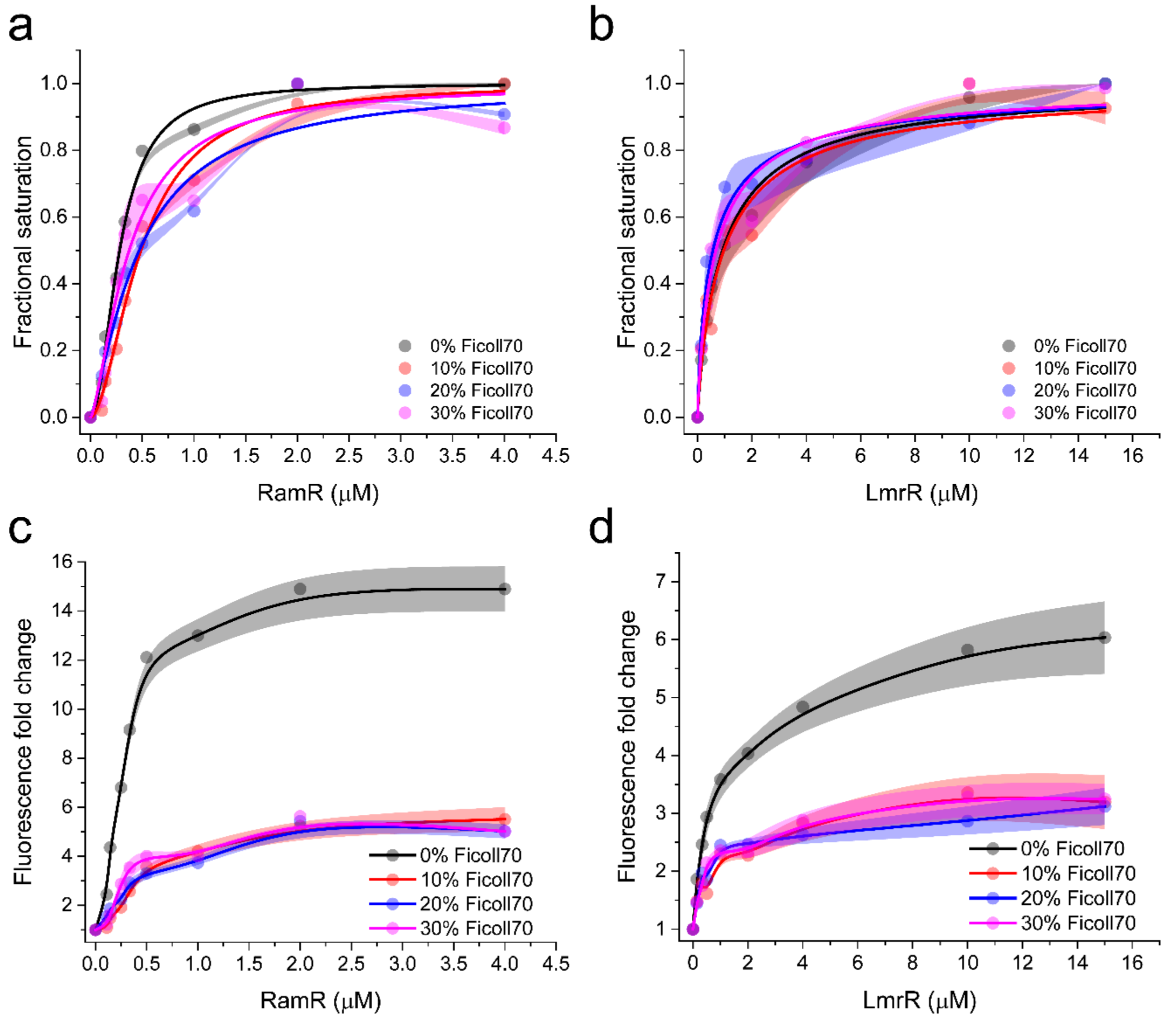


**Supplementary Figure 19**: Effect of Ficoll70 on the fluorescence of Bodipy495 in the presence of OICP tags (a,c) RamR and (b,d) LmrR. Saturation experiments were performed by titrating the respective OICP tag with 1 μM Bodipy495. The fold change in emission fluorescence of Bodipy495 with increasing concentrations of Ficoll70 is shown in panel c and d. All experiments were performed at 30 °C in 20 mM K-MOPS, 150 mM NaCl buffered at pH 7.0.

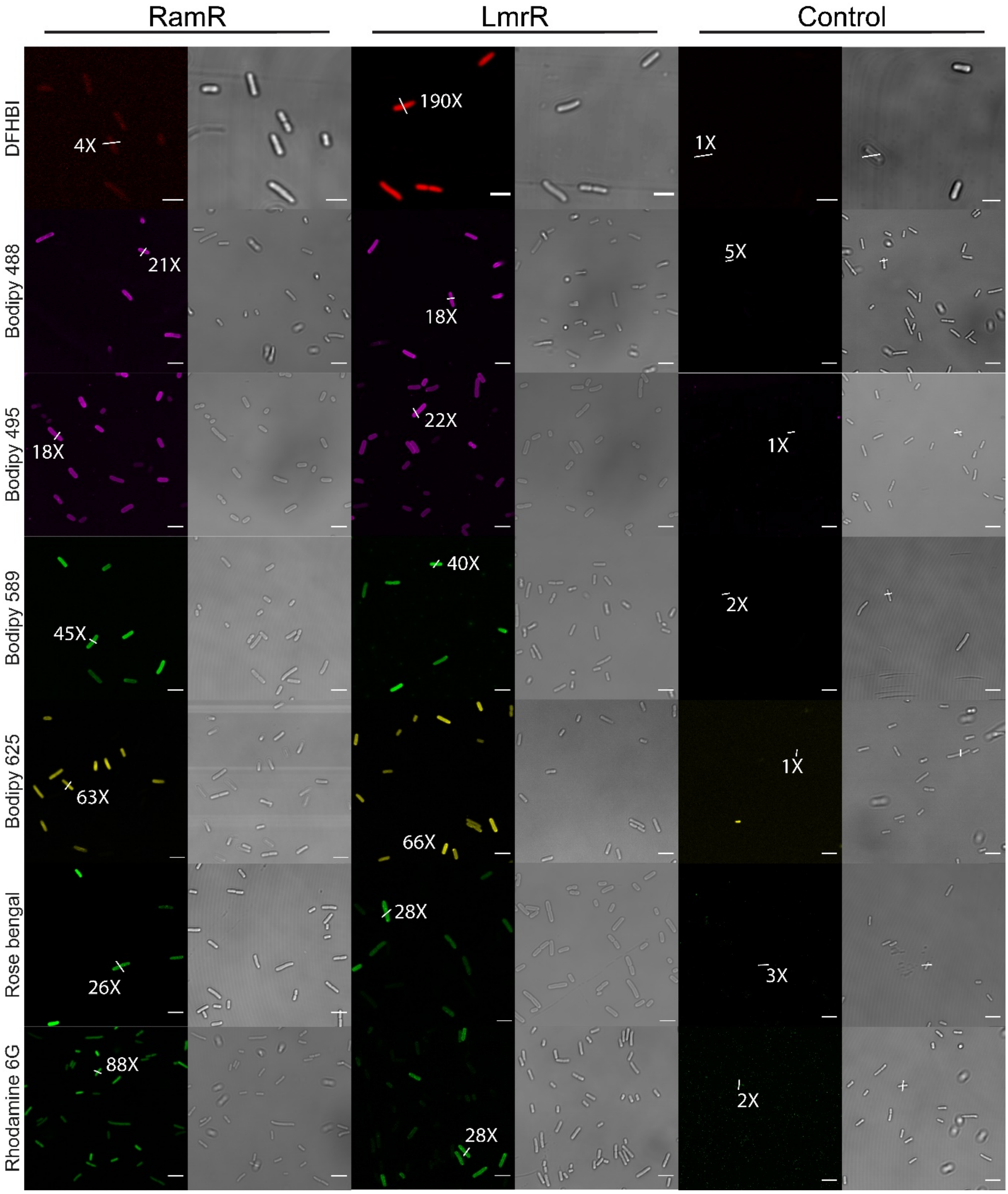


**Supplementary Figure 20**: Overview of labeled *E. coli* cells expressing cytoplasmic OICP tags from a pBAD vector. The signal to background fluorescence ratios for arbitrarily picked cells for each dye is mentioned next to white overlays. All measurements were performed at 30 °C. For controls, the signal to background ratio was minimal for all samples. Scale bar is 3 µm.

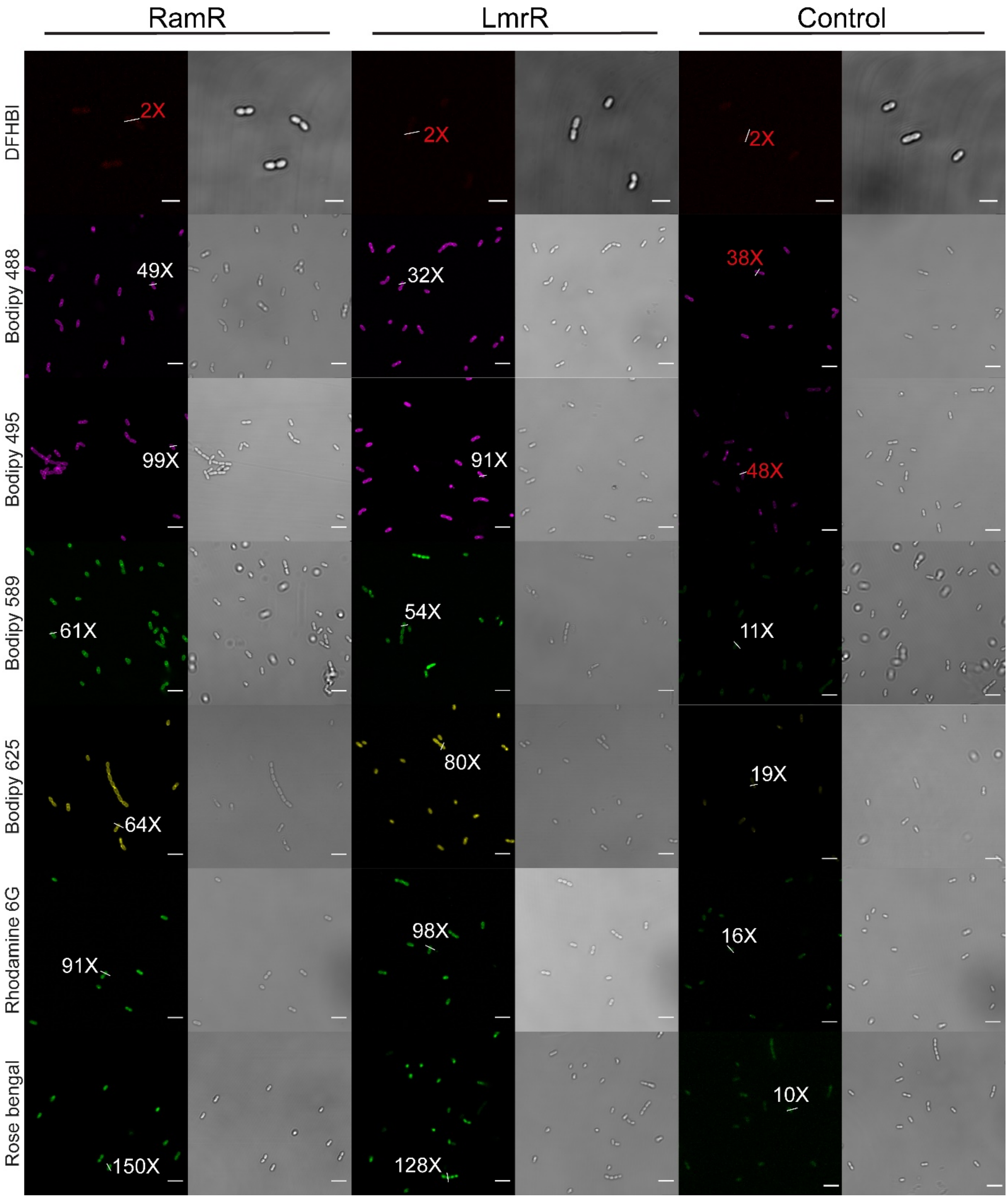


**Supplementary Figure 21**: Overview of labeled *L. lactis* cells expressing cytoplasmic OICP tags from a pNSC8048 (LmrR) and pNZC3GH vector (RamR) respectively. The signal to background fluorescence ratios for arbitrarily picked cells for each dye is mentioned next to white overlays and non-specific staining is depicted as red overlays. All measurements were performed at 30 $^{o}$C. For controls, the ratio was close to 14 ± 3 for all samples indicating some background staining. Scale bar is 3 µm.

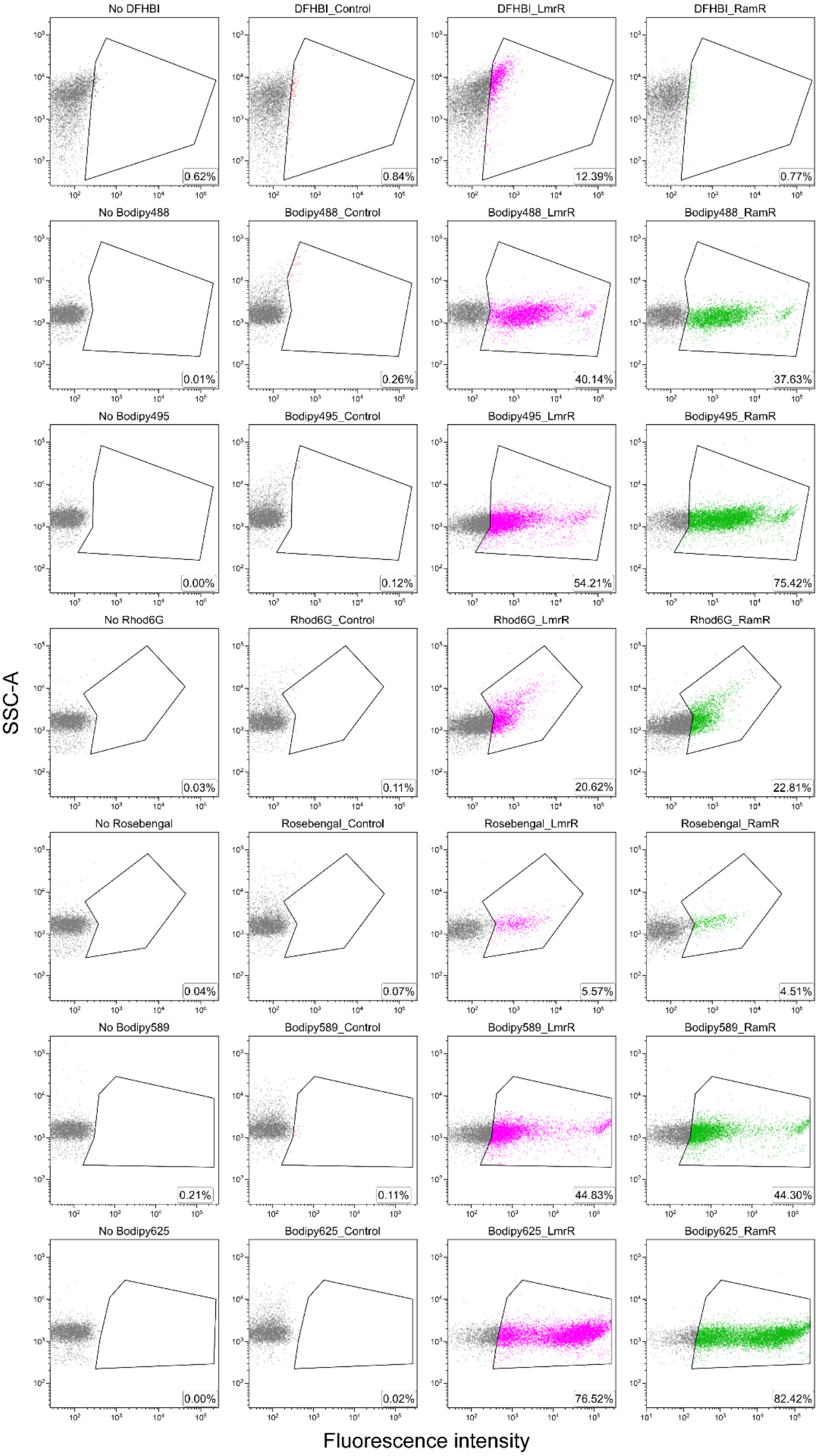


**Supplementary Figure 22**: Flow cytometry side scatter area plots (SSC-A) of *E. coli* cells without any dye (gated black), expressing the cytoplasmic protein OsmY with corresponding dye (control: gated red), expressing LmrR (gated magenta) and expressing RamR (gated green). The low fraction of DFHBI labeled cells with LmrR is likely due to inefficient excitation and collection channels (See methods) as the percentages are significantly lower than estimations of the number of fluorescent cells from microscopy images (**see Supplementary Fig. 20**). The bottom right in each panel shows the % of gated (labeled) cells from a total population of 10,000 cells.

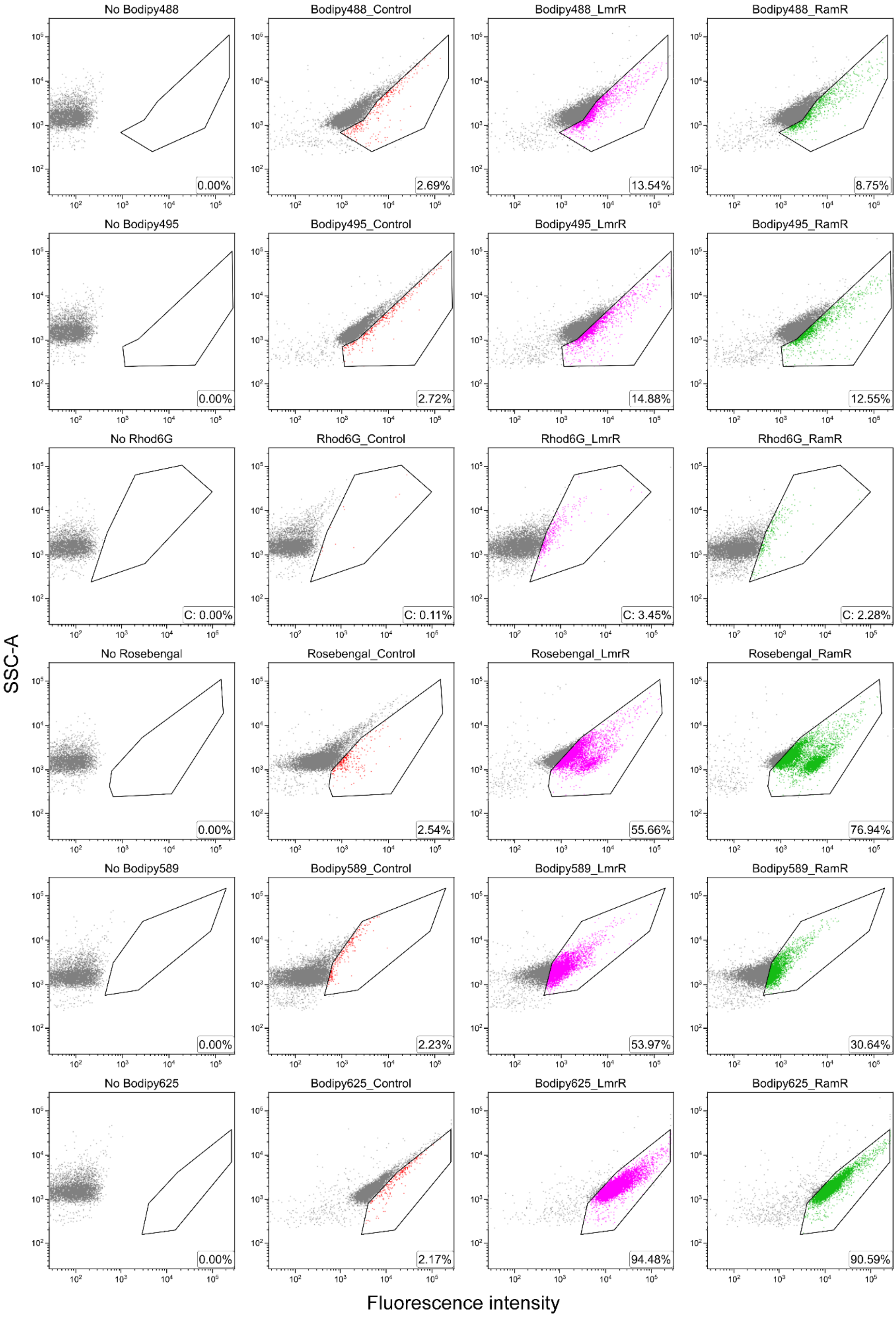


**Supplementary Figure 23:** Flow cytometry side scatter area plots (SSC-A) of Δ*lmrR L. lactis* cells without any dye (gated black), with corresponding dye (control; gated red), expressing LmrR (gated magenta) and RamR (gated green). The bottom right in each panel shows the % of gated (labeled) cells from a total population of 10,000 cells.

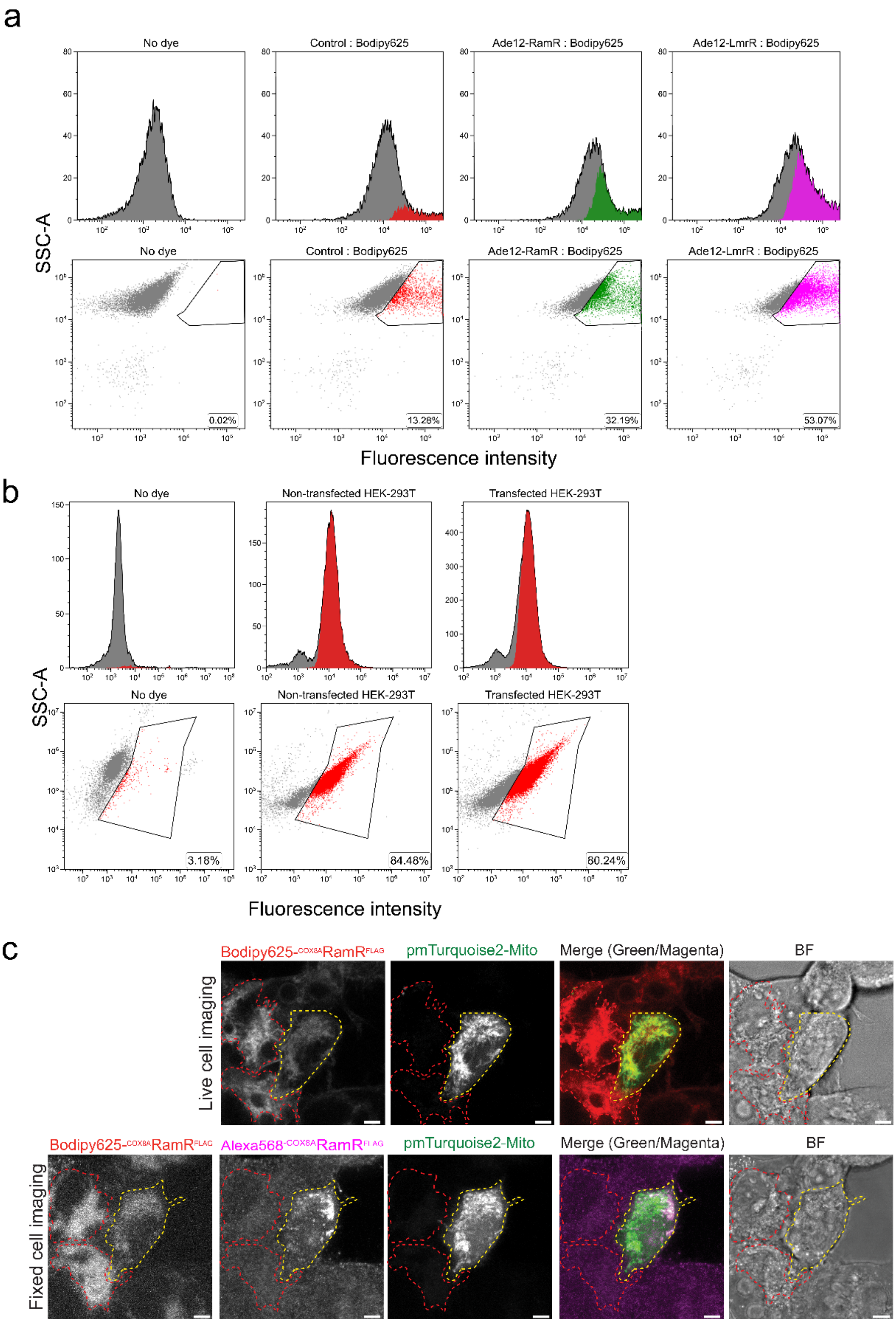


**Supplementary Figure 24:** Flow cytometry side scatter area plots (SSC-A) of *Saccharomyces cerevisiae* (panel a) and HEK-293T cells with Bodipy625 (panel b). Control *S. cerevisiae* (SR80) cells exhibit some background staining that possibly emanates from either non-specific interaction with the cell-wall and/or lipid membrane. *S. cerevisiae* cells expressing Ade-LmrR or Ade-RamR have enhanced fluorescence compared to untransformed *S. cerevisiae* (SR80) cells. FACS analysis of unspecific uptake of Bodipy625 by untransfected HEK-293T cells with no Bodipy625 added and those after the addition of Bodipy625. (b) Untransfected HEK-293T cells exhibit significant uptake (~ 84% labeled cells) of Bodipy625 compared to transfected HEK-293T cells. (c) High background staining was observed in both untransfected (red borders) and transfected (yellow borders) cells.

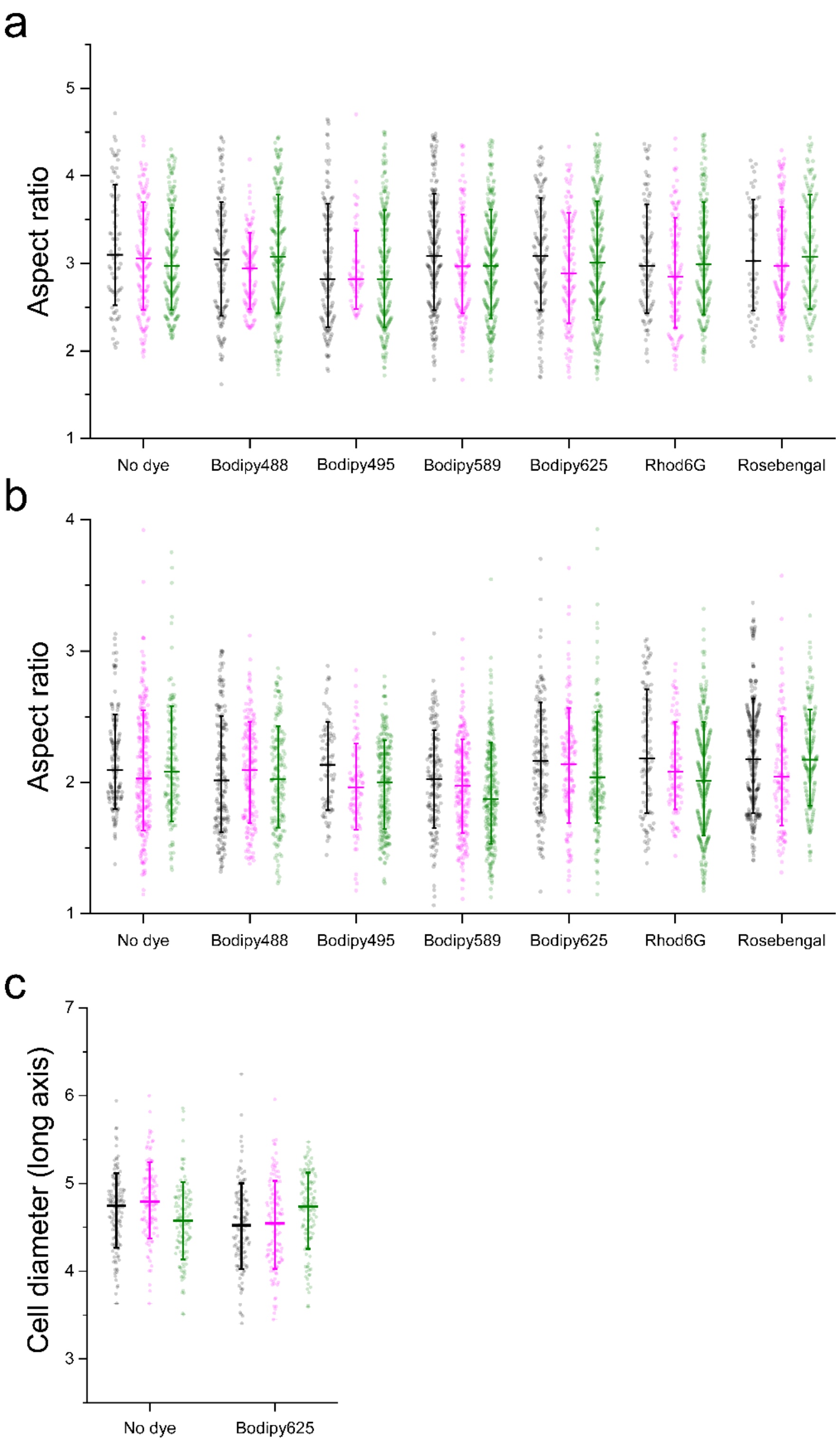


**Supplementary Figure 25**: Effect of dye labeling on OICP tags on cell morphology ascertained by phase contrast microscopy. The aspect ratio (length divided by width) plotted for unlabeled and dye-labeled (a) *E. coli*, (b) *L. lactis* and cell diameter along the long axis for (c) spherical *S. cerevisiae* cells represented in black (control), magenta (LmrR) and green (RamR) as measured from phase-contrast microscopy in the mid-exponential growth phase. The impact of cell diameters for *S. cerevisiae* cells was assessed only with Bodipy625 dye since other dyes did not permeate through the cell-wall. The horizontal bars indicate median values and the vertical bars indicate standard deviation in the sample of at least 100 cells in each case. Images were anlaysed using the MicrobeJ plugin(Ducret et al., 2016) in Fiji software.

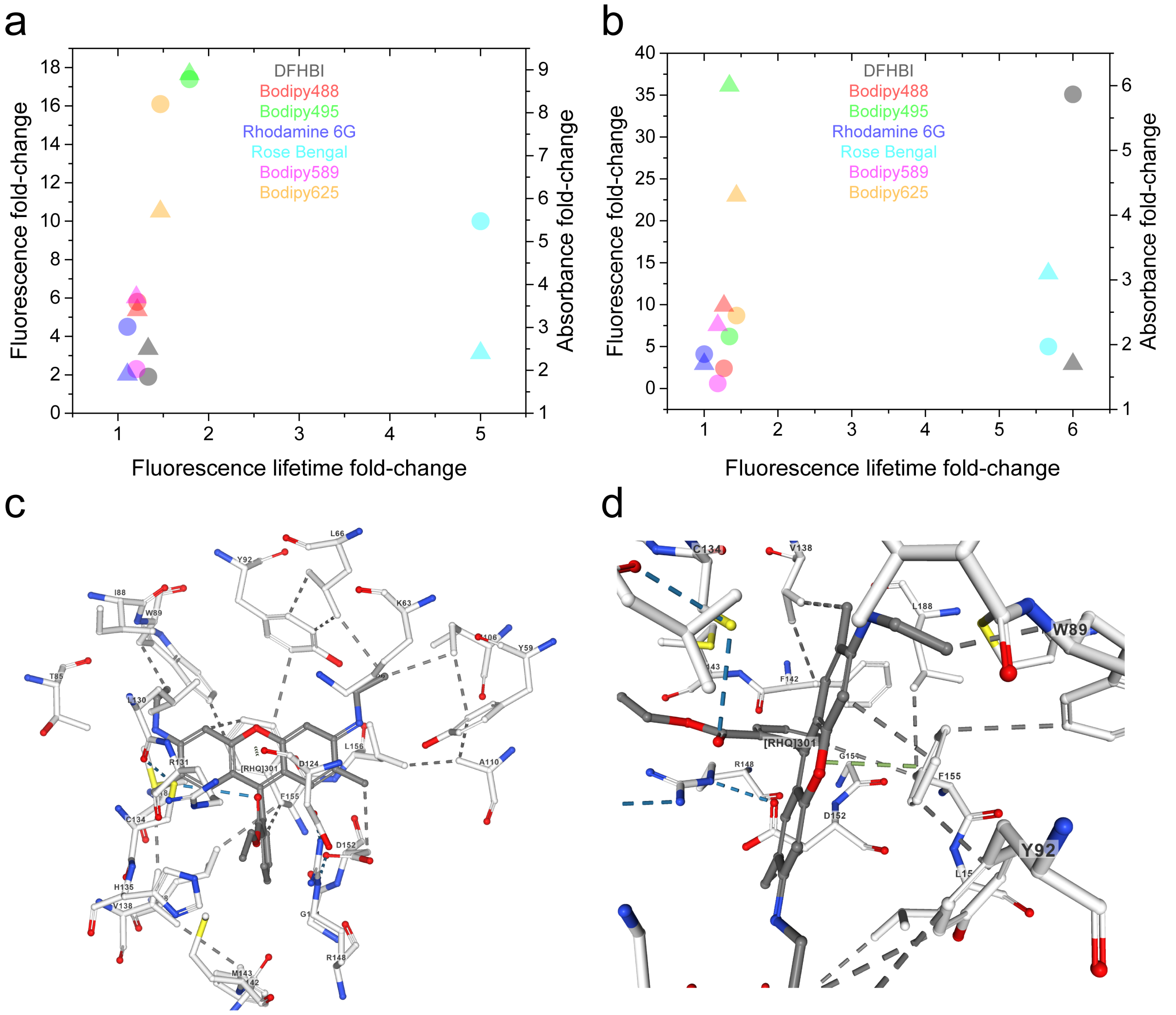


**Supplementary Figure 26**: a-b) Correlation plots of fold-changes in emission fluorescence and absorbance with fold-changes in fluorescence lifetimes for RamR (panel a) and LmrR (panel b). Identical colors depicting fluorescence fold-change (circles) and absorbance fold-changes (squares)indicate the same dye. c-d) An interaction map of a crystal structure of Rhodamine 6G-Bound form of RamR depicting hydrophobic contacts (grey lines) and hydrogen bonds (blue lines). The π-π stacking interactions between Phe155 and Rhodamine 6G are shown in light green (panel b). Interactions are determined by geometric criteria as described in PoseView(Stierand and Rarey, 2010). Image created with Rhodamine 6G-Bound form of RamR PDB ID: 3VVZ on NGL Viewer(Rose et al., 2018).

### Appendix 1: Strains used in the study

| Strain/type | Genotype | Reference |
|---|---|---|
| | ***Escherichia coli*** | |
| BL21(DE3) | fhuA2 [lon] ompT gal (λ DE3) [dcm] ΔhsdS<br>λ DE3 = λ sBamHIo ΔEcoRI-B int::(lacI::PlacUV5::T7 gene1) i21 Δnin5 | (Studier and Moffatt, 1986) |
| BW25113 | Δ(araD-araB)567 Δ(rhaD-rhaB)568 ΔlacZ4787 (::rrnB-3) hsdR514 rph-1 | (Datsenko and Wanner, 2000) |
| | ***Lactococcus lactis*** | |
| NZ9000 | *MG1363; pepN::nisR/K* | (Agustiandari et al., 2008) |
| NZ9000 *ΔlmrR* | MG1363; pepN::nisR/K ; ΔlmrR | (Agustiandari et al., 2008) |
| NZ9000 *ΔlmrR pNSC8048-lmrR* | NZ9000 *ΔlmrR* harboring the plasmid pNSC8048-lmrR | (Agustiandari et al., 2008) |
| NZ9000 *ΔlmrR pNZC3GH-ramR* | NZ9000 *ΔlmrR* harboring the plasmid pNZC3GH-ramR | This study |
| | ***Saccharomyces cerevisiae*** | |
| BY4709 | *MATα ura3Δ0* | (Brachmann et al., 1998) |
| SR80 | BY4709 harboring the empty plasmid pDD-ADH | This study |
| SR80 Ade12-LmrR | BY4709 harboring the plasmid pDD-ADH-Ade12-lmrR | This study |
| SR80 Ade12-RamR | BY4709 harboring the plasmid pDD-ADH-Ade12-ramR | This study |
| | ***Primary human embryonic kidney cells (HEK-293T)*** | |
| WT | Untransfected Wild-type HEK cells | |
| WT RamR | WT cells harboring the plasmid pcDNA3.1-Cox8A-RamR-FLAG | This study |

### Appendix 2: Plasmids used in the study

| Plasmid name | Protein | Promoter | Affinity tag | Inducer | Sub-cellular location | Organism |
|---|---|---|---|---|---|---|
| pET17b-ramR | Cytoplasmic RamR | T7 | Strep-Tag | IPTG | cytoplasm | *Escherichia coli BL21 (DE3)* |
| pET17b-lmrR | Cytoplasmic LmrR | T7 | Strep-Tag | IPTG | cytoplasm | *Escherichia coli BL21 (DE3)* |
| pBAD-cyto-ramR | Cytoplasmic RamR | araBAD | His6-Tag | arabinose | cytoplasm | *Escherichia coli* |
| pBAD-cyto-lmrR | Cytoplasmic LmrR | araBAD | His6-Tag | arabinose | cytoplasm | *Escherichia coli* |
| pBAD-osmY-ramR | osmotically inducible protein Y fused to RamR in the C-terminus | araBAD | - | arabinose | periplasm | *Escherichia coli* |
| pBAD-osmY-lmrR | osmotically inducible protein Y fused to LmrR in the C-terminus | araBAD | His6-Tag; Strep-Tag | arabinose | periplasm | *Escherichia coli* |
| pNM077-ramR-pbP5 | Penicillin-binding protein 5 fused to RamR in the N-terminus | ptrcdown | - | IPTG | inner membrane facing the periplasm | *Escherichia coli* |
| pNM077-lmrR-pbP5 | Penicillin-binding protein 5 fused to LmrR in the N-terminus | ptrcdown | - | IPTG | inner membrane facing the periplasm | *Escherichia coli* |
| - | Control strain | - | - | - | - | *Lactococcus lactis* pNZ9000 *ΔlmrR* |
| pNSC8048-lmrR | Cytoplasmic LmrR | PnisA | - | nisin A | cytoplasm | *Lactococcus lactis* pNZ9000 |
| pNZC3GH-ramR | Cytoplasmic RamR | PnisA | - | nisin A | cytoplasm | *Lactococcus lactis* pNZ9000 *ΔlmrR* |
| pDD-ADH | Prsii426: 2μ empty plasmid (multicopy) with URA3 marker, no protein expressed(Chee and Haase, 2012) | ADH1 | His6-Tag | constitutive | cytoplasm | *Saccharomyces cerevisiae BY4709* |
| pDD-ADH-ade12-lmrR | Cytoplasmic adenylosuccinate synthase fused to LmrR in the C-terminus | ADH1 | His6-Tag | constitutive | cytoplasm | *Saccharomyces cerevisiae BY4709* |
| pDD-ADH-ade12-ramR | Cytoplasmic adenylosuccinate synthase fused to RamR in the C-terminus | ADH1 | His6-Tag | constitutive | cytoplasm | *Saccharomyces cerevisiae BY4709* |
| pcDNA3.1-Cox8A-RamR-FLAG | Codon optimized RamR with a mitochondrial targeting signal (Cox8A) | CMV | FLAG-Tag | constitutive | Inner mitochondrial membrane | *HEK-293T from primary embryonic human kidney* |

### Appendix 3: DNA sequences (5’ to 3’) of constructs used in the study

#### RamR (native)

ATGGTGGCGCGTCCGAAGAGCGAGGACAAGAAACAAGCGCTGCTGGAAGCGGCGACCCAGGC
GATTGCGCAAAGCGGTATTGCGGCGAGCACCGCGGTGATTGCGCGTAACGCGGGTGTTGCGGA
GGGTACCCTGTTCCGTTACTTTGCGACCAAGGACGAACTGATTAACACCCTGTATCTGCACCTGA
AACAGGATCTGAGCCAAAGCATGATCATGGAGCTGGACCGTAGCATTACCGATGCGAAAATGAT
GACCCGTTTCATCTGGAACAGCTACATTAGCTGGGGCCTGAACCATCCGGCGCGTCACCGTGCG
ATCCGTCAGCTGGCGGTTAGCGAGAAGCTGACCAAAGAAACCGAACAACGTGCGGACGATATGT
TCCCGGAACTGCGTGATCTGAGCCACCGTAGCGTGCTGATGGTTTTTATGAGCGACGAGTACCG
TGCGTTCGGTGATGGCCTGTTTCTGGCGCTGGCGGAAACCACCATGGATTTTGCGGCGCGTGAT
CCGGCGCGTGCGGGCGAGTATATTGCGCTGGGCTTTGAAGCGATGTGGCGTGCGCTGACCCGT
GAGGAACAGGCGGCGTGGAGCCACCCGCAATTTGAAAAGTAA

#### RamR (codon-optimized for *L. lactis*)

ATGGGTGGCATGGTGGCGCGTCCGAAGAGCGAGGATAAGAAACAAGCTCTACTTGAAGCTGCTA
CACAAGCCATTGCGCAAAGTGGGATTGCCGCATCGACAGCTGTAATTGCTAGAAATGCCGGTGTT
GCAGAAGGTACCCTTTTCAGATATTTTGCTACCAAAGACGAATTAATTAATACACTGTATTTGCACT
TGAAACAAGATTTATCACAATCTATGATTATGGAATTGGACCGTTCAATTACCGATGCGAAAATGA
TGACAAGATTCATATGGAATTCATACATTAGCTGGGGACTTAACCATCCAGCACGTCATCGTGCAA
TCCGTCAGCTTGCTGTTTCTGAGAAATTAACAAAAGAAACAGAACAACGTGCTGACGATATGTTTC
CAGAACTACGTGATTTATCACATCGTTCAGTCCTTATGGTTTTTATGAGTGACGAGTATCGTGCAT
TCGGAGATGGATTATTTCTGGCACTAGCTGAAACTACTATGGATTTTGCTGCGCGTGATCCCGCG
CGTGCAGGAGAGTATATTGCTCTTGGCTTTGAAGCTATGTGGCGTGCTTTAACTCGTGAGGAACA
GGCGGCGTAA

#### LmrR(K55D/K59Q)

ATGGGTGCCGAAATCCCGAAAGAAATGCTGCGTGCTCAAACCAATGTCATCCTGCTGAATGTCCT
GAAACAAGGCGATAACTATGTGTATGGCATTATCAAACAGGTGAAAGAAGCGAGCAACGGTGAAA
TGGAACTGAATGAAGCCACCCTGTATACGATTTTTGATCGTCTGGAACAGGACGGCATTATCAGC
TCTTACTGGGGTGATGAAAGTCAAGGCGGTCGTCGCAAATATTACCGTCTGACCGAAATCGGCCA
TGAAAACATGCGCCTGGCGTTCGAATCCTGGAGTCGTGTGGACAAAATCATTGAAAATCTGGAAG
CAAACAAAAAATCTGAAGCGATCAAAGCCGCCTGGAGCCACCCGCAGTTCGAAAAA**CATCATCAT**
**CATCATCAT**[His6-Tag]TGA

#### OsmY-RamR

ATGACTATGACAAGACTGAAGATTTCGAAAACTCTGCTGGCTGTAATGTTGACCTCTGCCGTCGC
GACCGGCTCTGCCTACGCGGAAAACAACGCGCAGACTACCAATGAAAGCGCAGGGCAAAAAGTC
GATAGCTCTATGAATAAAGTCGGTAATTTCATGGATGACAGCGCCATCACCGCGAAAGTGAAGGC
GGCCCTGGTGGATCATGACAACATCAAGAGCACCGATATCTCTGTAAAAACCGATCAAAAAGTCG
TGACCCTGAGCGGTTTCGTTGAAAGCCAGGCCCAGGCCGAAGAGGCAGTGAAAGTGGCGAAAG
GCGTTGAAGGGGTGACCTCTGTCAGCGACAAACTGCACGTTCGCGACGCTAAAGAAGGCTCGGT
GAAGGGCTACGCGGGTGACACCGCCACCACCAGTGAAATCAAAGCCAAACTGCTGGCGGACGA
TATCGTCCCTTCCCGTCATGTGAAAGTTGAAACCACCGACGGCGTGGTTCAGCTCTCCGGTACC
GTCGATTCTCAGGCACAAAGTGACCGTGCTGAAAGTATCGCCAAAGCGGTAGATGGTGTGAAAA
GCGTTAAAAATGATCTGAAAACTAAGGGCGGTGGCATGGTGGCGCGTCCGAAGAGCGAGGACAA
GAAACAAGCGCTGCTGGAAGCGGCGACCCAGGCGATTGCGCAAAGCGGTATTGCGGCGAGCAC
CGCGGTGATTGCGCGTAACGCGGGTGTTGCGGAGGGTACCCTGTTCCGTTACTTTGCGACCAAG
GACGAACTGATTAACACCCTGTATCTGCACCTGAAACAGGATCTGAGCCAAAGCATGATCATGGA
GCTGGACCGTAGCATTACCGATGCGAAAATGATGACCCGTTTCATCTGGAACAGCTACATTAGCT
GGGGCCTGAACCATCCGGCGCGTCACCGTGCGATCCGTCAGCTGGCGGTTAGCGAGAAGCTGA
CCAAAGAAACCGAACAACGTGCGGACGATATGTTCCCGGAACTGCGTGATCTGAGCCACCGTAG
CGTGCTGATGGTTTTTATGAGCGACGAGTACCGTGCGTTCGGTGATGGCCTGTTTCTGGCGCTG

GCGGAAACCACCATGGATTTTGCGGCGCGTGATCCGGCGCGTGCGGGCGAGTATATTGCGCTG
GGCTTTGAAGCGATGTGGCGTGCGCTGACCCGTGAGGAACAGGCGGCGTGA

**OsmY-LmrR**

ATGACTATGACAAGACTGAAGATTTCGAAAACTCTGCTGGCTGTAATGTTGACCTCTGCCGTCGC
GACCGGCTCTGCCTACGCGGAAAACAACGCGCAGACTACCAATGAAAGCGCAGGGCAAAAAGTC
GATAGCTCTATGAATAAAGTCGGTAATTTCATGGATGACAGCGCCATCACCGCGAAAGTGAAGGC
GGCCCTGGTGGATCATGACAACATCAAGAGCACCGATATCTCTGTAAAAACCGATCAAAAAGTCG
TGACCCTGAGCGGTTTCGTTGAAAGCCAGGCCCAGGCCGAAGAGGCAGTGAAAGTGGCGAAAG
GCGTTGAAGGGGTGACCTCTGTCAGCGACAAACTGCACGTTCGCGACGCTAAAGAAGGCTCGGT
GAAGGGCTACGCGGGTGACACCGCCACCACCAGTGAAATCAAAGCCAAACTGCTGGCGGACGA
TATCGTCCCTTCCCGTCATGTGAAAGTTGAAACCACCGACGGCGTGGTTCAGCTCTCCGGTACC
GTCGATTCTCAGGCACAAAGTGACCGTGCTGAAAGTATCGCCAAAGCGGTAGATGGTGTGAAAA
GCGTTAAAAATGATCTGAAAACTAAGGGCGGTGGCATGGGTGCCGAAATCCCGAAAGAAATGCT
GCGTGCTCAAACCAATGTCATCCTGCTGAATGTCCTGAAACAAGGCGATAACTATGTGTATGGCA
TTATCAAACAGGTGAAAGAAGCGAGCAACGGTGAAATGGAACTGAATGAAGCCACCCTGTATACG
ATTTTTGATCGTCTGGAACAGGACGGCATTATCAGCTCTTACTGGGGTGATGAAAGTCAAGGCGG
TCGTCGCAAATATTACCGTCTGACCGAAATCGGCCATGAAAACATGCGCCTGGCGTTCGAATCCT
GGAGTCGTGTGGACAAAATCATTGAAAATCTGGAAGCAAACAAAAAATCTGAAGCGATCAAAGCC
GCCTGGAGCCACCCGCAGTTCGAAAAA**CATCATCATCATCATCAT**$^{\text{His6-Tag}}$TGA

**RamR-PBP5 with DsbA signal peptide**

ATGGGCAAAAAGATTTGGCTGGCGCTGGCTGGTTTAGTTTTAGCGTTTAGCGCATCGGCGGCGC
AGTATGAAGATCTCGAGGGTCCGGCTGGTCTGATGGTGGCGCGTCCGAAGAGCGAGGACAAGA
AACAAGCGCTGCTGGAAGCGGCGACCCAGGCGATTGCGCAAAGCGGTATTGCGGCGAGCACCG
CGGTGATTGCGCGTAACGCGGGTGTTGCGGAGGGTACCCTGTTCCGTTACTTTGCGACCAAGGA
CGAACTGATTAACACCCTGTATCTGCACCTGAAACAGGATCTGAGCCAAAGCATGATCATGGAGC
TGGACCGTAGCATTACCGATGCGAAAATGATGACCCGTTTCATCTGGAACAGCTACATTAGCTGG
GGCCTGAACCATCCGGCGCGTCACCGTGCGATCCGTCAGCTGGCGGTTAGCGAGAAGCTGACC
AAAGAAACCGAACAACGTGCGGACGATATGTTCCCGGAACTGCGTGATCTGAGCCACCGTAGCG
TGCTGATGGTTTTTATGAGCGACGAGTACCGTGCGTTCGGTGATGGCCTGTTTCTGGCGCTGGC
GGAAACCACCATGGATTTTGCGGCGCGTGATCCGGCGCGTGCGGGCGAGTATATTGCGCTGGG
CTTTGAAGCGATGTGGCGTGCGCTGACCCGTGAGGAACAGGCGGCGCATCATCATTCCGATGAC
CTGAATATCAAAACTATGATCCCGGGTGTACCGCAGATCGATGCGGAGTCCTACATCCTGATTGA
CTATAACTCCGGCAAAGTGCTCGCCGAACAGAACGCAGATGTCCGCCGCGATCCTGCCAGCCTG
ACCAAAATGATGACCAGTTACGTTATCGGCCAGGCAATGAAAGCCGGTAAATTTAAAGAAACTGA
TTTAGTCACTATCGGCAACGACGCATGGGCCACCGGTAACCCGGTGTTTAAAGGTTCTTCGCTGA
TGTTCCTCAAACCGGGCATGCAGGTTCCGGTTTCTCAGCTGATCCGCGGTATTAACCTGCAATCG
GGTAACGATGCTTGTGTCGCCATGGCCGATTTTGCCGCTGGTAGCCAGGACGCTTTTGTTGGCTT
GATGAACAGCTACGTTAACGCACTGGGCCTGAAAAATACCCACTTCCAGACGGTACATGGTCTGG
ATGCTGATGGTCAGTACAGCTCCGCGCGAGATATGGCGCTGATCGGCCAGGCATTGATCCGTGA
CGTACCGAATGAATACTCGATCTATAAAGAAAAAGAATTTACGTTTAACGGTATTCGCCAGCTGAA
CCGTAACGGCCTGTTATGGGATAACAGCCTGAATGTCGACGGCATCAAAACCGGACACACTGAC
AAAGCAGGTTACAACCTTGTTGCTTCTGCGACTGAAGGCCAGATGCGCTTGATTTCTGCGGTAAT
GGGCGGACGTACTTTTAAAGGCCGTGAAGCCGAAAGTAAAAAACTGCTAACCTGGGGCTTCCGT
TTCTTTGAAACCGTTAACCCACTGAAAGTAGGTAAAGAGTTCGCCTCTGAACCGGTTTGGTTTGGT
GATTCTGATCGCGCTTCGTTAGGGGTTGATAAAGACGTGTACCTGACCATTCCGCGTGGTCGCAT
GAAAGATCTGAAAGCCAGCTATGTGCTGAACAGCAGTGAATTGCATGCGCCGCTGCAAAAGAATC
AGGTCGTCGGAACTATCAACTTCCAGCTTGATGGCAAAACGATCGAGCAACGCCCGCTGGTTGT
GTTGCAAGAAATCCCGGAAGGTAACTTCTTCGGCAAAATCATTGATTACATTAAATTAATGTTCCA
TCACTGGTTTGGTTAA

**LmrR-PBP5 with DsbA signal peptide**

ATGGGCAAAAAGATTTGGCTGGCGCTGGCTGGTTTAGTTTTAGCGTTTAGCGCATCGGCGGCGC
AGTATGAAGATCTCGAGGGTCCGGCTGGTCTGATGGGTGCCGAAATCCCGAAAGAAATGCTGCG
TGCTCAAACCAATGTCATCCTGCTGAATGTCCTGAAACAAGGCGATAACTATGTGTATGGCATTAT
CAAACAGGTGAAAGAAGCGAGCAACGGTGAAATGGAACTGAATGAAGCCACCCTGTATACGATTT
TTGATCGTCTGGAACAGGACGGCATTATCAGCTCTTACTGGGGTGATGAAAGTCAAGGCGGTCG
TCGCAAATATTACCGTCTGACCGAAATCGGCCATGAAAACATGCGCCTGGCGTTCGAATCCTGGA
GTCGTGTGGACAAAATCATTGAAAATCTGGAAGCAAACAAAAAATCTGAAGCGATCAAAGCCGCC
TGGAGCCACCCGCAGTTCGAAAAACATCATCATTCCGATGACCTGAATATCAAAACTATGATCCC
GGGTGTACCGCAGATCGATGCGGAGTCCTACATCCTGATTGACTATAACTCCGGCAAAGTGCTC
GCCGAACAGAACGCAGATGTCCGCCGCGATCCTGCCAGCCTGACCAAAATGATGACCAGTTACG
TTATCGGCCAGGCAATGAAAGCCGGTAAATTTAAAGAAACTGATTTAGTCACTATCGGCAACGAC
GCATGGGCCACCGGTAACCCGGTGTTTAAAGGTTCTTCGCTGATGTTCCTCAAACCGGGCATGC
AGGTTCCGGTTTCTCAGCTGATCCGCGGTATTAACCTGCAATCGGGTAACGATGCTTGTGTCGCC
ATGGCCGATTTTGCCGCTGGTAGCCAGGACGCTTTTGTTGGCTTGATGAACAGCTACGTTAACGC
ACTGGGCCTGAAAAATACCCACTTCCAGACGGTACATGGTCTGGATGCTGATGGTCAGTACAGCT
CCGCGCGAGATATGGCGCTGATCGGCCAGGCATTGATCCGTGACGTACCGAATGAATACTCGAT
CTATAAAGAAAAAGAATTTACGTTTAACGGTATTCGCCAGCTGAACCGTAACGGCCTGTTATGGG
ATAACAGCCTGAATGTCGACGGCATCAAAACCGGACACACTGACAAAGCAGGTTACAACCTTGTT
GCTTCTGCGACTGAAGGCCAGATGCGCTTGATTTCTGCGGTAATGGGCGGACGTACTTTTAAAG
GCCGTGAAGCCGAAAGTAAAAAACTGCTAACCTGGGGCTTCCGTTTCTTTGAAACCGTTAACCCA
CTGAAAGTAGGTAAAGAGTTCGCCTCTGAACCGGTTTGGTTTGGTGATTCTGATCGCGCTTCGTT
AGGGGTTGATAAAGACGTGTACCTGACCATTCCGCGTGGTCGCATGAAAGATCTGAAAGCCAGC
TATGTGCTGAACAGCAGTGAATTGCATGCGCCGCTGCAAAAGAATCAGGTCGTCGGAACTATCAA
CTTCCAGCTTGATGGCAAAACGATCGAGCAACGCCCGCTGGTTGTGTTGCAAGAAATCCCGGAA
GGTAACTTCTTCGGCAAAATCATTGATTACATTAAATTAATGTTCCATCACTGGTTTGGTTAA

**Ade12-RamR-6His**

ATGGTTAACGTTGTATTGGGCTCCCAATGGGGTGATGAGGGTAAAGGTAAACTAGTTGATTTACT
GGTTGGTAAATATGATATTGTAGCCCGTTGCGCCGGTGGTAACAATGCCGGGCATACCATTGTTG
TAGACGGTGTTAAGTATGATTTCCATATGTTACCATCTGGTTTAGTCAACCCAAACTGCCAAAACC
TTTTGGGTAATGGTGTTGTTATTCATGTTCCATCTTTTTTCAAAGAGTTGGAAACCTTGGAAGCTAA
AGGTTTGAAGAACGCAAGGAGTAGATTATTTGTTTCTTCCAGAGCACATTTAGTCTTTGACTTTCA
TCAGGTGACTGACAAGCTAAGAGAATTGGAGTTATCAGGTCGTTCTAAAGATGGTAAAAATATCG
GTACTACCGGTAAAGGTATTGGTCCAACTTATTCCACAAAGGCTTCTAGATCTGGTTTGAGAGTTC
ATCATTTGGTGAATGATCAACCCGGTGCCTGGGAGGAATTTGTTGCTAGATATAAGAGATTATTG
GAAACGAGAAGACAAAGATACGGTGATTTCGAATACGACTTTGAAGCCAAGCTTGCTGAATACAA
GAAGTTAAGAGAGCAACTAAAGCCATTCGTCGTCGATTCTGTCGTTTTCATGCACAATGCTATTGA
AGCAAAGAAAAAGATATTGGTTGAGGGTGCTAATGCTTTGATGTTGGATATTGATTTTGGTACTTA
TCCATATGTGACTTCTTCCAATACTGGTATTGGTGGTGTCCTCACCGGTTTAGGTATTCCTCCACG
TACTATTGATGAAATTTATGGTGTTGTTAAAGCCTACACAACTAGAGTTGGTGAAGGTCCTTTCCC
AACGGAACAATTGAACGAAAATGGAGAAAAACTGCAGACCATTGGTGCTGAATTTGGTGTCACTA
CTGGTCGTAAGCGTCGTTGCGGTTGGTTAGACTTGGTAGTCTTGAAATACTCAACTTTGATCAATG
GATACACGAGTTTGAACATTACCAAGTTAGACGTCCTCGATACTTTCAAAGAAATCCCAGTGGGTA
TTTCATATTCTATTCAAGGTAAGAAGCTAGATTTGTTCCCTGAAGACTTGAATATTCTTGGTAAAGT
TGAAGTTGAATACAAAGTTTTGCCAGGTTGGGATCAAGATATTACCAAAATTACAAAGTATGAAGA
TTTGCCGGAAAACGCAAAGAAGTACTTAAAATATATTGAAGATTTTGTTGGCGTTCCTGTTGAATG
GGTTGGTACCGGCCCCGCAAGAGAAAGCATGTTGCATAAAGAAATTAAAATGGTGGCGCGTCCG
AAGAGCGAGGACAAGAAACAAGCGCTGCTGGAAGCGGCGACCCAGGCGATTGCGCAAAGCGGT
ATTGCGGCGAGCACCGCGGTGATTGCGCGTAACGCGGGTGTTGCGGAGGGTACCCTGTTCCGT
TACTTTGCGACCAAGGACGAACTGATTAACACCCTGTATCTGCACCTGAAACAGGATCTGAGCCA
AAGCATGATCATGGAGCTGGACCGTAGCATTACCGATGCGAAAATGATGACCCGTTTCATCTGGA
ACAGCTACATTAGCTGGGGCCTGAACCATCCGGCGCGTCACCGTGCGATCCGTCAGCTGGCGG
TTAGCGAGAAGCTGACCAAAGAAACCGAACAACGTGCGGACGATATGTTCCCGGAACTGCGTGA
TCTGAGCCACCGTAGCGTGCTGATGGTTTTTATGAGCGACGAGTACCGTGCGTTCGGTGATGGC

CTGTTTCTGGCGCTGGCGGAAACCACCATGGATTTTGCGGCGCGTGATCCGGCGCGTGCGGGC
GAGTATATTGCGCTGGGCTTTGAAGCGATGTGGCGTGCGCTGACCCGTGAGGAACAGGCGGCG
**CATCATCATCATCATCAT**$^{\textbf{His6-Tag}}$TAA

### Ade12-LmrR-6His

ATGGTTAACGTTGTATTGGGCTCCCAATGGGGTGATGAGGGTAAAGGTAAACTAGTTGATTTACT
GGTTGGTAAATATGATATTGTAGCCCGTTGCGCCGGTGGTAACAATGCCGGGCATACCATTGTTG
TAGACGGTGTTAAGTATGATTTCCATATGTTACCATCTGGTTTAGTCAACCCAAACTGCCAAAACC
TTTTGGGTAATGGTGTTGTTATTCATGTTCCATCTTTTTTCAAAGAGTTGGAAACCTTGGAAGCTAA
AGGTTTGAAGAACGCAAGGAGTAGATTATTTGTTTCTTCCAGAGCACATTTAGTCTTTGACTTTCA
TCAGGTGACTGACAAGCTAAGAGAATTGGAGTTATCAGGTCGTTCTAAAGATGGTAAAAATATCG
GTACTACCGGTAAAGGTATTGGTCCAACTTATTCCACAAAGGCTTCTAGATCTGGTTTGAGAGTTC
ATCATTTGGTGAATGATCAACCCGGTGCCTGGGAGGAATTTGTTGCTAGATATAAGAGATTATTG
GAAACGAGAAGACAAAGATACGGTGATTTCGAATACGACTTTGAAGCCAAGCTTGCTGAATACAA
GAAGTTAAGAGAGCAACTAAAGCCATTCGTCGTCGATTCTGTCGTTTTCATGCACAATGCTATTGA
AGCAAAGAAAAAGATATTGGTTGAGGGTGCTAATGCTTTGATGTTGGATATTGATTTTGGTACTTA
TCCATATGTGACTTCTTCCAATACTGGTATTGGTGGTGTCCTCACCGGTTTAGGTATTCCTCCACG
TACTATTGATGAAATTTATGGTGTTGTTAAAGCCTACACAACTAGAGTTGGTGAAGGTCCTTTCCC
AACGGAACAATTGAACGAAAATGGAGAAAAACTGCAGACCATTGGTGCTGAATTTGGTGTCACTA
CTGGTCGTAAGCGTCGTTGCGGTTGGTTAGACTTGGTAGTCTTGAAATACTCAACTTTGATCAATG
GATACACGAGTTTGAACATTACCAAGTTAGACGTCCTCGATACTTTCAAAGAAATCCCAGTGGGTA
TTTCATATTCTATTCAAGGTAAGAAGCTAGATTTGTTCCCTGAAGACTTGAATATTCTTGGTAAAGT
TGAAGTTGAATACAAAGTTTTGCCAGGTTGGGATCAAGATATTACCAAAATTACAAAGTATGAAGA
TTTGCCGGAAAACGCAAAGAAGTACTTAAAATATATTGAAGATTTTGTTGGCGTTCCTGTTGAATG
GGTTGGTACCGGCCCCGCAAGAGAAAGCATGTTGCATAAAGAAATTAAAATGGGTGCCGAAATC
CCGAAAGAAATGCTGCGTGCTCAAACCAATGTCATCCTGCTGAATGTCCTGAAACAAGGCGATAA
CTATGTGTATGGCATTATCAAACAGGTGAAAGAAGCGAGCAACGGTGAAATGGAACTGAATGAAG
CCACCCTGTATACGATTTTTGATCGTCTGGAACAGGACGGCATTATCAGCTCTTACTGGGGTGAT
GAAAGTCAAGGCGGTCGTCGCAAATATTACCGTCTGACCGAAATCGGCCATGAAAACATGCGCC
TGGCGTTCGAATCCTGGAGTCGTGTGGACAAAATCATTGAAAATCTGGAAGCAAACAAAAAATCT
GAAGCGATCAAAGCC**CATCATCATCATCATCAT**$^{\textbf{His6-Tag}}$TAA

### $^{\textbf{Cox8A}}$RamR$^{\textbf{FLAG}}$ with *HindII* and *EcoRI* restriction sites (red)

AAGCTTATGAGCGTGCTGACCCCCCTGCTGCTGCGCGGCCTGACCGGCAGCGCCCGC
CGCCTGCCCGTGCCCCGCGCCAAGATCCACAGCCTGATGGTGGCCCGCCCCAAGAGC
GAGGACAAGAAGCAGGCCCTGCTGGAGGCCGCCACCCAGGCCATCGCCCAGAGCGGC
ATCGCCGCCAGCACCGCCGTGATCGCCCGCAACGCCGGCGTGGCCGAGGGCACCCTG
TTCCGCTACTTCGCCACCAAGGACGAGCTGATCAACACCCTGTACCTGCACCTGAAGCA
GGACCTGAGCCAGAGCATGATCATGGAGCTGGACCGCAGCATCACCGACGCCAAGATG
ATGACCCGCTTCATCTGGAACAGCTACATCAGCTGGGGCCTGAACCACCCCGCCCGCC
ACCGCGCCATCCGCCAGCTGGCCGTGAGCGAGAAGCTGACCAAGGAGACCGAGCAGC
GCGCCGACGACATGTTCCCCGAGCTGCGCGACCTGAGCCACCGCAGCGTGCTGATGG
TGTTCATGAGCGACGAGTACCGCGCCTTCGGCGACGGCCTGTTCCTGGCCCTGGCCGA
GACCACCATGGACTTCGCCGCCCGCGACCCCGCCCGCGCCGGCGAGTACATCGCCCT
GGGCTTCGAGGCCATGTGGCGCGCCCTGACCCGCGAGGAGCAGGCCGCCTGGAGCC
ACCCCGACTACAAGGACGACGACGACAAGTAGGAATTC

### Appendix 4: Gating strategy for flow cytometry studies

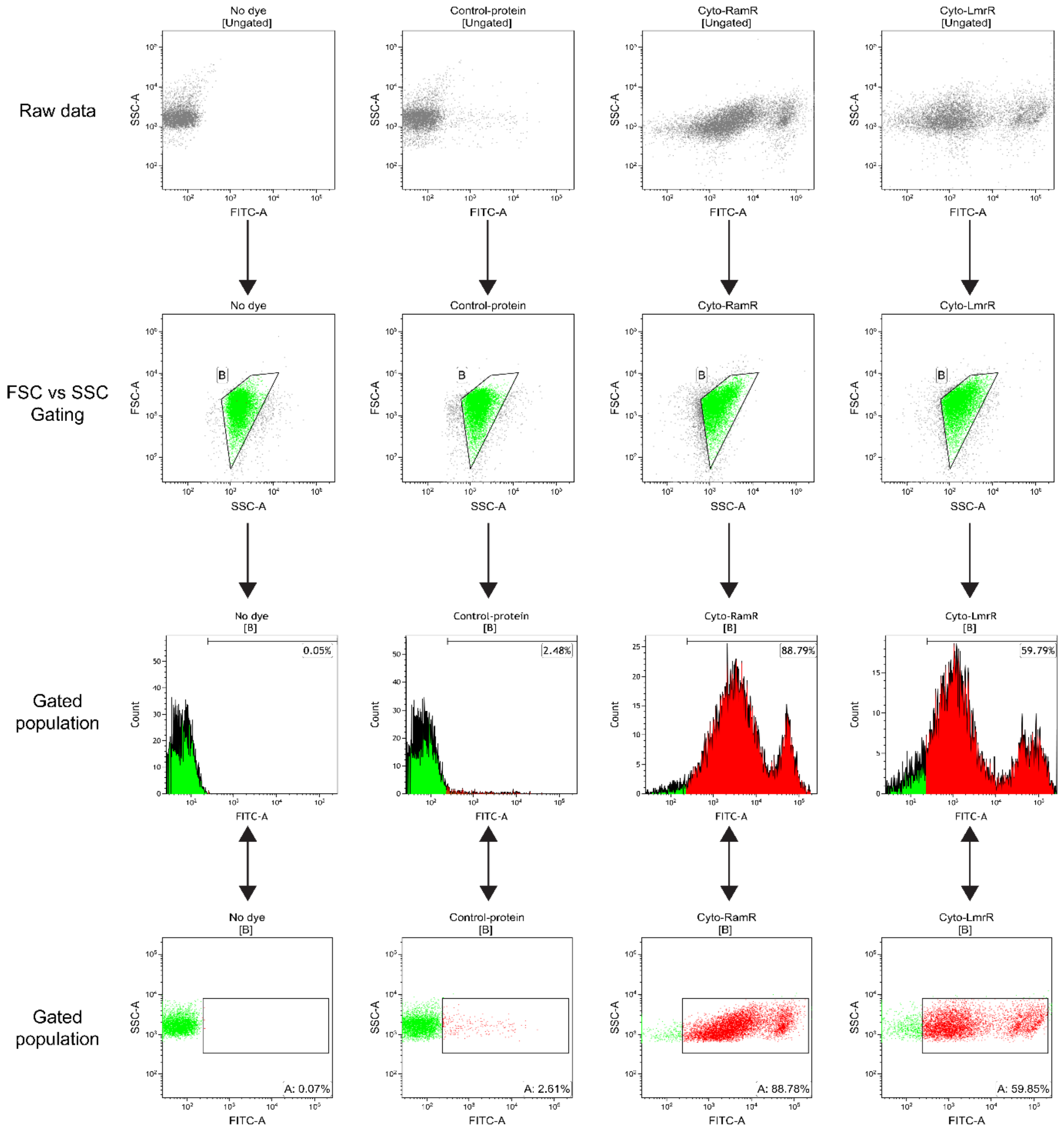


**Gating strategy:** For *E. coli*, *L. lactis,* and *S. cerevisiae* cells, firstly using the FSC/SSC gating, cell debris was removed from the main cell population. A positivity threshold gate for each sample was defined based on unlabeled (0%) and control cells expressing no or control protein yet labeled (< 3%). An identical positivity threshold gate was applied to all samples for a given organic dye. 10000 events were collected for each sample. For HEK293T cells, a starting cell population per sample was collected with the stopping rule of 30000 events per preliminary gate drawn in FSC/SSC. Next, the cells were analyzed placing gates on PE population indicating FLAG expression labeled with Alexa Fluor 568. APC filter was used to detect Bodipy-625 fluorescence. Negative/high background populations were defined by unstained cells visible in PE channel and untransfected but Bodipy-625 pulsed cells in APC channel.
